# Lysosomal Alkalinization Selects for Metabolically Plastic, Motile Cancer Cells under Nutrient Stress

**DOI:** 10.64898/2026.04.16.718649

**Authors:** Leman Nur Nehri, H. Hazal Hüsnügil, Aliye Ezgi Güleç Taşkıran, Hazal Beril Çatalak Yılmaz, Aybar C. Acar, Nalan Liv, Sreeparna Banerjee

**Author notes:** Institute for Chemistry and Bioanalytics, University of Applied Sciences and Arts Northwestern Switzerland, Muttenz, Switzerland (Leman Nur Nehri).

## Abstract

Cancer cells exposed to nutrient deprivation activate adaptive programs to survive metabolic stress, often acquiring enhanced plasticity and motility. We have previously reported that colon cancer cell lines that survived nutrient depletion underwent partial epithelial–mesenchymal transition (pEMT), which was further exacerbated when these cells also underwent lysosomal alkalinization. Here, we have attempted to dissect the molecular mechanisms that drive the motility and shape change from cobblestone to elongated in subpopulations of cells. Using RNA-seq-based bioinformatic analyses integrated with pathway scoring, protein–protein interaction networks, probabilistic modeling and confirmatory experimental data, we have identified the coordinated activation of sublethal apoptotic signaling, fatty acid oxidation, mitochondrial ROS generation, and Ca²⁺-dependent lysosomal exocytosis in the nutrient-depleted cells. Among these phenotypes, the cells undergoing starvation and lysosomal alkalinization exclusively mediated lysosomal exocytosis and cell motility. Probabilistic modeling further revealed non-linear relationships between metabolic stress signals and cell fate transitions, highlighting heterogeneous lysosomal functions as a key determinant of the altered phenotype of cells under nutrient depletion. Overall, our study has identified that aberrant lysosomal functioning in cells under nutrient depletion can specifically select for a subpopulation of cells that are highly viable, metabolically plastic and capable of motility.

## Introduction

Environmental stressors such as limited access to nutrients or oxygen, particularly in rapidly growing solid tumors that are poorly vascularized, selects for a population of cells that can activate adaptive pathways for survival. The same cues may also prompt cancer cells to transition from a proliferative epithelial phenotype to an invasive mesenchymal phenotype ^1^. Tumors are known to rely heavily on the influx of glucose and amino acids (such as glutamine) for survival, which can be used for energy generation, oxidative stress mitigation and protein synthesis. Using a pan-cancer transcriptomics approach, we have shown that fatty acid metabolism is repressed in many tumor types, while genes of the tricarboxylic acid (TCA) cycle and oxidative phosphorylation (OXPHOS) generally remained unaffected. ^2^ Nonetheless, cancer cells deprived of specific nutrients such as glucose or glutamine can rewire their metabolism to utilize alternative sources of energy by modulating the activities of key metabolic enzymes or by the preferential uptake of nutrients from the tumor microenvironment^3^. The available data therefore strongly supports plasticity and adaptation of cells in solid tumors based on the availability of nutrients and metabolites and the expression and/or activity of metabolic enzymes within a complex multicellular ecosystem.

We have previously reported that colorectal cancer cell lines cultured in a nutrient depleted medium containing 10% of the nutrients found in the complete culture medium led to the activation of nutrient sensing pathways and the activation of autophagy, as expected ^4,5^. Additionally, we observed an elongation in the shape of the nutrient deprived cells, along with an increase in the expression of both epithelial and mesenchymal markers, suggesting the activation of partial EMT ^6^. Since nutrient depleted cells have highly active acidic lysosomes for processes such as autophagy ^5^, we treated these cells with the v-ATPase inhibitor Bafilomycin A1 (Baf) with the aim of determining whether alkalinization of the lysosomes could revert the markers of partial EMT. However, to our surprise, we observed that the Baf treated cells acquired an even more elongated shape, expressed higher levels of the markers of partial EMT and were more motile in an *in vivo* chorioallantoic membrane (CAM) assay. ^4^

In the current study, using *in silico*, *in vitro* and *in vivo* tools we aimed to identify the mechanism behind why lysosomal alkalinization in nutrient depleted cancer cells led to an enhanced metastatic phenotype. Using RNA sequencing data (GSE245402) of Caco-2 cells grown in complete medium (nutrient rich, NR), NR + Baf, nutrient depleted (ND) medium and ND + Baf, ^4^ mathematical modeling and experimental validation we have discerned the biological pathways that play a collective role in enhancing the metastatic phenotype in nutrient depleted cells. Our data suggest that inhibition of lysosomal activity in nutrient depleted cancer cells with Baf selects for a group of cells with high viability, motility and metabolic plasticity. The cells survived by activating fatty acid oxidation (FAO) of stored lipids. This in turn generated reactive oxygen species (ROS), which could activate various signaling pathways such as sub-lethal apoptosis. Evaluation of lysosomal localization revealed that lysosomes localized on the long cellular extensions enhanced motility through calcium mediated lysosomal exocytosis. Overall, our data provides a model whereby lysosomal activity defines whether cancer cells under nutrient deprivation will die, or acquire motility to seek nutrients and metastasize.

## Materials and Methods

### Cell culture

Caco-2 cells were purchased from ŞAP Enstitüsü (Ankara, Turkey), RKO and T84 cells were purchased from ATCC. Caco-2 cells were grown in 1 g/L glucose-containing Eagle’s minimum essential medium (EMEM) (VivaCell Biosciences) supplemented with 20% fetal bovine serum (FBS), 1mM sodium pyruvate (VivaCell Biosciences) and 0.1 mM non-essential amino acids (NEAA) (Biological Industries). RKO cells were cultured in EMEM supplemented with 10% FBS, 1mM sodium pyruvate and 0.1 mM NEAA. T84 cells were cultured in DMEM-HAMS F12 (VivaCell Biosciences) medium supplemented with 10% FBS (Capricorn Scientific). All culture media were supplemented with 2 mM L-glutamine (Biological Industries) and 100 U/mL penicillin - 100 μg/mL streptomycin (Biological Industries), and the complete medium was labeled as nutrient rich (NR) medium. The cells were cultured in an incubator with 95% air and 5% CO2 at 37 °C. All cells were tested for mycoplasma contamination frequently and were treated with a maintenance dose of 2.5 µg/mL Plasmocin® (Invivogen). All cell lines underwent STR analysis prior to use.

For nutrient depletion (ND), the cells were incubated for 48h in glucose-and glutamine-free DMEM (VivaCell Biosciences) supplemented with 1% FBS, 0.1 g/L glucose and 0.2 mM L-glutamine, 1 mM sodium pyruvate, 0.1 mM non-essential amino acids and 1% penicillin/streptomycin.

### Treatments

Sub-confluent cells were treated with vehicle or respective drug for 48h in the NR or ND medium. Drug treatments (Table 1) with Bafilomycin A1 (Baf) and Q-VD-OPh (QVD) were started simultaneously with nutrient depletion and lasted for 48h. For the inhibition of lysosomal exocytosis, cells were first treated with the ND medium and 5 nM Baf for 24h, then the respective doses (0.1, 1, 10 μM) of Vacuolin-1 (Vac) were added and the treatments continued for a further 24h. Mitochondrial ROS generation was induced by the addition of antimycin A to the NR cells at the 47th hour of treatment and incubated for 1 h.

**Table 1.** Chemicals used in the study.

| Chemical | Function | Mechanism of Action | Vehicle | Concentration |
| --- | --- | --- | --- | --- |
| **Bafilomycin A1** | Lysosomal acidity inhibitor | Lysosomal V-ATPase inhibition | DMSO (0.1%) | 5 nM |
| **Vacuolin-1** | Lysosomal exocytosis inhibitor | Blocks Ca<sup>2+</sup>-dependent fusion of lysosomes with the plasma membrane | DMSO (0.1%) | 0.1, 1, 10 μM |
| **Q-VD-OPh** | Apoptosis inhibitor | Pan-caspase inhibition | DMSO (0.1%) | 10 μM |
| **Antimycin A** | Mitochondrial ROS generator | Mitochondrial electron transport chain complex III inhibition | Ethanol (0.1%) | 5 μM |

### Cell Proliferation Assay

Cell proliferation was evaluated with an MTT [3(4, 5-dimethylthiazol-2-yl)-2, 5-diphenyltetrazolium bromide] assay according to the manufacturer’s recommendations (Thermo Fisher Scientific, USA). Briefly, 10,000 cells per well were seeded on 96-well plates, with 5 technical replicates. After overnight attachment, cells were treated with vehicle or Baf under NR and ND conditions for 48h. The MTT solution was prepared by dissolving 5mg MTT powder in 1 mL PBS, which then diluted 1:10 with complete growth medium. After treatment period was finished, the medium was discarded and the MTT mixture was added to the wells. The cells were incubated with MTT solution for 4 hours at 37°C, then 1% SDS in 0.01M HCl solution was added to each well. After O/N incubation at 37°C, absorbance was measured at 570 nm with MultiskanGO microplate spectrophotometer (Thermo Fisher Scientific, USA).

### Total Protein Isolation and Western Blot

Cells were harvested with PBS on ice using a cell scraper. The solution was taken into Eppendorf tubes and centrifuged at 2,000 × *g* for 5 minutes at 4°C. The supernatant was discarded, and the cell pellets were washed with phosphate-buffered saline (PBS) and centrifuged again under the same conditions. After washing, the supernatant was removed. Total protein extraction was carried out using M-PER (Mammalian Protein Extraction Reagent, Thermo Fisher Scientific) supplemented with 1X protease inhibitor cocktail (Roche) and 1X phosphatase inhibitor (Roche), according to the manufacturer’s instructions. Cell pellets were resuspended in 30–50 µL of the lysis buffer, vortexed for 10 seconds, and incubated on ice for 10 min, repeated three times in total. The lysates were then centrifuged at 14,000 × *g* for 10 min at 4°C. Protein concentrations were determined using the Bradford Protein Assay.

For western blot, 30 µg of total protein per sample was separated on 12% SDS-PAGE gels that were run at 100V and transferred onto PVDF membranes at 175V for 90 minutes using wet transfer. Next, the membranes were blocked in 5% BSA (AppliChem) in 0.1% TBS-T for 1 h at room temperature (RT) on an orbital shaker. The membrane was incubated with the primary antibody (Table 2) at 4 °C overnight on a shaker. The next day, the membrane was rinsed with 0.1% TBS-T 3 x 10 min and incubated with the secondary antibody at RT for 1 h. Protein bands were visualized with Clarity ECL Substrate (Bio-Rad) by using ChemiDoc MP Imaging System (Bio-Rad). Densitometric analyses were carried out with the ImageLab software. Band intensities for each protein band were measured and the change in expression was determined by normalizing the band intensities to housekeeping protein GAPDH or β-actin, and then to control (NR) treated cells.

**Table 2.** List of antibodies used for western blot.

| **Antibody** | **Size (kDa)** | **Origin** | **Dilution** | **Brand** | **Catalog No** |
| --- | --- | --- | --- | --- | --- |
| **β-actin (C4)** | 45 | Mouse | 1:4000 | Santa Cruz Biotechnology | sc-47778 |
| **PARP (46D11)** | 116,89 | Rabbit | 1:1000 | Cell Signaling Technology | 9532 |
| **PINK1** | 60-66 | Rabbit | 1:1000 | Abcam | ab23707 |
| **ACC(F-2) (S78/80)** | 265 | Mouse | 1:1000 | Santa Cruz Biotechnology | sc-271965 |
| **α-Rabbit** | - | Goat | 1:4000 | Advansta | R05071 500 |
| **α-Mouse** | - | Goat | 1:4000 | Advansta | R05072 500 |

**Table 3.** List of antibodies used for immunofluorescence staining.

| Antibody | Origin | Dilution | Brand | Catalog No |
| --- | --- | --- | --- | --- |
| **LAMP1 (H4A3)** | Mouse | 1:100 | Santa Cruz Biotechnology | sc-20011 |
| **Alexa Fluor®-647 Phalloidin** | - | 1:500 | Cell Signaling Technologies | 8940 |
| **α-Mouse- Alexa Fluor®-488** | Goat | 1:500 | Invitrogen | A11011 |

### Immunofluorescence Assay

T84, RKO and Caco-2 cells (6×10^4^ cells/well) were seeded onto sterile, 12-15mm glass cover slips placed into 12-well plates in complete medium and left for attachment overnight. The following day, the cells were washed with PBS and the relevant treatments were applied. The next day, firstly half-strength fixation was carried out by adding 4% formaldehyde (FA, Sigma-Aldrich) in 0.1M Phosphate Buffer (PB) on top of each well in the same volume as the existing culture medium. The samples were incubated for 15 min at room temperature, and the medium containing the fixation solution was aspirated. Next, 500μL of 4% FA was added to each well for full-strength fixation. The cells were incubated at room temperature for 2h after which the 4% FA was replaced with 1% FA and the plates were stored at 4°C until use. The 4% FA was prepared freshly in PB using a 16% stock solution of FA.

To carry out immunofluorescence staining, firstly the coverslips were removed from the 12-well plate and the fixed cells were washed twice with PBS. Next, the cells were permeabilized with 1% Triton X-100 for 10 min. Following a quenching step (to quench free aldehydes) with 0.15% glycine solution for 10 minutes, the samples were blocked for 10 min in 1% BSA. The cells were incubated with primary LAMP-1 (Santa Cruz Biotechnology) antibody diluted in 1% BSA in PBS and incubated for 1h. Next, the cells were washed with PBS three times and incubated with the Alexa Fluor 488 anti-mouse secondary antibody (Invitrogen) in 1% BSA for 30 min followed by washing. The coverslips were mounted on microscope slides using Duolink® *in Situ* Mounting Medium with DAPI (Sigma-Aldrich) and dried overnight. All steps of IF staining were carried out at RT. The coverslips were visualized with the LSM800 Confocal Laser Scanning Microscope (Zeiss) with 40X and 63X water-based immersion objective. For each field of view, z-stack images were also acquired. Briefly, 5-10 optical sections were collected per image at intervals spanning a total depth of 10–15 µm. The individual slices were subsequently merged using maximum intensity projection in the licensed Zeiss confocal microscope software to generate the final images. The images acquired in the 40X objective were used in the analysis whereas the 63X images were used as representative images.

### Quantification of cell extensions

Images generated from T84, Caco-2 and RKO cells were analyzed for elongations using Fiji (ImageJ). The images were acquired in a LSM800 Confocal Laser Scanning Microscope (Zeiss) with 40X and 63X water-based immersion objective. For this, the longest and shortest axes of individual cells were measured for 25–30 cells per treatment group. The ratio of the longest axis to the shortest axis was calculated for each cell and used as a quantitative measure of cell elongation. These values were imported into GraphPad Prism (version 9.0.0) for bar graph generation and statistical analysis.

### Determination of Lysosomal Positioning

For quantitative spatial analysis of the lysosomes, fluorescence intensity distribution of signals from LAMP1 and F-actin was assessed by line-scan (transect) analysis. A single linear transect was drawn on confocal microscopy images from the center of the nucleus to the cell periphery in NR, ND, ND+Baf and ND+Baf+Vacuolin treated Caco-2 cells. The nuclear center was defined based on the centroid of the DAPI signal, while the cell edge was determined by the termination of the phalloidin signal. Fluorescence intensity profiles for LAMP1 and phalloidin were extracted along each transect. The diameter of the nucleus was defined as 2R; the average diameter of 20 cells from each treatment group were measured. Based on this reference distance, intracellular regions were defined as the perinuclear region (≤1.5R) and the peripheral region (≥3R) ^7^. A total of 80 cells were analyzed, with 20 cells per experimental condition, and normalized fluorescence intensity values were used for comparative analysis between the groups. The average radius (R) and perinuclear region (1.5R) were calculated as 7.7 and 11.5 micron for T84 cells, 4 and 5.7 micron for Caco-2 cells, and 8.1 and 12.1 micron for RKO cells, respectively. Images were acquired with the LSM800 Confocal Laser Scanning Microscope (Zeiss) with 40X and 63X water-based immersion objective.

### Lipid Droplet Analysis (Nile Red Assay)

To image lipid droplets, Caco-2 cells were counted and seeded as 500,000 cells/well onto 6 well plates and left for overnight for attachment. The following day, the medium was removed, the cells were washed with PBS once, then the cells were treated with either vehicle or 5nM Baf under nutrient-rich and nutrient-deficient conditions for 48h. Once treatments were completed, the cell were washed with PBS and fixed with 4% PFA for 15 min at RT. Next, 2.5µg/mL Nile Red in PBS were added onto the wells and incubated for 30 min at RT. The imaging was carried out using FLoid Cell Imaging Station (Thermo Fisher Scientific), excitation was applied with the LED light source (482/18) and fluorescence was captured using the green channel: 532/59 nm. Images were collected under fixed 20X Plan Flourite objective with numerical aperture of 0.45.

### Mitochondrial ROS Assay (MitoSOX)

Mitochondrial superoxide levels were measured using MitoSOX™ Red (Thermo Fisher Scientific) according to established protocols with minor modifications. Cells were cultured under nutrient rich and depleted conditions and treated with vehicle or 5 nM Baf and/or 10 μM Q-VD-OPh (QVD) for 48 h. As a positive control for mitochondrial ROS generation, antimycin A was added to NR cells at the 47th hour of treatment and incubated for 1 h. At the end of the treatment duration, the culture media were removed and the cells were washed once with pre-warmed PBS. Next, the cells were trypsinized and trypsin activity was neutralized with the complete medium. The samples were centrifuged at 500 x *g* for 5 min at 4°C, the supernatant was removed and the cell pellets were washed twice with PBS. MitoSOX Red was prepared as a working solution at a final concentration of 5 µM in PBS. The cells were resuspended in this solution and incubated for 30 minutes at 37°C. Following the incubation, the cells were analyzed with flow cytometer (Acea Novocyte 2060) equipped with a 561-nm laser. MitoSOX™ Red fluorescence was detected using the PI channel. At least 10,000 events were acquired per sample. Cell debris was excluded based on forward scatter (FSC) and side scatter (SSC) parameters, and single cells were gated using FSC-A versus FSC-H. Data were analyzed using FlowJo software. Mitochondrial superoxide levels were quantified as mean fluorescence intensity (MFI) and normalized to the NR medium.

### Chorioallantoic membrane (CAM) assay

The CAM assay was carried out as described previously (Menemenli et al., 2005). Fertilized, specific-pathogen-free Leghorn eggs were obtained from the Tavukçuluk Araştırma Enstitüsü (Ankara, Türkiye) and transported at an ambient temperature of approximately 12°C. Prior to incubation, the eggs were wiped delicately with distilled water and placed in a dedicated incubator maintained at 37 °C and 60–70% relative humidity. After 7 days of incubation, a small hole was created at the blunt end of each egg using sterile tweezers. On embryonic day 8, an observation window of approximately 1 cm in diameter was opened around this site. A drop of sterile PBS was applied to the eggshell membrane, which was gently pinched to facilitate PBS penetration, followed by careful removal of the membrane. The window was subsequently sealed using sterile silk tape.

In parallel, Caco-2 cells were treated with nutrient rich media and deficient media with or without 5nM Baf for 48h. For lysosomal exocytosis inhibition 10μM Vacuolin1 was added after 24h onto nutrient deficient cells treated with 5nM Baf. On embryonic day 9, a cell-media suspension containing 1 × 10⁶ viable cells was mixed at a 1:1 ratio with Matrigel (Corning), and the resulting cell-Matrigel droplets were grafted onto the CAM of the developing embryos. The eggs were returned to the incubator and maintained for an additional 5 days.

On day 14 of embryonic development, the formed ovografts were excised and their dimensions were recorded. A minimum of 10 fertilized eggs were inoculated with tumor cells, and at least ten microtumors per treatment group were collected. Tumor volume was measured and calculated using the formula length × width × height × 0.52. Following microtumor collection, embryos were euthanized immediately by decapitation using sharp scissors.

### Collection of chicken embryonic liver tissues and genomic DNA Isolation

Following decapitation, liver samples were harvested from the sacrificed chick embryos and immediately snap-frozen in liquid nitrogen. The liver was selected as the target organ because it represents the most frequent metastatic site in patients with colorectal cancer ^9^. To avoid cross-contamination of chick cells with human cells, separate sets of instruments were used for the excision of CAM microtumors and for organ collection.

Each collected organ was transferred individually into a Teflon/glass homogenizer (BD) and homogenized into 1 mL genomic DNA isolation buffer (10 mM Tris-HCl, 5 mM EDTA, 200 mM NaCl, 0.2% SDS, pH 8.5) using a pestle. Proteinase K (0.2 ng/mL; New England Biolabs), RNase A (20 ng/mL; New England Biolabs), and sodium acetate (final concentration 0.3 M) were added, followed by incubation at 60 °C for 1 h. Genomic DNA was precipitated by adding 2.5 volumes of ice-cold 99% ethanol and incubating the samples overnight at −20 °C. Samples were subsequently washed twice with 70% ethanol, centrifuged at 10,000 rpm for 10 min, and the supernatant was discarded. DNA pellets were air-dried and resuspended in 200 μL nuclease-free water. The DNA was quantified using a Nanodrop.

According to European Union regulations, the CAM assay is classified as a non-animal experimental model prior to hatching, complies with the 3R principles, and does not require approval from an institutional ethics committee ^10^.

### Detection of disseminated tumor cells by Alu qPCR

The dissemination capacity of Caco-2 cells following nutrient depletion and/or Baf and Vacuolin1 treatment was assessed by evaluating the levels of *Alu* in chick embryonic livers obtained from the CAM assay using qPCR. This approach is based on the fact that Alu repeat sequences are specific to primates; therefore, amplification of Alu from the chick DNA samples indicates the presence of human tumor cells that have disseminated from the ovograft on the CAM into the chick organs.

Human *Alu* sequences were amplified from 200 ng of liver genomic DNA using Alu-specific primers (Table 4) and the SYBR® Green PCR Kit (Promega) according to the manufacturer’s instructions. qPCR was performed using a CFX Connect Real-Time PCR System (Bio-Rad). Ct values obtained with Alu primers were normalized to chicken *GAPDH* Ct values for each sample. Relative human Alu levels, reflecting tumor cell dissemination, were calculated relative to a human genomic DNA control isolated from *in vitro* cultured Caco-2 cells, as described previously. ^4^

**Table 4.**
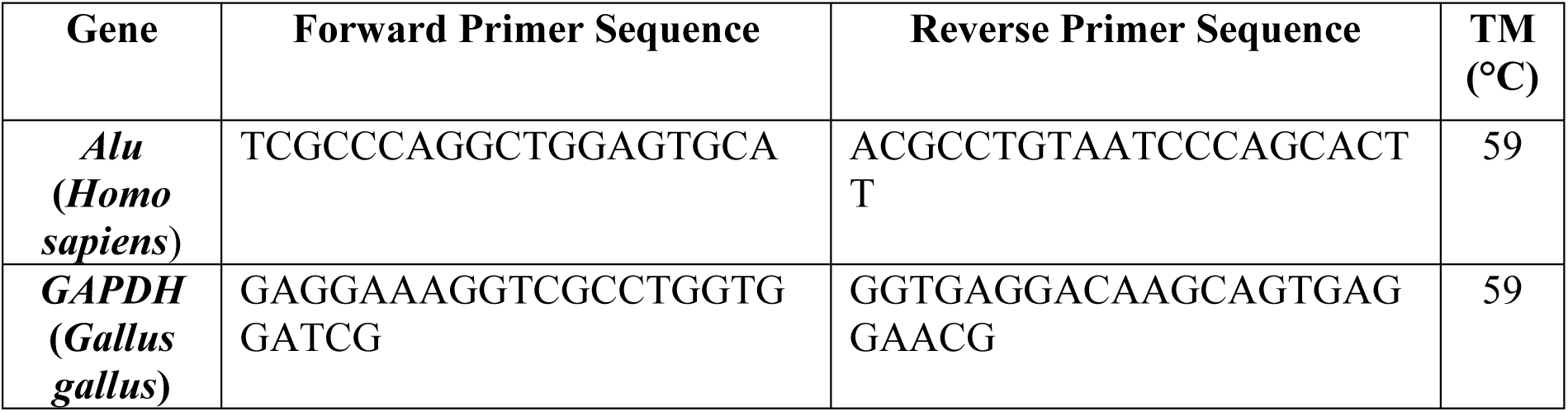
Primers used for Alu qRT-PCR.

For calibration, livers from embryos that had not been inoculated with tumor cells were collected and processed for genomic DNA isolation. To simulate human cell contamination, 200 ng of chicken liver genomic DNA was spiked with 0.01 ng/mL Caco-2 genomic DNA (n = 3). After qPCR analysis, Alu Ct values were normalized to chicken *GAPDH*, and the ΔCt value obtained from 0.01 ng/mL human DNA was defined as 1.0. Relative dissemination levels in liver samples from NR, NR+Baf, ND, and ND+Baf treatment groups were calculated accordingly.

Based on the sensitivity of the Alu qPCR assay, a cut-off value of 0.5 was set for metastasis detection, with relative values >0.5 considered positive for dissemination. All qPCR reactions were performed in technical duplicates for each sample (n ≥ 10). Data are presented as mean ± SEM, and statistical analysis was conducted using the Mann–Whitney U test.

### Bioinformatic Analyses

RNA sequencing data from GSE245402 ^4^ was analyzed. BAM files were obtained for 4 samples (NR, NR+Baf, ND and ND+Baf), with three biological replicates (12 samples in total), aligned and converted to count data using the Galaxy Europe platform (https://usegalaxy.eu/) using featureCount and DESeq2 (Supplementary Methods). All analyses and models were developed using R (RStudio) and Python (PyCharm).

### Comparative Transcriptomic and Protein Analyses

RNA-seq data from GSE245402 were analyzed to assess differential gene expression (DGE) and protein–protein interaction (PPI) dynamics. Differentially expressed genes (DEGs) were identified using thresholds of |log₂FC| > 1 and adjusted p value (p_adj_) < 0.01. Genes were categorized as up- or downregulated and matched against externally curated gene sets related to metabolism, regulated cell death, stress response, and cellular communication (Extended data Table S2). PPI networks of DEGs were analyzed in STRING to identify highly connected, co-expressed proteins, and core protein hubs, and Gene Ontology (GO) enrichment was carried out to determine the associated biological processes. For significant pathways, gene-level scores were calculated using a normalization-based scoring algorithm distinguishing inducers and suppressors (https://github.com/lnnehri/ND-Baf). Comparative analyses between treatment conditions were used to explore whether nutrient depletion and lysosomal alkalinization acted in synergistic or additive manners. Synergy assessment was carried out using logic-based threshold rules adapted from Schrode et al. (2021)^11^ and Trosset and Carbonell (2013)^12^ (https://github.com/lnnehri/ND-Baf).

### Validation of data

To validate our findings, we analyzed independent single-cell RNA seq and proteomics datasets. Differentially expressed apoptosis-related genes from our dataset (NR vs. ND+Baf, |log₂FC| > 1) were compared with metastatic DEGs reported previously. ^13^ Single-cell RNA-seq data (PRJNA611719) from Bernal et al. (2020) ^13^ was processed using RNAStarSolo, followed by Louvain clustering, PCA, and the identification of highly expressed genes. PPI and GO enrichment analyses were carried out on the detected genes and proteins. To validate our findings at the protein level, proteogenomic data from Tanaka et al. ^14^ were re-analyzed by separating primary and metastatic cohorts, filtering for stably expressed genes, and carrying out subsequent network and enrichment analyses (Supplementary Methods)

### Pathway Specific Analyses

DEGs (|log₂FC| > 1, p_adj_ < 0.01) were analyzed across Regulated Cell Death (RCD), apoptosis and calcium-related pathways to explore potential regulatory interactions (Extended data Table S4). Synergy analysis was applied to identify regulatory relationships among highly expressed genes based on parameters derived from previous i*n silico* analyses (Supplementary Methods). PPIs were generated using STRING, and GO enrichment and validation experiments were carried out to identify the associated biological processes (Supplementary Methods). DepMap dependency and expression data were also analyzed to assess the association of the identified gene set with EMT- and apoptosis-related gene expression (Supplementary Methods, https://github.com/lnnehri/ND-Baf).

### Probabilistic Modeling

Parameters obtained from bioinformatics analyses and quantitative experimental data were integrated into probabilistic models to infer state-transition dynamics. The resulting models were designed to generate predictive outcomes that could inform subsequent experimental designs, enabling targeted validation or perturbation of specific transition steps between phenotypic states. The framework described transitions between elongated, rounded, and dead cell states, where rate parameters represented interconversion among live cell phenotypes (elongated and rounded) and loss through death within the ND+Baf Caco-2 cell population. Logistic and exponential models were used to characterize the viability over time of the cells under ND+Baf, ND, and NR+Baf conditions, while Poisson- and Gamma-based frameworks were applied to estimate phenotype probabilities and accumulation-to-threshold behaviors. Categorical distributions were compared using Chi-square tests, and variance homogeneity among phenotypic groups was evaluated using Levene’s test. To integrate bioinformatic, probabilistic, and wet-lab findings into a unified framework, a Bayesian network was constructed by combining bioinformatic and data-driven analyses with probabilistic inference outputs and experimentally validated results to assess causal relationships among key molecular and phenotypic variables under the ND+Baf condition (https://github.com/lnnehri/ND-Baf).

### Statistical Analysis

Each wet lab experiment was carried out with at least 2-3 biological replicates, each having at least two technical replicates. For data analysis, GraphPad Prism v 9 (GraphPad Software Inc.) was used. To evaluate significance, Mann-Whitney U test was used for all *in vivo* experiments and Student’s t test was used for all *in vitro* assays. All statistical analyses were carried out with respect to control (NR) cells, unless indicated otherwise. Statistical significance was defined as a p value of less than 0.05. *p<0.05, **p<0.01, ***p<0.001.

## Results

### Survival and morphology of cancer cell lines to nutrient deprivation and lysosomal alkalinization

The colon cancer cell lines Caco-2, RKO and T84 cells were incubated in the nutrient depleted medium for 24-96 h in the presence or absence of Bafilomycin A1 (Baf, 5nM) and analyzed for cell proliferation using an MTT assay (Extended data Figure S1). Next, the cell proliferation data for Caco-2, T84, and RKO epithelial lines were fitted using both logistic (sigmoidal) and exponential decay models to compare their proliferative capacity. The model performance was compared by residual sum of squares (RSS) and Akaike Information Criterion (AIC) differences (Extended data Table S1). We observed that with nutrient depletion (ND) all three cell lines showed a temporal decrease in proliferation, as expected (Figure 1A). We have previously reported that nutrient depletion led to a modest increase in the mono-phosphorylation of Rb protein in these cell lines, ^4^ suggesting that the cells were arrested at the G1 phase of the cell cycle. Both sigmoidal and exponential decay models performed similarly for Caco-2 and T84, whereas RKO cells exhibited a markedly better fit to the logistic model (ΔAIC = +13.5), indicating that upon reaching a certain duration or threshold of ND, these cells succumbed to cell death (Figure 1A, Extended data Table S1). Upon treatment of the NR cells with Baf, we observed no change in proliferation in Caco-2 and T84 cells, while RKO cells showed a temporal decrease (Figure 1A). The sigmoidal behavior became even more evident in RKO cells (ΔAIC = +38.5), while Caco-2 and T84 maintained exponential kinetics with a slow loss of viability (Extended data Table S1). When Baf was added to the ND cells (ND+Baf), cell death in Caco-2 and T84 remained largely exponential, showing continuous and low-rate decline (ΔAIC ≈ 0). In contrast, RKO cells showed a pronounced transition to sigmoidal kinetics (ΔAIC = +15), suggesting that treatment of the ND cells with Baf amplified a cooperative or threshold-dependent mode of cell death (Figure 1A, Extended data Table S1,). Cell death after the treatment of the NR cells with Baf was observed only in RKO cells (Figure 1A). Treatment of the ND RKO cells with Baf led to a shift from gradual to threshold-driven death dynamics, whereas treatment of ND Caco-2 and T84 cells with Baf caused these cell lines to follow a continuous decay pattern regardless of nutrient availability (Figure 1A, Extended data Table S1). Collectively, these findings highlight distinct death kinetics among epithelial lines subjected to the same nutrient depletion.

**Figure 1.**
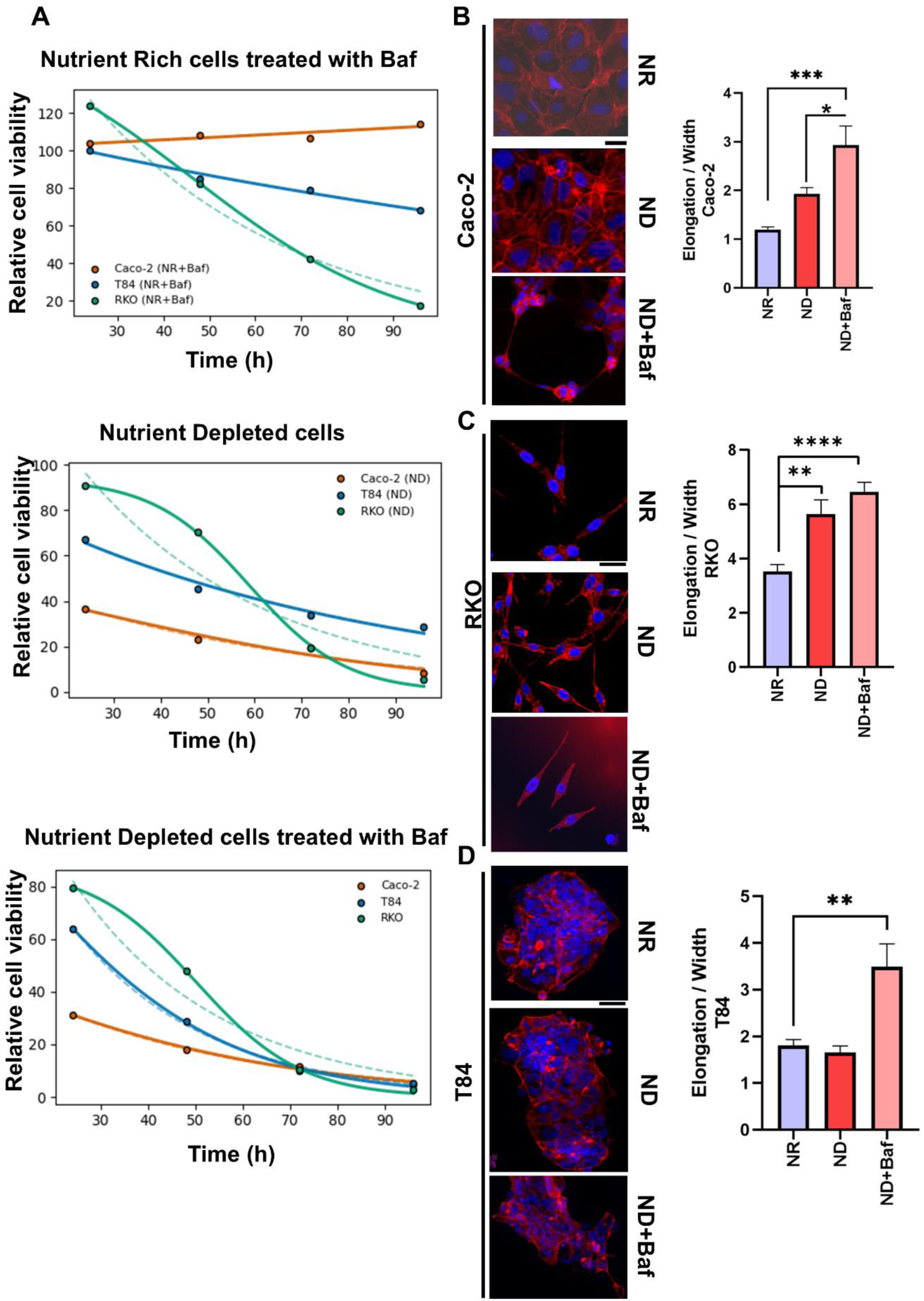
Comparison of cell death kinetics across three colorectal cancer cell lines under treatment conditions. **A.** Time-course viability curves of Caco-2, T84, and RKO cells were modeled using both logistic (solid lines) and exponential (dashed lines) fits at 24, 48, 72, and 96 hours. Panels represent (top) NR+Baf, (middle) ND+Baf, and (bottom) ND conditions. **B.** Phalloidin staining showing a remarkable increase in elongation in the ND+Baf Caco-2 cells (***p<0.001, *p<0.05, ANOVA). **C**. Phalloidin staining in RKO cells showing a significant increase in the elongation of ND cells as well as ND+Baf cells (****p<0.0001, **p<0.01, ANOVA). **D**. Phalloidin staining in T84 cells showing no change in the elongation of ND cells and an increase in the ND+Baf cells (**p<0.01, ANOVA). n=25 for elongation analyses. Scale bar: 50µm.

We have previously reported that several different colon cancer cell lines incubated in the ND medium showed a remarkable elongation in their morphology and expression of partial EMT markers, which could be rapidly reversed when the cells were incubated in complete medium. ^6^ We also observed that when the nutrient depleted Caco-2 cells were treated with the lysosomal alkalinization agent Baf, the cells became even more elongated (Figure 1B) and exhibited even more robust expression of p-EMT markers.^4^ RKO cells also showed an elongated structure when the cells were incubated in the ND medium, which was further exacerbated when the cells were treated with Baf as well (ND+Baf) (Figure 1C). T84 cells, on the other hand, did not show any protrusions with ND and a modest, albeit significant increase in elongation with the ND+Baf medium (Figure 1D).

Based on the kinetic fitting data, Caco-2 cells exhibited a gradual and continuous decline in viability across all tested conditions, with no sharp transition between survival and death (ΔAIC ≈ 0). This contrasted with RKO, which consistently followed a logistic, threshold-dependent death pattern (ΔAIC ≫ 2). Importantly, this death pattern was observed in both NR and ND RKO cells treated with Baf (Figure 1A). The smooth, exponential-like decay observed in the ND and ND+Baf Caco-2 cells indicates that the cell population maintained a progressive, time-resolvable transition rather than an abrupt collapse (Figure 1A, Extended data Table S1).

We next determined the subcellular distribution of lysosomes in the ND+Baf Caco-2 cells, and compared the positioning with T84 and RKO cells versus their controls. For this, we carried out immunocytochemistry using LAMP1 (lysosomal marker) and DAPI (nucleus marker) in the NR, ND, ND+Baf cells. We drew a transect line starting from the middle of nucleus to edge of each cell (n=20 for each group) and a histogram for the signal from LAMP1 and Phalloidin was generated through the transect line (Extended data Figure S2). We observed that nutrient depletion led to the perinuclear localization of lysosomes in T84 cells as also previously shown by us and others. ^5,15,16^ Alkalinization of the lysosomes with Baf led to a shift in the lysosomal distribution towards the periphery. RKO cells had a very similar profile to T84 cells whereby nutrient depletion led to the localization of lysosomes towards the perinuclear region, alkalinization of the ND cells led to some of the lysosomes moving towards the periphery (Extended data Figure S2). Caco-2 cells, on the other hand, did not show any dramatic shift in the population of lysosomes. Nutrient depletion led to a slight shift of the lysosomes towards the periphery, while with the addition of Baf, most of the lysosomes were localized all over the cytosol. This suggested the presence of both peripheral and perinuclear lysosomes. We also stained the cells with siRLysosome, which marks active cathepsin D in cells, and confirmed the near complete loss of lysosomal activity via the loss of siRLysosome puncta in the ND+Baf Caco-2 cells (Extended data Figure S3). Additionally, we used Lysotracker to quantify lysosomal acidity and again observed a complete loss of signal when the ND cells were treated with Baf (Extended data Figure S3).

Caco-2 cells, which showed high survival and remarkable elongation as well as a broader distribution of lysosomes, was further examined for the mechanistic dissection of intermediate or adaptive states under nutrient depletion and lysosomal alkalinization.

### Activation of survival pathways in nutrient deprived Caco-2 cells treated with Baf

We hypothesized a role of lysosomal activity in the survival of the ND+Baf Caco-2 cells. To address this, we first examined the differentially expressed genes (DEGs) from RNA seq data from nutrient rich (NR), NR + Baf, ND, and ND + Baf Caco-2 cells (GSE245402).^4^ DEGs (|logFC2|>1 and p_adj<0.01) (Extended data Figure S4) were analyzed for various PPIs via STRING (Extended data Figure S5) and GO Resource was used to identify the biological processes (Extended data Table S2) in which proteins with dense interaction networks were involved for each comparative treatment conditions. The co-expressed proteins were identified for specific groups of cells (NR, ND, NR+Baf and ND+Baf), and were used for further analysis to compare the differences and similarities in biological properties and pathway-specific genes between these experimental groups. Notably, the ND+Baf cells showed numerous downregulated Extended Data Figure S6) rather than upregulated proteins (Extended Data Figure S7), which exhibited close interactions with each other suggesting that inhibition of biological pathways was more significant in these cells. However, among the upregulated genes, the significant GO terms included regulation of apoptosis, heterotypic cellular interaction signaling, fatty acid metabolism and lipoprotein modeling pathways (Figure 2A). Of note, most of these proteins were upregulated exclusively in the ND+Baf Caco-2 cells (Extended Data Figure S7).

**Figure 2.**
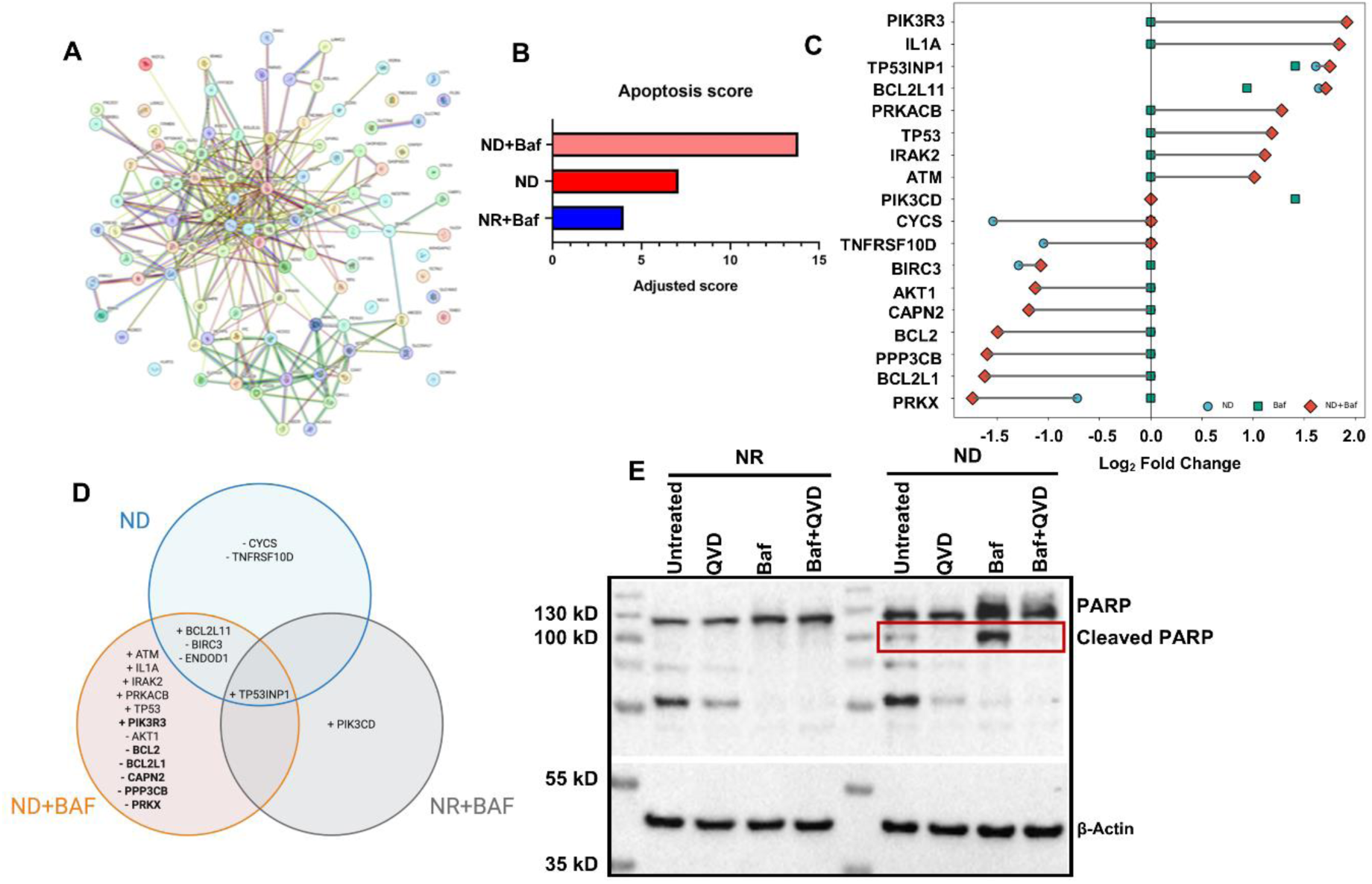
Activation of apoptosis signaling in Caco-2 cells undergoing nutrient depletion and lysosomal alkalinization. **A.** PPI network showing a dense and coordinated regulation between apoptosis, oxidative stress, and lipid metabolic pathways under lysosomal alkalinization and nutrient deprivation (|log₂FC| > 1, p_adj < 0.01). **B.** An apoptosis score showing the highest upregulation of pro-apoptotic genes in the ND+Baf cells. **C.** Dumbbell plot showing log₂ fold changes of apoptosis-related genes under ND (blue circles), NR+Baf (green squares), and ND+Baf (red diamond) conditions. Each horizontal line connects ND and ND+Baf values, illustrating expression shifts between single and combined treatments, while colored points represent individual condition effects, highlighting synergistic effects. **D**. Venn diagram summarizing condition-specific apoptosis-related genes across ND, ND+Baf, and NR+Baf conditions (|logFC2|>1, p.adj<0.01), using NR as control. Gene names shown in bold were uniquely altered in the corresponding condition. Many more apoptosis related genes were differentially regulated in the ND+Baf cells. **E.** PARP cleavage as a marker of apoptosis in the ND+Baf cells. The highest cleavage was observed in the ND+Baf cells, which could be reversed with the pan caspase inhibitor QVD-OPH. Source data for western blots are provided in Extended Data Figure S28.

Since ND+Baf Caco-2 cells underwent extensive cellular detachment, we first focused on regulated cell death (RCD) pathways and observed that the highest number of significant DEGs was associated with apoptosis (Extended Data Table S4). We next generated RCD scores (Extended data Figure S8, S9) to scale the accumulation of RCD signals where a higher score corresponded to stronger signaling. We observed that compared to the NR cells, the ND+Baf cells had the highest apoptosis score, followed by the ND cells and then by the NR+Baf cells (Figure 2B, Extended data Table S5, Extended data Figure S10). None of the other cell death pathways showed a similar trend. This suggests that nutrient depletion combined with decreased acidification of the lysosomes may be responsible for the activation of apoptosis signaling.

To understand whether nutrient depletion and lysosomal alkalinization acted in an additive or synergistic manner to induce apoptosis, we carried out a synergy analysis (see Methods). We observed that many more apoptosis genes were significantly differentially expressed (up or downregulated) in the ND+Baf cells compared to the other experimental groups (Figures 2C, 2D, Extended data Table S6). More importantly, except two, all other genes were up or downregulated in the ND+Baf cells more than the sum of their up or downregulation in ND (nutrient depletion alone) or NR+Baf (lysosomal alkalinization alone) cells (Extended Data Table S6). This suggests that both processes worked in a synergistic manner to induce apoptosis signaling in the ND+Baf cells.

We next examined whether apoptosis signaling could be implicated in the activation of partial EMT in the ND+Baf Caco-2 cells. We observed that the genes upregulated exclusively in the ND+Baf cells formed a close interaction network and were associated with GO terms related to both apoptosis and EMT (Extended data Figure S11). Additionally, DepMap dependency revealed that apoptosis-related genes displayed specific patterns across epithelial and mesenchymal models. Both CRISPR dependency scores (Extended data Figure S12) and transcript abundance (Extended data Figure S13) showed strong linear correlations between EMT-high and EMT-low states, indicating that core dependency on the apoptosis related genes differentially expressed in ND+Baf cells was largely preserved along the epithelial-mesenchymal axis. Notably, several apoptosis-related genes exhibited selective dependency trends, suggesting context-dependent vulnerabilities despite globally conserved apoptotic programs. This pattern is consistent with a partial EMT state, in which epithelial and mesenchymal features coexist without a complete rewiring of apoptotic dependencies.^17^

We also validated the enhancement of apoptosis in metastatic cells from independent datasets. For this, we re-analyzed a publicly available metastatic scRNA-seq data from Bernal et al (2020) (PRJNA611719) (Extended data Figure S14). We observed that the genes formed a connected PPI network that was enriched for apoptosis, mitochondrial electron transport, cytochrome-c release, and regulation of mitochondrial membrane permeability (Extended data Figure S15). GO analysis further highlighted enrichment of both intrinsic and extrinsic apoptotic signaling pathways, together with terms linked to cell migration. These findings may suggest that, despite differences at the individual gene level, apoptosis-related signaling can converge at the pathway level, reflecting a conserved and biologically relevant program in motile cancer cells.

We next validated the induction of apoptosis in the model using PARP cleavage. Indeed, we observed that the ND+Baf cells showed the highest cleavage of PARP compared to the NR, NR+Baf and ND cells, which could be reversed when the cells were incubated with the pan caspase inhibitor QVD-OPH (Figure 2E). Since the RNA seq data was generated from attached (live) Caco-2 cells, we reasoned that the cells show apoptotic signaling, but do not complete apoptosis. A recent study has shown that mouse embryonic fibroblasts (MEFs) treated with the lysosomotropic drug L-leucyl-L-leucine methyl ester (LLOMe) could revert from the initiation of apoptosis via a process that the authors termed as programmed cell revival (PCR).^18^ These cells showed a rounded morphology, organellar dysfunction, caspase activation and PARP cleavage; yet, the floating cells were able to re-attach and were restored to their wildtype phenotype within 30 hours of reattachment.^18^ In our case, the ND+Baf Caco-2 cells also exhibited PARP cleavage. Additionally, when we treated Caco-2 cells with increasing doses of Baf (5 and 100 nM), we also observed a dose dependent detachment of the cells; re-plating of these detached cells could lead to nearly 100% cell survival (data not shown). These data suggest that ND+Baf may also activate apoptosis signaling without actually inducing cell death.

### Activation of lipid oxidation and mitochondrial ROS production

We next evaluated the source of energy for the survival of the nutrient depleted cells. We reasoned that since the nutrient depleted medium contained very limited amounts of glucose and serum, the surviving cells may utilize stored lipids as a source of energy. Indeed, gene set analyses and scoring analyses revealed that lipid metabolism-related pathways, already elevated under ND conditions, became strongly enriched upon ND+Baf treatment (Extended data Figure S6). PPI network results (Extended data Figure S16) and scoring analyses (Extended data Figure S17) suggested a coordinated metabolic adaptation where increased fatty acid oxidation (FAO) was accompanied by elevated ROS activity, reflecting a stress response mechanism that linked metabolic reprograming with survival under nutrient and lysosomal perturbation. We did not observe any difference in the mRNA expression of lipid transporters such as CD36 by qRT-PCR (data not shown) and therefore focused on the utilization of cellular lipids. We observed the highest FAO score in the ND+Baf cells followed by the ND cells and then the NR+Baf cells, using NR cells as the control (Figure 3A). Using a Nile Red assay (Figure 3B), we observed that the NR Caco-2 cells had numerous lipid droplets, which were not significantly affected when the NR cells were treated with Baf. A remarkable decrease in the lipid droplets was seen when the cells were incubated in the ND medium for 48h, suggesting the utilization of lipid stores. When the ND cells were treated with Baf, we observed an increase in the overall signal of Nile Red, although the typical lipid droplet structures were not visible. The diffuse signal from Nile Red may be related to staining of non-polar structures (organelle membranes, free fatty acids) that are localized outside of lipid droplets.^19^ To further substantiate whether the decrease in stored lipid droplets could be attributed to increased fatty acid oxidation (FAO), we evaluated the phosphorylation of Acetyl CoA Carboxylase (ACC) (Figure 3C). We observed that the highest phosphorylation of ACC was observed in the ND and ND+Baf cells, suggesting that the survival of these cells may rely on the energy produced by FAO.

**Figure 3.**
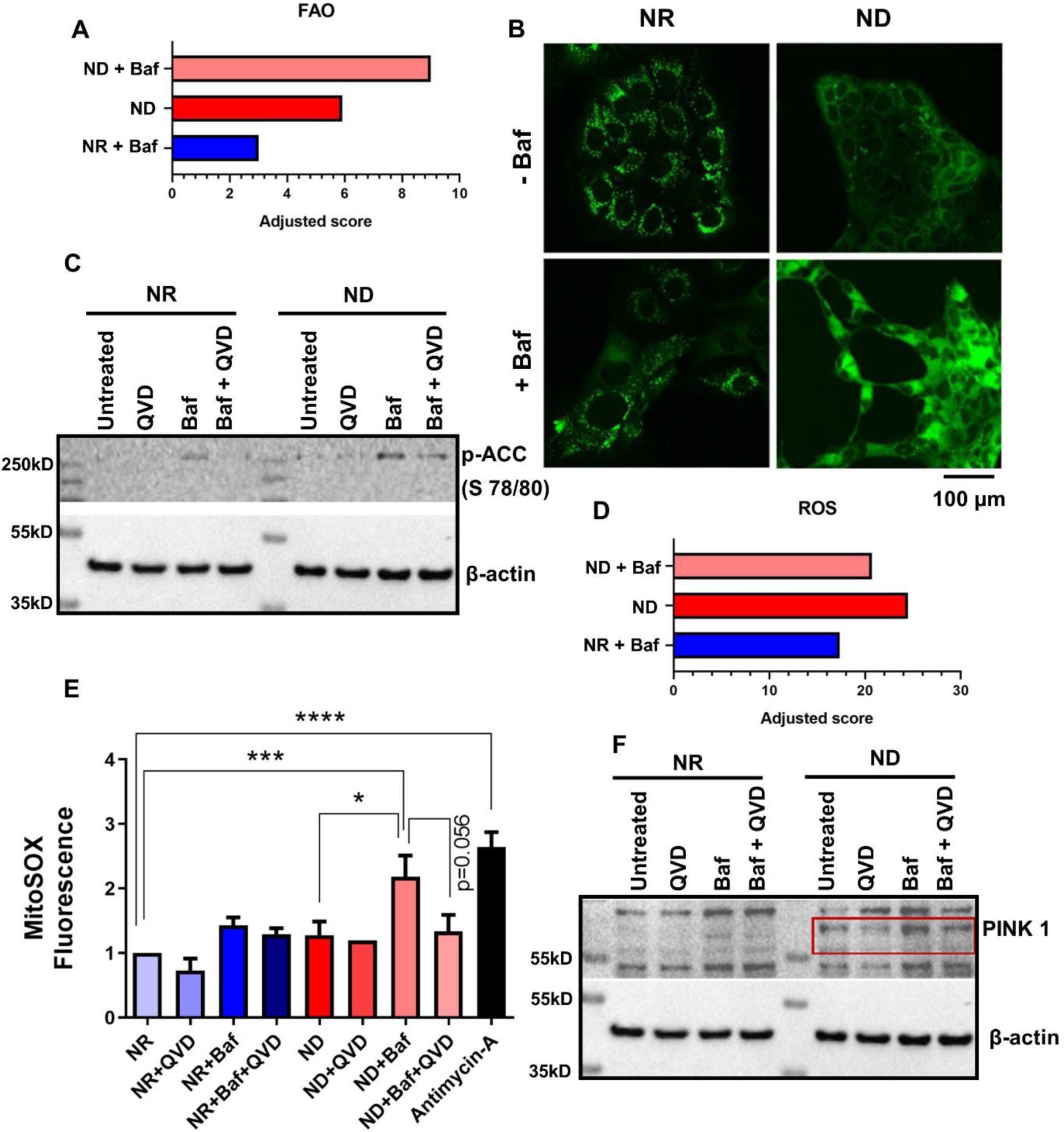
Mitochondrial function defines the survival of nutrient depleted cells undergoing lysosomal alkalinization. **A**. Fatty acid oxidation (FAO) score showing the highest score in ND+Baf cells followed by ND and NR+Baf cells when the NR cells are considered as controls. **B**. Nile Red assay showing the presence of lipid droplets in the NR cells that are decreased in the ND cells and decreased even further in the ND+Baf cells suggesting the utilization of stored lipids when nutrients are scarce. No difference in lipid droplet amount was observed in the NR+Baf cells. **C.** ACC phosphorylation defining the ability of cells to carry out FAO was the highest in the ND+Baf cells. The phosphorylation was reversed when the ND+Baf cells were treated with the pan Caspase inhibitor QVD-OPH. **D**. A ROS score showing the highest score in the ND cells, followed by the ND+Baf cells and then to ND cells. **E**. MitoSOX assay showing the highest mitochondrial ROS levels in the ND+Baf cells. A decrease in the ROS levels was observed when the ND+Baf cells were treated with QVD-OPH. Antimycin A was used as positive control. **F**. Accumulation of PINK1 in the ND and ND+Baf cells suggesting the activation of mitophagy, which was decreased when the cells were treated with QVD-OPH. Source data for western blots are provided in Extended Data Figure S28. The same membrane was used for the western blot shown in Figures 2E, 3C and 3F. Therefore, the same β-actin loading control image is shown.

Since the activation of FAO may also entail the production of mitochondrial ROS, we next evaluated whether ROS production was altered with nutrient depletion and/or lysosomal alkalinization. PPI network analyses under the ND+Baf condition revealed interconnected apoptosis-, FAO-, and mitochondria-related proteins (Extended data Figure S16), while ROS scoring analysis indicated elevated ROS levels in ND cells, whereas the score in the ND+Baf and the NR+Baf cells was slightly lower (Figure 3D, Extended data Figure S17). Moreover, a PPI network indicated that ROS related genes were connected to apoptosis and lipid metabolism-related genes in the ND+Baf cells (Figure 2A). Accordingly, a MitoSOX assay revealed that the mitochondrial ROS levels were significantly increased the ND+Baf cells compared to the ND; however, no such increase was observed when the NR cells compared to NR+Baf cells (Figure 3E). The increase in ROS in the ND+Baf cells may have resulted from increased FAO, suggesting that increased energy generation from mitochondrial metabolism may contribute towards enhanced ROS production.

Integrative PPI analyses revealed that genes associated with apoptosis, mitochondria, and lipid metabolism expressed in ND and ND+Baf conditions formed a close interaction network (Extended data Figure S16). We reasoned that this could be a functional cross-talk to support cellular adaptation and survival during nutrient stress with apoptosis signaling acting as a lynchpin. To support our hypothesis, we treated the NR, ND, NR+Baf and ND+Baf cells with QVD-OPH and observed a decrease in MitoSOX levels when the ND+Baf cells were treated with the apoptosis inhibitor QVD-OPH (p=0.056).

An additional characteristic of cells that survive despite low availability of nutrients is the activation of mitophagy. Moreover, ROS signaling is known to activate mitophagy, often to decrease the levels of ROS by degrading mitochondria. ^20^ To support our finding, we evaluated the levels of PINK1. This protein is normally localized on the outer mitochondrial membrane and is cleaved and inactivated in cells with healthy mitochondria. A decrease in the mitochondrial membrane potential either through mitochondrial stress or mitophagy abrogates the destruction of PINK1 and allows the protein to accumulate on the outer mitochondrial membrane ^21^. We observed that the ND+Baf cells had the highest level of PINK1, followed by the ND cells suggesting the initiation of mitophagy in these cells. The NR cells did not show any PINK1 accumulation suggesting that the mitochondria were functional in these cells (Figure 3F). Once again, we observed that treatment of the ND+Baf (but not NR+Baf) cells with the apoptosis inhibitor QVD-OPH could decrease (albeit modestly) the levels of PINK1, suggesting a crosstalk between apoptosis and mitophagy pathways (Figure 3F).

Overall, our data suggests that ND+Baf cells activate an intricate mechanism of lysosome-mitochondria crosstalk to increase lipid oxidation, ROS generation and apoptosis signaling in order to obtain enough energy for survival.

### Activation of pathways for cellular motility in nutrient depleted Caco-2 cells treated with Baf

We next evaluated the mechanism behind the high motility of the ND+Baf cells. We reasoned that the positioning of the lysosomes could be implicated in cellular motility. Thus, the lysosomes positioned closer to the extensions of the plasma membrane in the ND+Baf cells were more likely to undergo Ca²⁺-mediated exocytosis, ^15^ thereby contributing to extracellular matrix modulation and enhanced cell motility. This hypothesis was supported by our previous finding that the lysosomal Ca^2+^ channel TRPML1/MCOLN1 was highly expressed in the ND+Baf cells, followed by the ND and the NR+Baf cells, with the lowest expression in the NR cells.^4^

Revisiting the RNA seq data generated from these cells, we observed that the combination of both nutrient depletion and lysosomal alkalinization was necessary for processes such as ROS generation, lipid metabolism and apoptosis signaling (Extended data Figure S7). However, Ca^2+^ related regulatory pathways were enriched (|logFC2|>1 and p_adj<0.01) exclusively in the ND+Baf cells compared to the ND cells (Extended data Figure S18, S19). Among the Ca^2+^ related GO terms that were significantly upregulated were Ca^2+^ ion-dependent exocytosis, SNARE complex assembly, and protein secretion, among others. Of note, none of these genes showed any significant differential regulation between NR and NR+Baf cells. Additionally, the exocytosis-related genes formed a tightly connected interaction network in STRING, and several proteins within the ND+Baf exocytosis gene set were directly associated with Ca²⁺-dependent regulatory processes (Extended data Figure S20). This was also corroborated by the remarkable increase in the expression of the Ca^2+^ channel TRPML1 in the ND+Baf cells. ^4^ Further analysis of the Ca^2+^ mediated pathways indicated a downregulation of genes related to cell-cell communication and adhesion in the ND+Baf cells (Extended data Figure S21).

Given that Ca²⁺-mediated cellular processes can influence cytoskeletal remodeling and cellular shape, and that our bioinformatic analyses highlighted calcium signaling and FAO as key pathways that were enriched in the ND+Baf treated cells, we next incorporated these parameters into a probabilistic framework to better evaluate the population dynamics of Caco-2 cells under starvation and lysosomal alkalinization. In this model, we linked the morphological states of the cells (elongated/metastatic, rounded/dormant, and dead) with lipid and calcium dynamics (Extended data Figure S22). Using Poisson-based probability distributions, we estimated the likelihood of each morphological state after 48 hours, showing that a maximum of ∼15% of the living cells acquired an elongated morphology, which closely fitted the Poisson model and was consistent with the imaging data (Extended data Figure S23). These probabilities were then used to parameterize Gamma distributions that connected molecular parameters, in this case, FAO and intracellular calcium release identified from prior bioinformatic pathway analyses, to phenotypic outcomes that included rounded, death or elongated cell shape.

Categorical comparison of the Gamma-modeled distributions using Chi-square (Extended data Figure S24, S25) and Levene’s tests (Figure 4A) revealed significant differences in the variances between the phenotypes, with particularly high molecular heterogeneity in the elongated cells compared to rounded and dead populations. Chi-square analysis further confirmed that all three phenotypes deviated significantly from the expected distributions (elongated: X² = 101.15, df = 4, p < 2.2×10⁻¹⁶; rounded: X² = 32.85, df = 4, p = 1.28×10⁻⁶; dead: X² = 19.10, df = 4, p = 7.53×10⁻⁴), with the strongest deviation observed in elongated cells, indicating the presence of molecular heterogeneity, very likely mediated via calcium signaling within this phenotype. Of note, the model assumes baseline viability of the ND+Baf cells as a prerequisite, presumably by getting energy from FAO. Moreover, apoptotic signaling may be activated within this viable population without inducing cell death. We reasoned that only those ND+Baf cells capable of Ca²⁺ release from the lysosomes followed by lysosomal exocytosis, would transition to an elongated phenotype.

**Figure 4.**
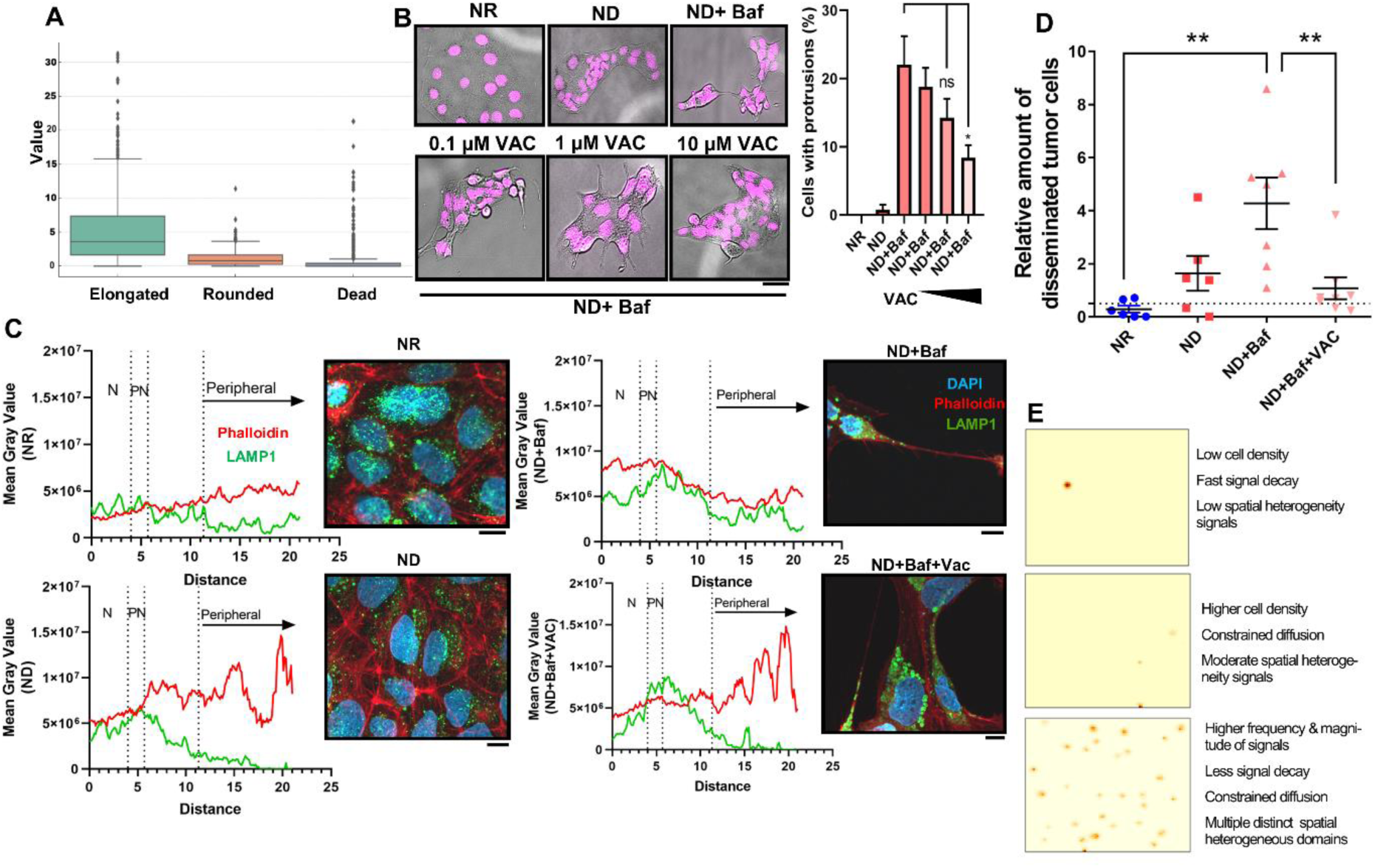
The role of Calcium in the shape and motility of ND+Baf cells. **A.** Levene’s test indicated that the elongated cells had the highest variation which suggests a stronger role of effectors such as calcium mediated signaling and fatty acid oxidation in these cells compared to the Rounded (Live) or Dead cells**. B.** Inhibition of lysosomal exocytosis with increasing doses of Vacuolin (Vac, 0.1, 1 and 10µM) led to a decrease in the number of protrusions in the ND+Baf cells in a dose dependent manner. **C.** The localization of lysosomes (marked by LAMP1) was scattered throughout the cytoplasm in the NR cells, became more perinuclear in the ND cells. The ND+Baf cells had more lysosomes towards the periphery, primarily at the projections. The ND+Baf+Vac cells showed a perinuclear shift of the lysosomes **D.** CAM assay showing significantly higher motility of *Alu* positive cancer cells from the ovograft to the liver of the chicken in the ND+Baf cells. Treatment of the ND+Baf cells with Vac led to a decrease in cellular motility. **E**. Modeling of spatial heterogeneity in based on local cellular interactions. Upper panel: Single localized heterogeneity hotspot produced under low signal generation conditions. Middle Panel: Sparse, weakly distributed heterogeneity signals in a low-density cell configuration. Lower panel: Spatial heterogeneity emerges through the coordinated modulation of three interdependent processes that jointly determine the frequency, strength, and spatial confinement of lysosomal exocytosis–associated signals.

Based on our modeling assumptions, and given that Ca²⁺ release is known to be a critical determinant of lysosomal exocytosis, ^22^ we treated the ND+Baf cells with increasing concentrations (0.1, 1 and 10µM) of the lysosomal exocytosis inhibitor vacuolin-1 (Vac). Vac was shown to specifically inhibit the Ca^2+^-dependent fusion of lysosomes with the plasma membrane and the release of lysosomal content. ^23^ We observed that the elongated structure and protrusions of the ND+Baf Caco-2 cells could be reversed with Vac in a dose dependent manner (Figure 4B). One of the markers of lysosomal exocytosis is the presence of lysosomal proteins such as LAMP1 on the plasma membrane.^24^ For this, we carried out immunocytochemistry using LAMP1, Phalloidin and DAPI in the NR, ND, ND+Baf and ND+Baf cells treated with 1 and 10µM Vac (ND+Baf+Vac) (Figure 4C). We observed considerably more lysosomes at the protrusions of the ND+Baf cells, compared to the ND or NR cells. The localization of the lysosomes (LAMP1 signal) became more perinuclear in the ND+Baf+Vac cells compared to the ND+Baf cells (Figure 4C). Ca^2+^ dependent lysosomal exocytosis in cancer cells was shown to modulate the ECM ^25^; therefore, we reasoned that inhibition of the lysosomal fusion with the plasma membrane may reverse the increased in cellular motility observed in the ND+Baf cells. To substantiate this, we carried out an *in vivo* chorioallantoic membrane (CAM) assay whereby we evaluated the presence of human cells in the liver of the chicken by carrying out PCR for the mammalian marker *Alu*. We observed that while ND+Baf cells had the highest cellular motility, the ND+Baf cells treated with Vac showed significantly decreased motility, to the level observed with the NR cells (Figure 4D).

We next carried out a stochastic, grid-based agent-based model that was designed to investigate how local lysosomal release dynamics could generate spatial heterogeneity and directional behavior at the population scale (Supplementary Methods). Rather than assuming uniform cellular responses, this model simulated probabilistic release events that occurred preferentially at sites of peripheral activation and cell-cell contact. In the model, each lysosomal calcium release event contributes to a local heterogeneity signal that accumulates on a two-dimensional grid and is modulated by four key processes: the likelihood of release initiation, the magnitude of each release event, the persistence of the signal over time, and its spatial diffusion. By jointly tuning these parameters, the system transitions from isolated, rare signal events to multiple, reinforced heterogeneity hotspots. Increased probabilities of lysosomal alkalinization and contact-dependent activation led to more frequent release events, while stronger per-event contributions amplified local signal intensity. Concurrently, slower signal decay and reduced diffusion prevented global homogenization, allowing localized signal accumulation (Figure 4E). Biologically, this modeling framework captures a scenario of lysosomal exocytosis as a cumulative and spatially organized process rather than a uniform cellular response. It supports our experimental finding of a select group of cells that show the elongated shape and the presence of lysosome at the membrane protrusions; it is likely that these very cells also contribute towards enhanced metastasis. Our data also supports the presence of cellular heterogeneity whereby starvation and lysosomal alkalinization specifically selects for viable cells that are either elongated or rounded.

Regions exposed to repeated peripheral stress and intercellular contact become enriched in persistent signals, forming discrete microenvironments that differ from surrounding cell groups. Spatial heterogeneity in the model emerges through the coordinated modulation of three processes: (i) Increasing the baseline and contact-dependent probabilities of lysosomal release elevates the frequency of local signal generation, particularly at sites of peripheral activation and cell-cell contact. (ii) Enhancing the contribution of each individual release event while decreasing signal decay enables repeated release events at the same spatial location to accumulate over time, resulting in progressively stronger local signals. (iii) Constraining spatial diffusion limits signal spread, preventing global homogenization and promoting local signal retention.

In the first simulation regime (Figure 4E, upper panel), signal generation probabilities and cell density were kept low, and signal decay was relatively fast. Under these conditions, heterogeneity signals were rarely produced and remained spatially isolated. Consequently, a single, localized hotspot emerged on the grid, representing rare and stochastic cellular events with minimal collective organization. In the second regime (Figure 4E, middle panel), cell density and contact-dependent signal generation probabilities were increased, while spatial diffusion was moderately constrained. This parameter combination led to the emergence of multiple, but still limited, heterogeneity hotspots. These hotspots were spatially separated and less intense, indicating a transition from isolated events toward regionally clustered, interaction-driven heterogeneity. In the third regime (Figure 4E, lower panel), both the frequency and magnitude of signal-generating events were further increased, signal decay was reduced, and diffusion was more strongly restricted. This allowed signals to persist longer and remain localized near their points of origin. As a result, multiple distinct and pronounced hotspots formed across the grid. This regime captured a scenario in which collective cellular behavior and local positive feedback gave rise to robust, spatially heterogeneous domains. Thus, various mechanisms work together to drive the transition of a group of cells, selected with nutrient depletion and lysosomal alkalinization, from rare and isolated events to persistent, spatially confined hotspots where Ca²⁺-dependent lysosomal exocytosis takes place.

We next evaluated the temporal dynamics of the transition to the elongated phenotype as a function of calcium signaling and FAO. Given an average population lifespan of 48 hours, our analysis indicates that the commitment to shape transition from rounded to elongated must occur within roughly the first quarter of the lifespan before a decline of the cell population (Extended data Figure S26). A cumulative probability distribution was applied within the modeling framework and the model predicted that cells reached the elongation threshold (probability = 0.894) in their early life-time (Extended data Figure S27).

### Combined modeling of the acquisition of survival and motility in the ND+Baf cells

We propose in the current study that nutrient depletion, when also accompanied by dysfunctional lysosomes, can select for a subset of cells that maintain viability through FAO, and increased cell motility via calcium mediated exocytosis. We next integrated these findings into a minimal Bayesian network to simulate and explain the verified-coordinated cellular states observed under ND+Baf conditions. Based on our distribution models (Figure 4A, Extended data Figures S22-S27) we used a vectoral model to explain how cancer cells acquire motility when they are in an unfavorable environment (Figure 5A). Consistent with probabilistic predictions (Extended data Figure S28), we validated that the alkalinized lysosomes preferentially accumulated the cellular extensions, where they facilitated Ca²⁺-dependent exocytosis and promoted directional migration (Figure 4). To model these experimentally validated parameters and explore their potential causal relationships, we constructed a Bayesian network framework (Figures 5B, 5C). The model results revealed condition-dependent shifts in cell phenotypes across various evidence combinations. Eight evidence combinations (Combo_0-Combo_7) were defined based on the presence or absence of nutrient depletion, lysosomal alkalinization, ROS, and fatty acid oxidation and used as inputs for Bayesian inference of cell phenotypes (Figure 5C, Extended data Figure S28). Basal conditions (Combo_0–1) showed higher probabilities of cell death (∼50%). ROS or FAO alone increased the likelihood of elongation (up to 50% in Combo 2-3), while lysosomal alkalinization further favored the elongated phenotype (up to 50% in Combo 4) (Figure 5C, Extended data Figure S28). Nutrient depletion markedly increased cell death probability (50% in Combo 6), and the combination of all stressors (Combo 7) resulted in the highest death probability (60%). Overall, lysosomal alkalinization promoted elongation, whereas nutrient depletion, ROS, and FAO cooperatively regulated cell death/survival irrespective of lysosomal alkalinization.

**Figure 5:**
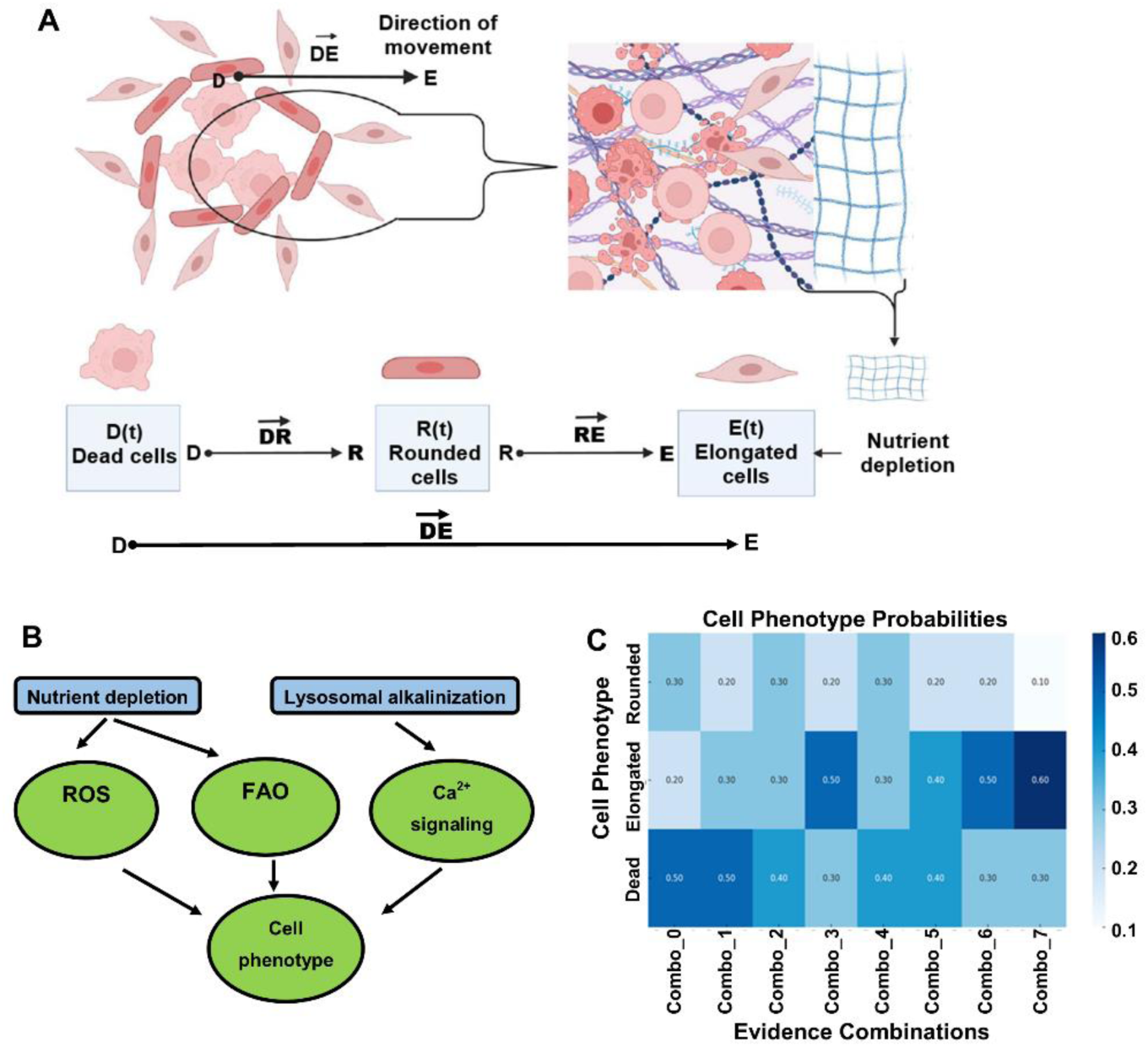
A Schematic representation of the proposed model describing cell states and directional movement in ND+Baf. **A**. The viable cells exhibit fatty acid oxidation and accumulation of apoptotic signaling without undergoing cell death; however, only a subset of living cells undergo stress-induced elongation (E) and acquire directional motility driven by the DE vector, enabling migration toward more favorable niches. Dead cells (D) contribute to local environmental signaling (DR), influencing neighboring cells and potentially promoting dormancy. Elongated (E) cells actively remodel lysosomal heterogeneity during migration, thereby supporting survival under nutrient-deprived conditions. In contrast, rounded (R) cells remain centrally confined, lacking heterotypic signaling and lysosomal heterogeneity, driven Ca²⁺-dependent exocytosis, and consequently fail to acquire migratory capacity. **B**. Bayesian Network illustrating the relationship between nutrient depletion, lysosomal alkalinization, ROS, fatty acid oxidation, calcium signaling, and cell phenotype. **C.** Heatmap of Bayesian network-inferred probabilities for cell phenotypes (Rounded, Elongated, Dead) across eight evidence combinations integrating nutrient depletion, lysosomal alkalinization, ROS, and lipid oxidation.

## Discussion

The availability of nutrients to tumor cells is largely dependent on the presence of an effective vasculature and competition for metabolites between the different cell types in the tumor microenvironment (TME). ^26^ Cancer cells develop numerous strategies for survival in such an environment, including increased uptake of available nutrients, *de novo* synthesis of amino acids or lipids or nutrient recycling through autophagy. ^27–29^ *De novo* synthesis utilizes resources and reducing electrons that may be re-allocated from other pathways, while autophagy primarily preserves certain crucial nutrients and is primarily used to enable cell survival during periods of nutrient deprivation. ^26^ We have previously shown that nutrient depletion in colorectal cancer cells led to extensive cell death; however, the surviving cells were highly viable, could form tumors *in vivo* and showed characteristics such as drug resistance through lysosomal trapping of drugs,^5^ and partial EMT.^6^ Alkalinization of the lysosomes with Baf further exacerbated cellular motility in Caco-2 cells. ^4^

In the current study, we evaluated the mechanism by which the cells retained their high viability and motility under low nutrient availability and lysosomal alkalinization. For this, we focused on Caco-2 cells and evaluated RNA sequencing data generated from NR +/-Baf and ND+/-Baf cells. Our aim was to identify the genes and pathways that were exclusively activated in the ND+Baf Caco-2 cells. We observed that although a majority of the genes were downregulated in these cells, the upregulated genes formed critical and well-connected signaling network that could contribute towards cell viability and motility. Apoptosis, while essential for cell death, has increasingly been implicated in signaling without causing cell death. ^30^ A modest alteration of mitochondrial outer membrane potential (MOMP) was shown to lead to the release of cytochrome c without activating cell death; rather it led to the activation of the integrated stress response and cell survival. ^31^ Such “persister” cells were also shown to be drug resistant.^32^ We have observed that the expression of *CYCS*, the gene encoding cytochrome c, showed synergy, i.e., significant upregulation in the ND+Baf cells, but not in the ND or the NR+Baf cells when NR was considered as control. In addition to cytochrome c, several other apoptosis related proteins were specifically upregulated in the ND+Baf cells. However, the high viability of these cells ^4^ points towards the use of apoptosis signaling as a means for cell survival rather than death.

Nutrient depletion led to an increase in the FAO score compared to the NR cells, which was further increased when the cells were treated with Baf. Corroborating this, we observed the greatest decrease in the lipid droplets in the ND+ Baf cells, along with an increase in phosphorylation of Acetyl CoA Carboxylase (ACC) at Ser 80, suggesting that these cells were utilizing FAO for energy generation. ACC Ser 80 phosphorylation can be mediated by AMPK when the cell has low energy levels, and leads to the activation of the FAO enzyme carnitine palmitoyltransferase 1 (CPT1) for energy generation.^33^ The mobilization of fatty acids from lipid droplets is particularly important during nutrient depletion, during which catabolism of stored lipids ensures an efficient and rapid energy supply to cancer cells.^34^ Beta oxidation can also be accompanied by the release of ROS, which, while lethal at high doses, can act as an effective signaling molecule at lower doses. ^34^ Indeed, we observed an increase in the ROS score in both ND and ND+Baf cells (both of which showed a decrease in lipid droplet content) and higher mitochondrial ROS levels in the ND+Baf cells. ROS can act as a nodal point as the convergence of environmental and intracellular cues, such as metabolic stress, or hypoxia, and orchestrate the decision between life and death of the cell via programmed cell death mechanisms. ^35^

Functional interdependence, contact, and cross talk between lysosomes and mitochondrial signaling have been widely reported and can contribute towards changes in cell fate well beyond the boundaries of these organelles. TRPML1, a well-established lysosomal calcium efflux channel, was shown to mediate the communication between mitochondria and lysosomes and was important for mitochondrial structure and bioenergetics.^36^ We have observed a remarkable increase in TRPML1 expression in the ND+Baf cells,^4^ which suggests that the increased Ca^2+^ release from the lysosomes may have contributed towards increased beta oxidation in the mitochondria. Additionally, TRPML1 was also shown to increase apoptosis through the inhibition of mitophagy and accumulation of ROS. ^37^ Indeed, we have observed the activation of mitophagy in the ND+Baf Caco-2 cells, suggesting that the lysosomes could initiate mitophagy; however, due to the lysosomal alkalinization, very likely could not complete it. Thus, the treatment of the ND cells with Baf led to FAO for energy generation, initiation of mitophagy and increase in mitochondrial ROS all of which were accompanied by the induction, but not completion of apoptosis. Apoptosis signaling was likely to coordinate these responses [as also corroborated by the close interaction network formed genes related to FAO, ROS, and apoptosis in the ND+Baf Caco-2 cells (GSE245402)] to ensure cell survival, since most of these phenotypes could be reversed with the pan-caspase inhibitor QVD-OPH.

Lysosomal functions rely on an acidic luminal pH that is tightly regulated, which may shift under conditions such as nutrient deprivation and oxidative stress, resulting in changes in their positioning. ^16,38^ We observed that lysosomes were often localized very close to the long extensions in the ND+Baf cell population. The elongated cells also showed the highest molecular heterogeneity in the Gamma distribution modeling compared to the non-elongated (rounded) or dead cells, suggesting the activation of unique signaling pathways in these cells. Due to the increased motility of the ND+Baf cells and the localization of the lysosomes close to the long extensions, we hypothesized a role of induction of lysosomal exocytosis in cell motility. Ca^2+^ released from TRPML1 is known to regulate multiple lysosomal functions, including membrane trafficking and exocytosis.^39^ PIKfyve (also known as 1-phosphatidylinositol 3-phosphate 5-kinase) can convert phosphatidylinositol-3-phosphate (PI(3)P) into PI(3,5)P and is critical for endolysosomal trafficking.^40^ To establish whether the ND+Baf cells exhibited lysosomal exocytosis, we treated the NR, NR+Baf, ND and ND+Baf Caco-2 cells with the PIKfyve inhibitor Vacuolin-1 and observed a decrease in extensions in the ND+Baf cells. Additionally, we observed a decrease in the motility of Caco-2 cells from ovografts to the liver in the CAM, as well as a restoration of lysosomal positioning towards the perinuclear region in the cells treated with Vacuolin-1. Lysosomal exocytosis, especially in cancer cells, has been widely implicated in extracellular acidification, tumor dissemination through extracellular matrix (ECM) degradation via the released cathepsins, treatment resistance to lysomotropic drugs such as doxorubicin, and the release of exosomes.^41^ Thus, the increased motility of the ND+Baf Caco-2 cells may have resulted from lysosomal exocytosis and subsequent ECM remodeling.

We next used Bayesian network modeling to simulate the different cellular states observed in the ND+Baf cells. The stressors, i.e., nutrient depletion and lysosomal alkalinization played different roles individually and in combination. While all stressors led to cell death to a certain extent, metabolic plasticity via alternative energy utilization (FAO) and survival was observed in both ND and ND+Baf cells. More importantly, cellular elongation and Ca^2+^dynamics were exclusively associated with ND+Baf cells. Thus, the combination of the stressors allowed the selection of a group of cells that could survive and have enhanced motility. We also propose that lysosomal exocytosis was not a random event, but was spatially regulated, which could provide directional motility of the cancer cells.

Overall, our study introduces an integrative framework to define the molecular drivers underlying cell behavior under metabolic stress. We have uncovered a crucial role of lysosomes in orchestrating a well-coordinated series of the events with spatial relevance. The loss of enzymatic function of the lysosomes in cells with limited nutrient availability led to enhanced communication with the mitochondria for FAO and energy generation, while the lysosomes located towards the periphery mediated calcium driven lysosomal exocytosis for cell motility and dissemination. Our data provide additional mechanisms to support a critical role of lysosomes in cancer, suggesting that alkalinization of lysosomes, especially in cells that are under metabolic stress, may select specifically for cells that have metabolic plasticity and high motility.

## Acknowledgements

Seljan Khatun Abdullazade and Özün Özcan for technical assistance and valuable discussions. The study was supported in part by TÜBİTAK (Project no: 124Z190 to HHH) and ODTÜ BAP (Project no: GAP-108-2023-11313, to SB).

## Author contributions

LNN conceptualized the study, carried out the *in silico* experiments and mathematical modeling and wrote the paper, HHH carried out most of the wet lab experiments with help from HBÇY and AEGT. NL supervised the imaging assays; ACA supervised the *in silico* and modeling experiments. SB conceptualized the study, obtained funding with help from HHH and wrote the paper. All authors read and approved the final version of the manuscript.

## Declaration of interests

The authors declare no competing interests.

## Notes

https://www.omicsdi.org/dataset/geo/GSE245402

### Competing Interest Statement

The authors have declared no competing interest.

