## Supplementary Methods for "Lysosomal Alkalinization Selects for Metabolically Plastic, Motile Cancer Cells under Nutrient Stress"

### Role of the lysosome in mediating survival, metabolic plasticity and motility in nutrient deprived cancer cells.

##### 1. Detection of Significantly Differentially Expressed and Overlapping Genes:

DGE results for the experimental combinations (Figure S2) were first separated into upregulated (log2FC > 1) and downregulated (log2FC < –1) genes (adjusted p-value < 0.01). Entrez Gene IDs were then converted to gene symbols using the SynGO ID conversion tool (https://www.syngoportal.org/convert). All gene sets and DGE results from RNA-seq analyses for each condition (Figure S4) were imported into R Studio to determine the common genes shared between the gene sets and DEGs under each experimental condition. Common genes from both up- and downregulated groups across all control–treatment combinations were identified and subsequently used for further analyses and modeling. The STRING database (https://string-db.org/) was employed to identify proteins involved in strong protein–protein interaction networks. Gene Ontology (GO) Resource was then used to explore the biological processes enriched among these highly connected proteins. Co-expressed proteins were detected for specific cell groups (NR, ND, NR+Baf, and ND+Baf) and compared to reveal similarities and differences in biological properties among the experimental groups.

##### 2. Development of Scoring Algorithms:

All scoring algorithms are available at https://github.com/lnnehri/ND-Baf. In the provided code, genes were categorized as inducers or suppressors based on the literature, each supported by DOI numbers. The relevant gene lists for each group were derived through gene set analysis, and scoring was performed accordingly.

The normalization approach followed the same principle across all scoring methods: to standardize gene expression values across samples, two types of min-max normalizations were applied based on the gene's functional role. For genes classified as inducers, values were normalized using a standard min-max scaling formula: ( x _− m _i _n _( x _) ) / ( m _a _x _( x _) − m _i _n _( x _) ) $(x-\min(x))/(\max(x)-\min(x))$ This scaled the expression values between 0 and 1, with higher values indicating stronger induction. For suppressor genes, an inverse version of the same formula was used: 1 − ( ( x _− m _i _n _( x _) ) / ( m _a _x _( x _) − m _i _n _( x _) ) ) $1-((x-\min(x))/(\max(x)-\min(x)))$

This approach ensured that higher raw expression values of suppressors, normally associated with reduced pathway activity, resulted in lower normalized scores, thereby maintaining biological consistency across both gene types during scoring.

An Adjusted Score was calculated for each sample by summing the normalized expression values of all selected genes related to the pathway of interest. This composite score integrated the relative contribution of both inducers and suppressors (after appropriate normalization) into a single quantitative value. The higher the score, the stronger the overall activation of the pathway, reflecting cumulative gene-level activity. This approach allowed for the comparison of pathway activity across different experimental conditions by reducing complex multi-gene expression profiles into a simplified, interpretable metric.

**3. Development of Synergy Algorithms:**

To identify potential synergistic effects in specific gene expression profiles, a set of comparative analyses were performed across three experimental conditions: ND, NR+Baf, and ND+Baf. The approach focused on genes that were significantly differentially expressed (adjusted p-value < 0.01) and unique to the selected genes' pathways. Genes were classified as showing either synergistic or additive effects based on predefined threshold rules comparing individual (ND or Baf) and combined (ND+Baf) expression patterns. These predefined rules involve conditional comparisons between individual (ND or Baf) and combined (ND+Baf) gene expression levels, identifying synergy when the combined response exceeded or deviated from the expected additive or multiplicative effect. Genes were labeled as showing synergy in the following cases:

1. When the gene was expressed only in ND+Baf

2. When gene expression of missing in both ND and Baf but was expressed in ND+Baf; or

3. When the expression in the combined condition significantly exceeded or deviated from what would be expected based on either individual treatment alone.

Additional rules considered scenarios where both ND and Baf were upregulated or downregulated, and ND+Baf showed a non-additive or non-multiplicative response. If none of these synergy conditions were met, the gene was classified as having an additive response. These logic-based rules enabled systematic identification of non-linear interactions between treatment conditions at the gene expression level.

To distinguish synergistic from additive regulation more precisely, quantitative thresholds of ±0.3 and ±0.4 were incorporated into the decision rules. In cases of upregulation (ND and Baf > 0), synergy was assigned when the combined condition (ND+Baf) exceeded the multiplicative effect of the individual treatments by more than 0.4, while a decrease beyond this margin indicated suppression. Similarly, for downregulated genes (ND and Baf < 0), synergy was considered when ND+Baf deviated from the expected product by more than 0.3 in either direction. These empirically defined margins allowed for the detection of biologically meaningful non-additive shifts in gene expression. In other words, if the expression-change in the combined condition (ND+Baf) exceeded the expected additive or multiplicative effect based on the pre-defined conditions and thresholds of the individual treatments, the gene was labeled as showing synergy.

Genes were annotated for the direction of regulation (increase or decrease) and for their trend of overall expression (upregulated or downregulated), based on log-fold changes. This strategy enabled the identification of genes with synergistic induction or repression in response to combined treatment (ND+Baf), providing mechanistic insights into the regulation of the selected pathway under dual perturbation. The synergy analysis and scoring workflows for FA oxidation, ROS response, apoptosis, and related gene sets were developed specifically for this study and are available at the following

**4. Analysis of single cell RNA seq data**

To validate that the expression of apoptosis related genes was also high in metastatic cells (as observed in the motile ND+Baf Caco-2 cells compared to the non-motile controls NR), the DEGs between metastatic and non-metastatic cell groups provided in Supplementary Table S2 of Bernal et al. (list of genes showing differential expression in CC14 clusters determined by scRNAseq) [2] were compared with the DEGs for NR vs. ND+Baf, logFC>1 and logFC<-1 from in bulk RNA-seq from GSE245402 [1].

Single-cell analysis of the CC14 clone cells from PRJNA611719 Bioproject sample was carried out using RNAStarSolo through the Galaxy platform. Clustering, trajectory inference, and embedding (tl.louvain) were conducted, and annotated data matrix PCA, KNN, UMAP, and clustering=0.2 were used to generate clusters. Gene grouping based on 3 PCAs was obtained. Using pl.highest_expr_genes, the most highly expressed genes in all analyzed cells were identified by calculating the contribution of each gene to the total gene expression and limiting it to those with the highest contributions. We evaluated how these common genes are distributed within single-cell clusters using Louvain clustering. We implemented a stochastic, grid-based agent-based simulation to study how local cellular interactions give rise to spatial heterogeneity at the tissue scale. Cells were modeled as agents that probabilistically generate a heterogeneity signal upon peripheral activation and cell-cell contact, representing alkalinized lysosomal/exocytosis- and calcium release–associated events. These events accumulate on a two-dimensional grid as a signal field (H), which evolves through decay and spatial diffusion. Spatial patterns were controlled by four key parameters: the frequency of signal-generating events, the magnitude of each event, the signal decay rate, and the diffusion coefficient. Model dynamics were simulated using Monte Carlo sampling at discrete time steps, where release activation, signal emission, state transitions, and movement are updated probabilistically. Although individual updates were locally memoryless, complex global patterns emerged from repeated stochastic updates across space and time, without explicitly defining a Markov transition matrix.

**5. Modeling of spatial heterogeneity**

In the simulation, key parameters were designed to control distinct aspects of signal generation and persistence. The baseline and context-dependent probabilities (p_alk_base, boost_edge, boost_contact) determined how readily a release event was triggered at the cellular periphery, particularly upon cell–cell contact. Once triggered, p_emit_if_edge_alk_contact governed whether a release event actually occurred, introducing stochasticity to the process. The magnitude of each release event was encoded by the signal increment (H += 0.9), defining how much each event contributed to the local heterogeneity field. Signal persistence was controlled by the decay parameter (signal_decay), which defined how long the deposited signal remained over time, while the diffusion coefficient (signal_diffuse) determineed whether the signal spread into neighboring regions or remained spatially confined. Together, these parameters were considered to regulate the frequency, strength, lifetime, and spatial extent of heterogeneity signals on the grid.

To enhance the emergence of spatial heterogeneity in the simulation, we simultaneously increased multiple regulatory parameters controlling signal generation and persistence. Specifically, the baseline and contact-dependent probabilities of release events were elevated, resulting in more frequent signal emission per cell. In parallel, the contribution of each individual release to the heterogeneity field was increased, allowing even single events to produce stronger local signals. Signal decay was reduced, enabling the accumulated signals to persist longer over time, while spatial diffusion was constrained to limit signal spread and promote local retention. Together, these coordinated parameter adjustments led to more intense, long-lived, and spatially confined heterogeneity signals, manifested as an increased number and strength of localized hotspots.

**6. Accumulation modeling:**

For accumulation-to-threshold modeling, we treated fatty acid oxidation and calcium accumulation as Gamma processes. Fatty acid oxidation accumulation used dgamma(x, shape = L_threshold, scale = 1/L_hour) to evaluate the probability density at a normalized half-time x (e.g., 0.5 ≈ 48/96 h), where L_threshold encodes the fatty acid oxidation amount required to trigger a response and L_hour the per-hour accumulation rate. An analogous formulation was used for calcium with shape = Ca_threshold and scale = 1/Ca_hour. To estimate time-to-threshold, we iteratively increased time in 1-h steps and computed the cumulative probability pgamma (accumulation_value * time, shape, scale) until it exceeded a predefined probability threshold; the first time meeting this criterion was recorded as the decision time.
