## Extended data for "Lysosomal Alkalinization Selects for Metabolically Plastic, Motile Cancer Cells under Nutrient Stress"

### Role of the lysosome in mediating survival, metabolic plasticity and motility in nutrient deprived cancer cells.

#### 1. Comparison of response of different cell lines to nutrient depletion and lysosomal alkalinization

##### 1.1 Temporal regulation of cell death

The proliferation of three different colon cancer cell lines (Caco-2, T84, and RKO) cultured for 24, 48, 72, and 96 hours in the nutrient rich (NR, complete medium), nutrient depleted (ND, 10% of complete medium), ND+ 5nM Bafilomycin A1 (ND+Baf) and NR+Baf was determined using an MTT assay (Figure S1). 10,000 cells were plated in 96 well plates and allowed to attach overnight in complete medium. Next, the medium was changed to either ND, or Baf (5nM) was added to the cells maintained in NR or ND medium. The cells were collected after 24, 48, 72 and 96 hours and an MTT assay was carried out using standard techniques.

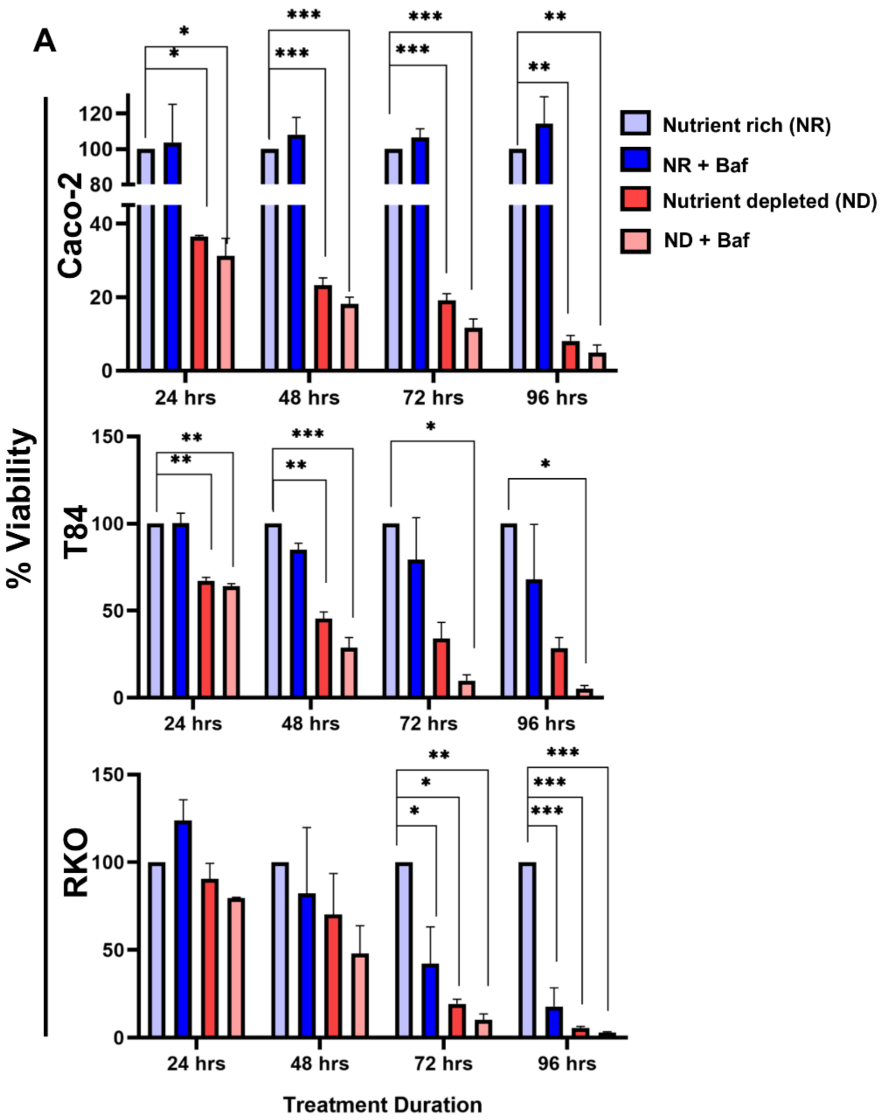


**Figure S1: Comparison of proliferation of Caco-2, T84 and RKO cells under nutrient depletion (ND) and/or lysosomal alkalinization**. The cells were incubated in nutrient rich (complete medium, NR, light blue bars), NR cells treated with 5nM Bafilomycin A1 (dark blue bars), nutrient depleted medium (ND, coral bars) or ND cells treated with 5nM Baf (pink bars) for 24-96 hours.

Using the data shown in Figure S1, two alternative kinetics were fitted: (i) a logistic model defined as $y(t) = L / [1 + \exp(-k \times (t - t_0))]$ and (ii) an exponential decay model defined as $y(t) = A \times \exp(-b \times t)$. Model parameters were estimated using nonlinear least-squares fitting implemented in Python (SciPy curve_fit function). Initial parameter values were set as L = maximum(y), k = 0.1, and $t_0$ = median(t) for the logistic model, and A = maximum(y), b = 0.05 for the exponential model. Goodness of fit was evaluated using the residual sum of squares (RSS) and the Akaike Information Criterion (AIC), calculated as $AIC = n \times \ln(RSS / n) + 2 \times p$, where p represents the number of model parameters (p = 3 for logistic and p = 2 for exponential) (Table S1). Model preference was determined by the difference ΔAIC = AIC_exp − AIC_log; values greater than 2 favored the logistic model, values smaller than −2 favored the exponential model, and values between −2 and 2 indicated no clear preference. For visualization purposes, smooth curves were generated using quadratic B-spline interpolation (make_interp_spline, k = 2), and vertical reference lines were added at 24, 48, 72, and 96 hours. A schematic sigmoid curve was also rendered using L = 45,000, $t_0$ = 72 hours, and k = 0.1. Analyses were conducted in PyCharm.

**Table S1.** Summary of model comparison results (ΔAIC = AIC_exp − AIC_log). Positive ΔAIC indicates better logistic (sigmoidal) fit; negative values indicate better exponential fit.

| **Condition** | **Cell line** | **ΔAIC** | **Preferred model** |
|---|---|---|---|
| ND | Caco-2 | −1.6 | Similar (no preference) |
| ND | T84 | −2.0 | Exponential |
| ND | RKO | +13.5 | Logistic |
| ND+Baf | Caco-2 | −1.3 | Similar (no preference) |
| ND+Baf | T84 | +0.3 | Similar (no preference) |
| ND+Baf | RKO | +15.0 | Logistic |
| NR+Baf | Caco-2 | −2.0 | Exponential |
| NR+Baf | T84 | −2.0 | Exponential |
| NR+Baf | RKO | +38.5 | Logistic |

#### 2. Immunocytochemistry to determine lysosomal positioning

The position of the lysosomes T84, RKO and Caco-2 cell lines that were used to determine the positioning of the lysosomes with respect to the nucleus (Figure S2). We observed a shift in lysosomal positioning from a uniform distribution in the NR cells of T84 and RKO to the perinuclear regions in the ND cells (Figure S2). With ND+Baf we observed a slight shift in the lysosomes from the perinuclear to the peripheral regions (Figure S2) in T84 and RKO cells. Caco-2 cells showed a reasonably uniform distribution (perinuclear and peripheral) in NR, ND and ND+Baf cells with no major shifts (Figure 2).

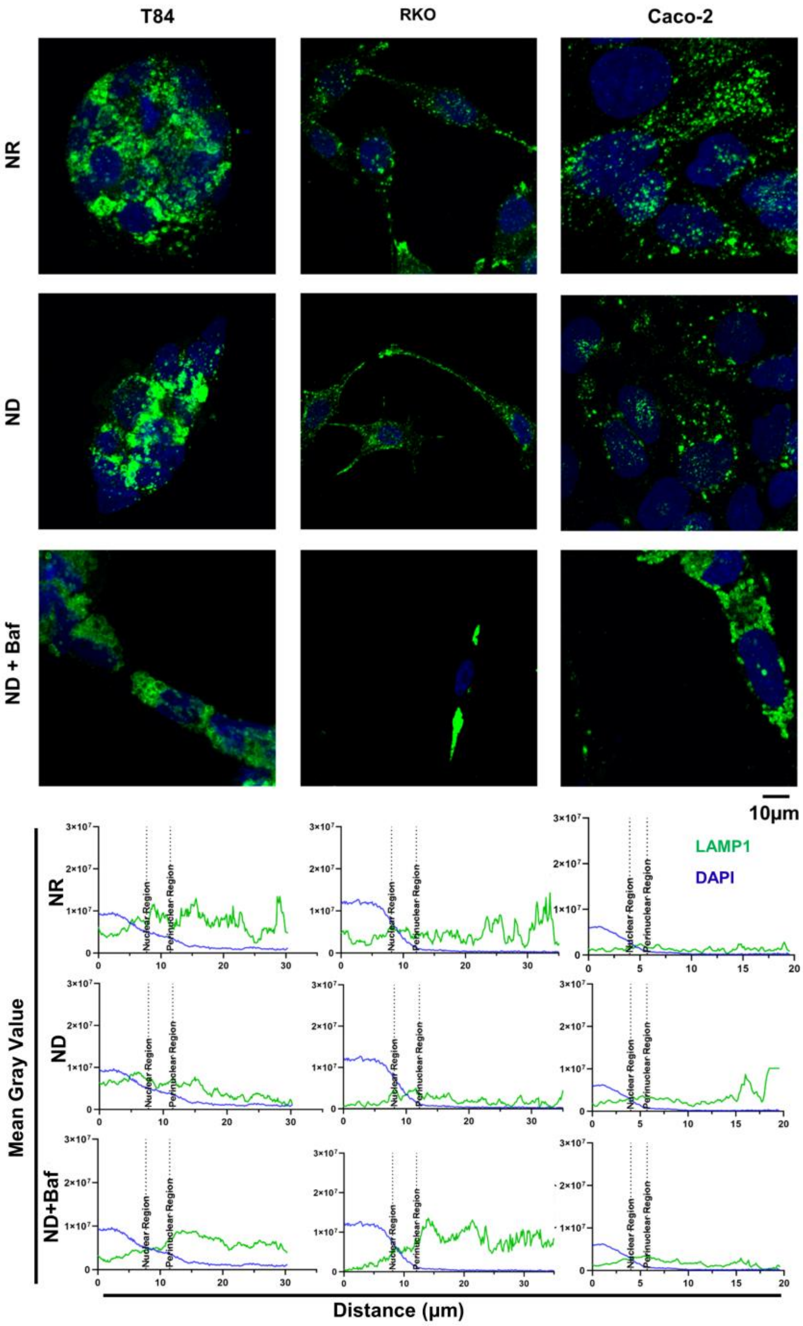


**Figure S2: Positioning of the lysosome in T84, RKO and Caco-2 cells.** Upper panel: Confocal microscopy images of NR, ND and ND+Baf cells. The lysosomes are stained with LAMP1, nuclei with DAPI. Lower panel: Quantification of lysosomal positioning. A total of 80 cells were analyzed, with 20 cells per experimental condition, and normalized fluorescence intensity values were used for comparative analysis between the groups (lower panel). The images were acquired with the LSM800 Confocal Laser Scanning Microscope (Zeiss) with 40X and 63X water-based immersion objective.

We next determined the acidity of the lysosomes in Caco-2 and T84 cells (Figure S3) and observed a remarkable decrease in active lysosomes and in both cell lines.

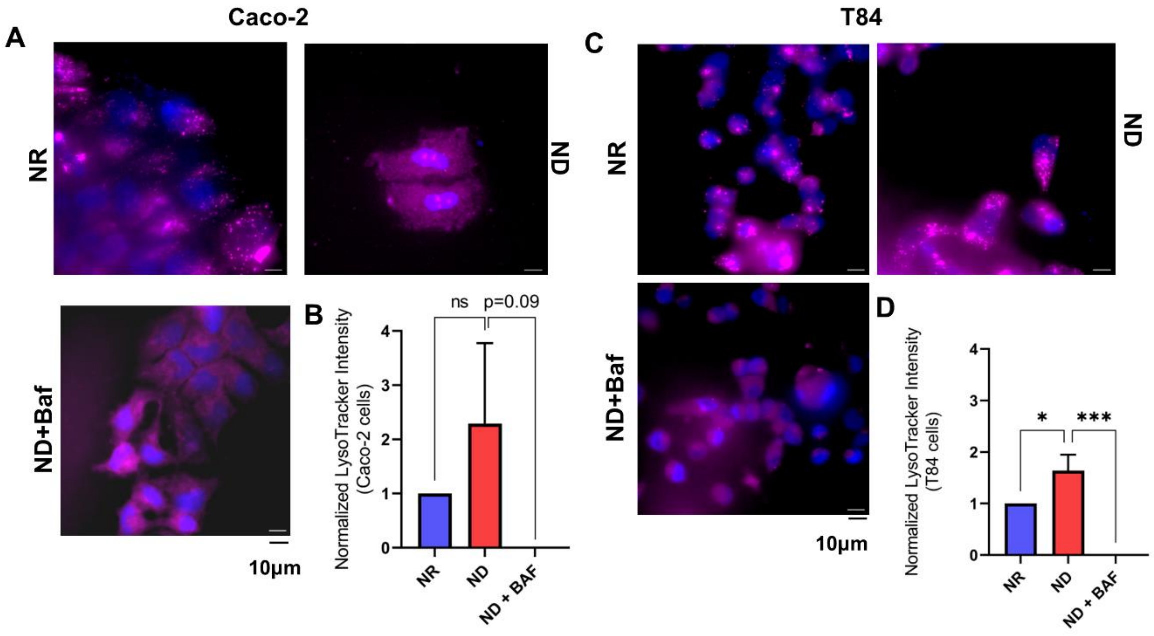


**Figure S3: Lysosomal activity in the NR, ND and ND+Baf Caco-2 and T84 cells. A & C.** SiR Lysosome staining of Caco-2 and T84 cells showing a loss of punctate signals upon treatment with Baf. **B & D**. Lysotracker Intensity measurement, showing the complete loss of signal when the ND cells were treated with Baf.

#### 2. Comparative Matrix of Differential Gene Expression in Caco-2 cells

All analyses were conducted in R (with functions of dpois, dgamma, pgamma, rgamma, t.test, aov, etc.) with visualization using ggplot2 and panel assembly using cowplot, and in Python using NumPy, SciPy, and Matplotlib, etc. for modeling, fitting, and visualization.

RNA seq data from Caco-2 cells grown under nutrient rich (NR), nutrient deficiency (ND), NR cells treated with the v-ATPase inhibitor Bafilomycin A1 (BAF) and ND cells treated with BAF (GSE245402) were analyzed. The data were compared as pairs considering one condition as the control and the other as the experimental group (e.g.: NR vs NR+BAF, NR vs ND, NR vs ND+BAF, ND vs NR+BAF, ND vs NR, ND vs ND+BAF, NR+BAF vs NR, NR+BAF vs ND, NR+BAF vs ND+BAF, ND+BAF vs NR, ND+BAF vs NR+BAF, ND+BAF vs ND). These comparisons were used for differential gene expression (DGE) analyses (Figure S4).

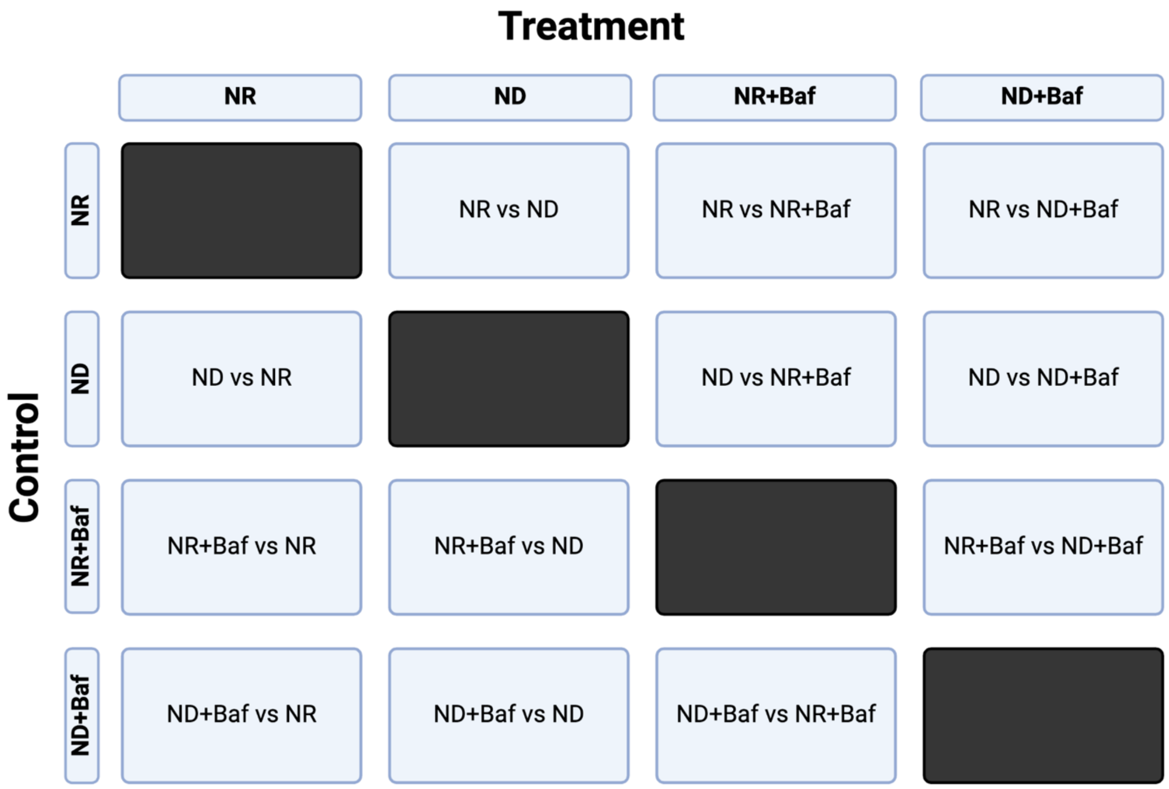


**Figure S4.** The matrix of differential gene expression (DGE) analyses. Each node of the matrix except the filled boxes were analyzed for both up and down regulated genes via separate analysis.

##### 3. Evaluation of publicly available Gene Sets:

To identify genes of interest within the differentially expressed genes (DEGs), multiple gene sets obtained from various repositories were incorporated (Table S2).

**Supplementary Table S2** Sources of selected gene sets.

| **Process** | **Gene Set / Source** | **URL** |
|---|---|---|
| Cytoskeleton | CYTOSKELETON | (https://www.gsea-msigdb.org/gsea/msigdb/cards/CYTOSKELETON) |
| Starvation and dietary restriction | GenDR | https://genomics.senescence.info/diet/ |
| EMT | EMTome | http://www.emtome.org/ |
| Lysosome | The Human Lysozyme Gene Database - hLGDB | http://lysosome.unipg.it/ |
| Senescence | CellAge Database | https://genomics.senescence.info/cells/ |
| Anoikis resistance | GeneRIF Biological Term Annotations | https://maayanlab.cloud/Harmonizome/gene_set/anoikis/GeneRIF+Biological+Term+Annotations |

| | | |
|---|---|---|
| Stem-cell | CSC database | http://dibresources.jcbose.ac.in/ssaha4/bcscdb/download.php |
| Calcium signaling pathways | KEGG Pathways | https://maayanlab.cloud/Enrichr/enrich?dataset=5cc763ed8b49a5edd91343b256ccf30f |
| Mitochondrial Fatty Acid Beta-Oxidation | KEGG Pathways | https://maayanlab.cloud/Enrichr/enrich?dataset=1a25af6dc86f9f36a5b1e767f800904b |
| Fatty Acid Biosynthesis | KEGG Pathways and Human Gene Set: WP_FATTY_ACID_BIOSYNTHESIS | https://maayanlab.cloud/Enrichr/enrich?dataset=f72be244c9637ecdc6b449f2e0f4e4f6 , https://www.gsea-msigdb.org/gsea/msigdb/cards/WP_FATTY_ACID_BIOSYNTHESIS |
| Apoptosis | Human Gene Set: KEGG_APOPTOSIS | https://www.gsea-msigdb.org/gsea/msigdb/cards/KEGG_APOPTOSIS |
| Mitochondria | Human Gene Set: MOOTHA_MITOCHONDRIA | https://www.gsea-msigdb.org/gsea/msigdb/cards/MOOTHA_MITOCHONDRIA |
| Fatty acid metabolism | Human Gene Set: HALLMARK_FATTY_ACID_METABOLISM | https://www.gsea-msigdb.org/gsea/msigdb/human/geneset/HALLMARK_FATTY_ACID_METABOLISM.html |
| Glucose metabolism | Human Gene Set: REACTOME_GLUCOSE_METABOLISM | https://www.gsea-msigdb.org/gsea/msigdb/cards/REACTOME_GLUCOSE_METABOLISM |
| Amino acid metabolism | Human Gene Set: REACTOME_METABOLISM_OF_AMINO_ACIDS_AND_DERIVATIVES | https://www.gsea-msigdb.org/gsea/msigdb/human/geneset/REACTOME_METABOLISM_OF_AMINO_ACIDS_AND_DERIVATIVES.html |
| Lipid accumulation | GWASdb SNP-Phenotype Associations | https://maayanlab.cloud/Enrichr/enrich?dataset=5610bf49d63e6b3b527de54cb813d94c |
| Lipogenesis | GeneRIF Biological Term Annotations | https://maayanlab.cloud/Harmonizome/gene_set/lipogenesis/GeneRIF+Biological+Term+Annotations |
| Immediate early genes | Mammalian Immediate Early Genes | https://esbl.nhlbi.nih.gov/Signaling-Pathways/Im-early/ |
| Growth Factor | GOMF_GROWTH_FACTOR_ACTIVITY | https://www.gsea-msigdb.org/gsea/msigdb/cards/GOMF_GROWTH_FACTOR_ACTIVITY |
| Endoplasmic Reticulum (ER) | COMPARTMENTS Experimental Protein Localization Evidence Scores Data Set | https://maayanlab.cloud/Harmonizome/gene_set/endoplasmic+reticulum/COMPARTMENTS+Experimental+Protein+Localization+Evidence+Scores |
| Angiogenesis | Angiogenesis | https://maayanlab.cloud/Harmonizome/gene_set/Angiogenesis/PANTHER+Pathways |
| Cell to cell communication | REACTOME_CELL_CELL_COMMUNICATION | https://www.gsea-msigdb.org/gsea/msigdb/cards/REACTOME_CELL_CELL_COMMUNICATION.html |

| | | |
|---|---|---|
| Nutrient response | GOBP_RESPONSE_TO_NUTRIENT | https://www.gsea-msigdb.org/gsea/msigdb/human/geneset/GOBP_RESPONSE_TO_NUTRIENT.html |
| ROS response | HOUSTIS_ROS | https://www.gsea-msigdb.org/gsea/msigdb/cards/HOUSTIS_ROS |
| Immune escape | LIN_TUMOR_ESCAPE_FROM_IMMUNE_ATTACK | https://www.gsea-msigdb.org/gsea/msigdb/cards/LIN_TUMOR_ESCAPE_FROM_IMMUNE_ATTACK |
| Immune system processes | IMMUNE_SYSTEM_PROCESS | https://www.gsea-msigdb.org/gsea/msigdb/cards/IMMUNE_SYSTEM_PROCESS |
| 'Eat-me' signals of apoptotic cells | GOBP_PHOSPHATIDYLSERINE_EXPOSURE_ON_APOPTOTIC_CELL_SURFACE | https://www.gsea-msigdb.org/gsea/msigdb/geneset_page.jsp?geneSetName=GOBP_PHOSPHATIDYLSERINE_EXPOSURE_ON_APOPTOTIC_CELL_SURFACE |
| Extracellular Matrix (ECM) Organization | Extracellular Matrix Organization Gene Set | https://maayanlab.cloud/Enrichr/enrich |
| Exocytosis | EXOCYTOSIS | https://www.gsea-msigdb.org/gsea/msigdb/cards/GOBP_EXOCYTOSIS |
| Actin related cell contraction | GOBP_ACTIN_MEDIATED_CELL_CONTRACTION | https://www.gsea-msigdb.org/gsea/msigdb/cards/GOBP_ACTIN_MEDIATED_CELL_CONTRACTION |
| Fatty Acid oxidation | Fatty Acid Oxidation | https://maayanlab.cloud/Enrichr/enrich |
| ROS metabolism | Reactive Oxygen Species Metabolic Process | https://maayanlab.cloud/Enrichr/enrich |
| Lipid metabolism | Human Gene Set: REACTOME_METABOLISM_OF_LIPIDS | https://www.gsea-msigdb.org/gsea/msigdb/cards/REACTOME_METABOLISM_OF_LIPIDS |
| Cell motility | GO Biological Process Annotations | https://maayanlab.cloud/Harmonizome/gene_set/cell+motility/GO+Biological+Process+Annotations |
| Heterotypic Cell-Cell Adhesion | Regulation Of Heterotypic Cell-Cell Adhesion | https://maayanlab.cloud/Harmonizome/gene_set/regulation+of+heterotypic+cell-cell+adhesion/GO+Biological+Process+Annotations |
| Mitochondrial Permeability Transition Pore Complex genes | Mitochondrial Permeability Transition Pore Complex | https://maayanlab.cloud/Harmonizome/gene_set/mitochondrial+permeability+transition+pore+complex/COMPARTMENTS+Text-mining+Protein+Localization+Evidence+Scores |
| Detoxification of ROS | Human Gene Set: REACTOME_DETOXIFICATION_OF_REACTIVE_OXYGEN_SPECIES | https://www.gsea-msigdb.org/gsea/msigdb/cards/REACTOME_DETOXIFICATION_OF_REACTIVE_OXYGEN_SPECIES |
| Ferrotopsis | Human Gene Set: WP_FERROPTOSIS | https://www.gsea-msigdb.org/gsea/msigdb/human/geneset/WP_FERROPTOSIS.html?keywords=Ferroptosis |

| | | |
|---|---|---|
| Lipophagy | Human Gene Set: REACTOME_LIPOPHAGY | https://www.gsea-msigdb.org/gsea/msigdb/cards/REACTOME_LIPOPHAGY |
| Mitophagy | Human Gene Set: REACTOME_MITOPHAGY | https://www.gsea-msigdb.org/gsea/msigdb/human/geneset/REACTOME_MITOPHAGY |

**4. Protein-protein interaction in the different experimental groups:**

We analyzed the protein-protein interaction (PPI) network of significantly differentially expressed genes (logFC2>1 for upregulated and logFC2>-1, p_adj<0.01) between NR and ND Caco-2 cells to determine the effect of nutrient depletion, NR and NR+Baf to determine the effect of lysosomal alkalinization, NR and ND+Baf to determine the effect of nutrient depletion and lysosomal alkalinization and ND vs ND+Baf to determine the effect of lysosomal alkalinization in the nutrient depleted cells. The workflow used is described in Figure S5 and Table S3 and the PPIs under the different experimental conditions are shown in Figure S6 (upregulated proteins) and Figure S7 (downregulated proteins).

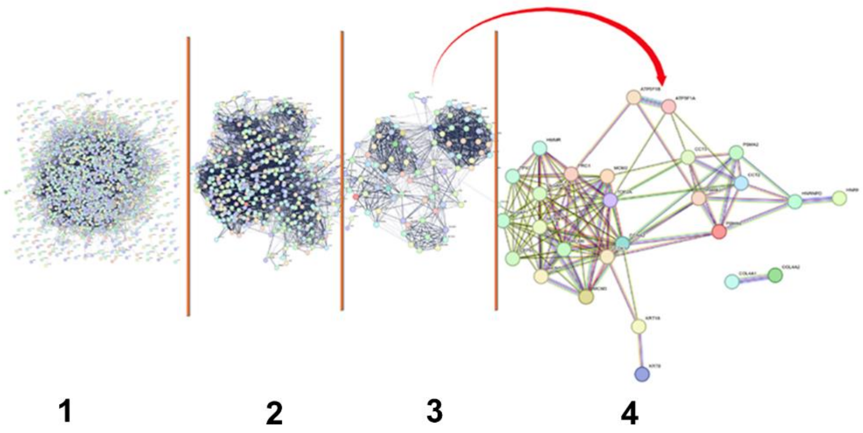


**Figure S5. Workflow illustrating the extraction of core proteins from PPI analyses.** Differentially expressed proteins identified under ND, NR+Baf, and ND+Baf conditions were filtered by statistical significance (adjusted p < 0.01), then subjected to network topology–based selection to retain only highly connected nodes. These core proteins represent the most functional components within the PPI networks and were used for subsequent comparative and enrichment analyses. The outputs numbered 1, 2, 3, 4 represent the steps that were used to identify the core functional proteins and is described in the Table 3S.

Table S3 summarizes the STRING-based protein–protein interaction (PPI) filtering and network analysis parameters used to identify core proteins within each experimental comparison. Differentially expressed proteins (|log2FC| > 1, adjusted p < 0.01) were subjected to stepwise STRING filtering based on combined interaction scores and co-expression thresholds. The sequential outputs represent progressively stringent interaction filtering and network refinement steps, including re-analysis of high-confidence interaction subsets and broader co-expression network evaluation.

**Table S3: STRING-based protein–protein interaction filtering and clustering parameters applied to different experimental comparison**

| Conditions | Output 1 | Output 2 | Output 3 | Output 4 |
|---|---|---|---|---|
| ND Upregulated | logFC2>1, p_adj<0.01 | Proteins with combined score=0.999 re-analyzed in STRING | Co-expression >0.500 & combined >0.750 | Combined ≥0.900 & co-expression ≥0.500 (from Output 1, MCL clustering) |
| ND Downregulated | logFC2<-1, p_adj<0.01 | Proteins with combined score=0.999 re-analyzed in STRING | Co-expression >0.990 & combined >0.999 | — |
| NR+Baf Upregulated | logFC2>1, p_adj<0.01 | Proteins with combined >0.950 re-analyzed in STRING | Co-expression >0.750 & combined >0.500 | — |
| NR+Baf Downregulated | logFC2<-1, p_adj<0.01 | Proteins with combined ≥0.990 re-analyzed in STRING | Co-expression >0.700 & combined >0.800 | Combined ≥0.900 & co-expression ≥0.900 (from Output 1) |
| NR *vs* NR+Baf Upregulated | logFC2>1, p_adj<0.01 | Combined >0.995 re-analysis | Co-expression >0.750 & combined >0.500 | Combined ≥0.950 & co-expression ≥0.950 |
| NR *vs* NR+Baf Downregulated | logFC2<-1, p_adj<0.01 | Combined =0.999 re-analysis | Co-expression >0.900 & combined =0.999 | Co-expression =0.990 & combined =0.999 |
| NR vs ND+Baf Upregulated | logFC2>1, p_adj<0.01 | Combined ≥0.900 re-analysis | Co-expression ≥0.750 & combined ≥0.500 | Combined ≥0.750 & co-expression ≥0.500 |
| NR vs ND+Baf Downregulated | logFC2<-1, p_adj<0.01 | Combined ≥0.950 re-analysis | Co-expression >0.750 & combined >0.500 | Combined ≥0.900 & co-expression ≥0.500 |
| ND vs ND+Baf Upregulated | logFC2>1, p_adj<0.01 | Combined ≥0.900 re-analysis | Co-expression ≥0.750 & combined ≥0.500 | Combined ≥0.750 & co-expression ≥0.500 |
| ND vs ND+Baf Downregulated | logFC2<-1, p_adj<0.01 | Combined ≥0.950 re-analysis | re-analysis Co-expression >0.750 & combined >0.500 | Combined ≥0.900 & co-expression ≥0.500 |

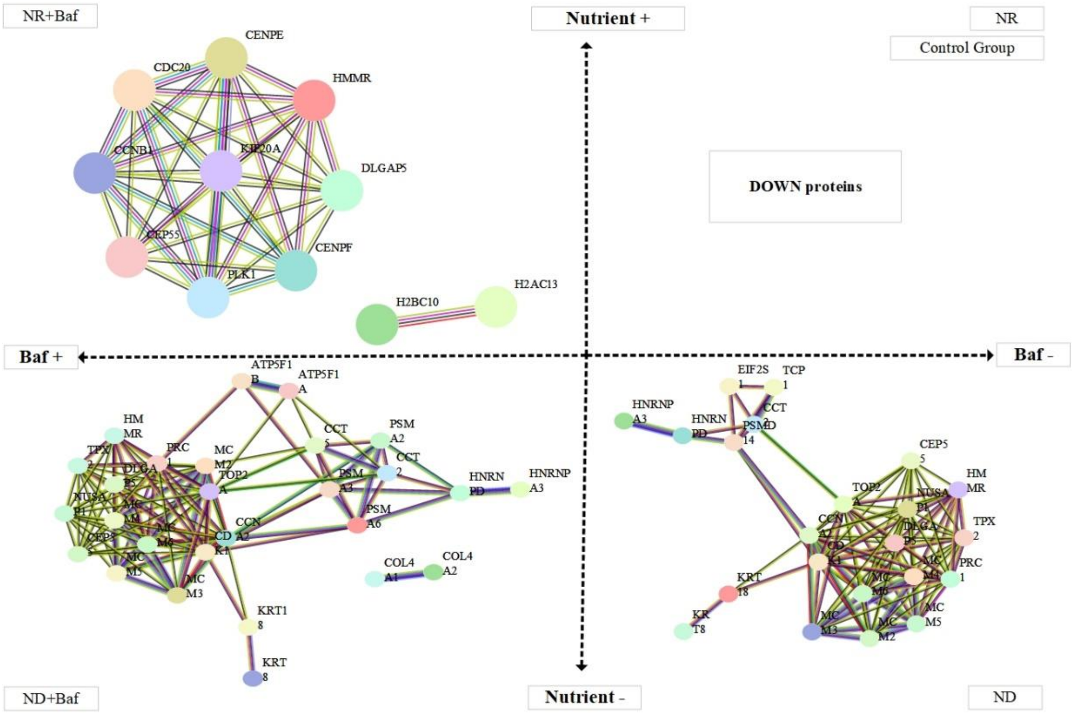


**Figure S6. Quadrant-based PPI network of downregulated core proteins under different conditions of nutrient availability and lysosomal acidity.** Core proteins were extracted based on differential expression and network centrality measures (Table S3). The quadrant layout represents nutrient availability (vertical axis) and lysosomal inhibition (horizontal axis), separating conditions as NR, NR+Baf, ND, and ND+Baf.

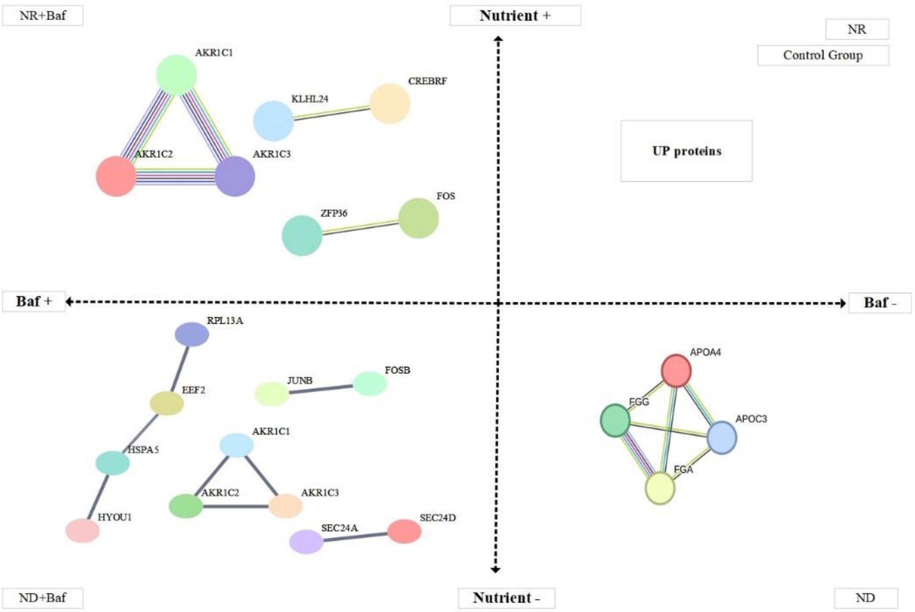


**Figure S7. Quadrant-based PPI network of upregulated core proteins under different nutrient and lysosomal acidity conditions.** Downregulated core proteins were determined by integrating differential expression (adjusted $p < 0.01$) with interaction topology.

#### 5. Regulated cell death (RCD) analyses

The expression of RCD related genes was compared across the four different experimental groups (NR, NR+Baf, ND and ND+Baf). Among the types of RCD programs, the percentage of upregulated genes for NR vs ND+Baf was as follows: 17.78% in the apoptosis gene set, 31.25% in the necrosis gene set, 18.52% in the pyroptosis gene set, and 25% in the ferroptosis gene set (Table S4).

**Table S4. Percentage of upregulated genes for four different types of regulated cell death phenomenon**

| | **NR vs ND** | **NR vs NR+Baf** | **ND vs ND+Baf** | **NR vs ND+Baf** |
|---|---|---|---|---|
| **Apoptosis**<br>**90 genes** | Up: 2<br>Down: 4<br>All: 6 | Up: 2<br>Down: 0<br>All: 2 | Up: 1<br>Down: 5<br>All: 6 | Up: 8<br>Down: 8<br>All: 16 |
| **Necrosis**<br>**16 genes** | Up: 1<br>Down: 3<br>All: 4 | Up: 1<br>Down: 0<br>All: 1 | Up: 0<br>Down: 0<br>All: 0 | Up: 2<br>Down: 3<br>All: 5 |
| **Pyroptosis**<br>**27 genes** | Up: 4<br>Down: 2<br>All: 6 | Up: 0<br>Down: 1<br>All: 1 | Up: 0<br>Down: 0<br>All: 0 | Up: 4<br>Down: 1<br>All: 5 |
| **Ferroptosis**<br>**64 genes** | Up: 7<br>Down: 5<br>All: 12 | Up: 4<br>Down: 1<br>All: 5 | Up: 2<br>Down: 3<br>All: 5 | Up: 12<br>Down: 4<br>All: 16 |

To capture the coordinated response of RCD-related pathways, RCD scores was defined as a metric integrating multiple RCD-associated genes, allowing quantitative comparison across conditions and cell states (Figure S8).

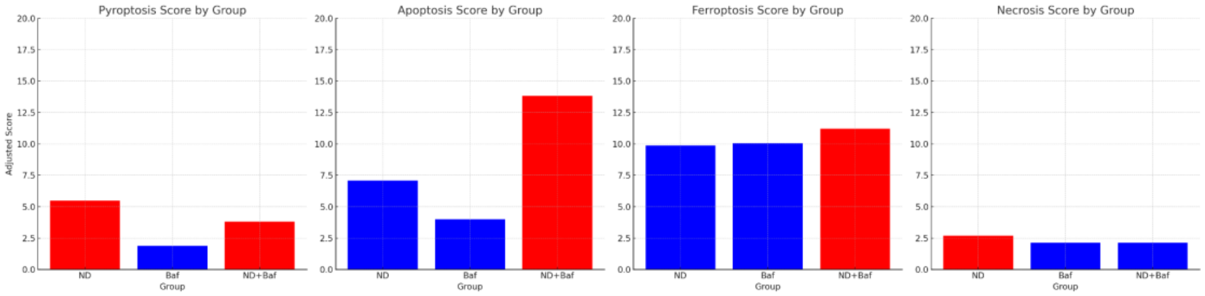


**Figure S8. Comparative analysis of cell death pathway scores across the different experimental conditions (ND, NR+Baf, and ND+Baf).** Pyroptosis and necrosis scores were decreased in the ND+Baf cells compared to ND, while apoptosis and ferroptosis scores were significantly elevated, suggesting condition-specific regulation of cell death pathways.

Significantly upregulated genes associated with apoptosis, necrosis, pyroptosis, and ferroptosis in NR *vs* ND+Baf conditions formed a highly connected PPI network (Figure S9).

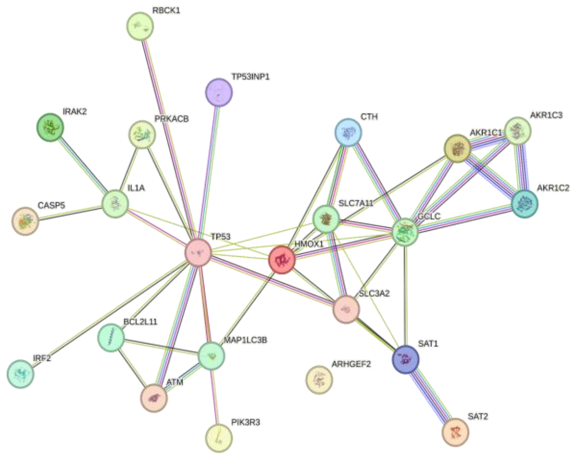


**Figure S9. RCD proteins in ND+Baf.** STRING analysis showing close collaboration of the upregulated RCD proteins under conditions of nutrient depletion and lysosomal alkalinization.

#### 6. Activation of Apoptosis in the ND+Baf cells

The apoptosis-related genes that were identified among the differentially expressed genes (DEGs) across the different experimental groups are shown in Table S5. To facilitate interpretation, logFC2 values were rounded, and only genes meeting a significance threshold of p_adj < 0.01 were included. Data in the column "Functions in Apoptosis" were taken from Gene Cards (https://www.genecards.org/), references other than Gene Card are noted separately.

**Table S5. Apoptosis related genes that are differentially expressed in the four different experimental groups.**

| Gene Name | Gene ID | Function in Apoptosis | $log_2$ FC | Groups | Up/Down |
|---|---|---|---|---|---|
| BCL2L11 | 10018 | Apoptosis activator / facilitator | 1.6 | NR vs ND | Up |
| TP53INP1 | 94241 | Positive regulation of apoptosis | 1.6 | NR vs ND | Up |
| BIRC3 | 330 | Apoptosis inhibitor | -1.3 | NR vs ND | Down |
| CYCS | 54205 | Initiation of apoptosis | -1.5 | NR vs ND | Down |
| ENDOD1 | 23052 | No definite apoptosis relationship - *endonuclease activity by cytosolic calcium* | -1.4 | NR vs ND | Down |
| TNFRSF10D | 8793 | Inhibitory role in TRAIL (cytotoxic ligand) induced cell apoptosis | -1.0 | NR vs ND | Down |

| | | | | | |
|---|---|---|---|---|---|
| PIK3CD | 5293 | Preventing apoptosis (DOI: 10.1007/s13277-016-5225-5) | 1.4 | NR vs NR+Baf | Up |
| TP53INP1 | 94241 | Positive regulation | 1.4 | NR vs NR+Baf | Up |
| PIK3R3 | 8503 | Preventing apoptosis (DOI: 10.1007/s13277-016-5225-5) | 1.3 | ND vs ND+Baf | Up |
| BCL2 | 596 | Blocks the mitochondria mediated apoptotic death | -1.9 | ND vs ND+Baf | Down |
| BCL2L1 | 598 | Potent inhibitor of mitochondria mediated cell death | -1.2 | ND vs ND+Baf | Down |
| CAPN2 | 842 | Inhibitor (DOI: 10.3892/or.2018.6625), *Calcium Mediated T-Cell Apoptosis* | -1.1 | ND vs ND+Baf | Down |
| PPP3CB | 5532 | Inducer (https://doi.org/10.1016/j.bbrep.2023.101603) *Calcium Mediated T-Cell Apoptosis* | -1.2 | ND vs ND+Baf | Down |
| PRKX | 5613 | Inducer (DOI: 10.1016/j.ydbio.2011.05.673) Apoptotic and calcium mediated signaling pathways (https://rgd.mcw.edu/rgdweb/report/gene/main.html?id=1564076) | -1.0 | ND vs ND+Baf | Down |
| ATM | 472 | Apoptosis inducer | 1.0 | NR vs ND+Baf | Up |
| BCL2L11 | 10018 | Apoptosis activator / facilitator | 1.7 | NR vs ND+Baf | Up |
| IL1A | 3552 | Apoptosis inducer | 1.8 | NR vs ND+Baf | Up |
| IRAK2 | 3656 | ER stress induced apoptosis (https://doi.org/10.1371/journal.pone.0064256) | 1.1 | NR vs ND+Baf | Up |
| PIK3R3 | 8503 | Preventing apoptosis (DOI: 10.1007/s13277-016-5225-5) | 1.9 | NR vs ND+Baf | Up |
| PRKACB | 5567 | Inducer (doi: 10.3892/ol.2013.1294 )*14-3-3 Induced Apoptosis* | 1.3 | NR vs ND+Baf | Up |
| TP53 | 7157 | Apoptosis inducer | 1.1 | NR vs ND+Baf | Up |
| TP53INP1 | 94241 | Positive regulation of | 1.7 | NR vs ND+Baf | Up |
| AKT1 | 207 | Preventing apoptosis | -1.1 | NR vs ND+Baf | Down |
| BCL2 | 596 | Blocks the mitochondria mediated apoptotic death | -1.5 | NR vs ND+Baf | Down |
| BCL2L1 | 598 | Potent inhibitor of mitochondria mediated cell death | -1.6 | NR vs ND+Baf | Down |
| BIRC3 | 330 | Apoptosis inhibitor | -1.1 | NR vs ND+Baf | Down |

| | | | | | |
|---|---|---|---|---|---|
| CAPN2 | 824 | *Calcium Mediated T-Cell Apoptosis* | -1.2 | NR vs ND+Baf | Down |
| ENDOD1 | 23052 | No definite apoptosis relationship - *endonuclease activity by cytosolic calcium* | -1.9 | NR vs ND+Baf | Down |
| PPP3CB | 5532 | *Calcium Mediated T-Cell Apoptosis* | -1.6 | NR vs ND+Baf | Down |
| PRKX | 5613 | Apoptotic and calcium mediated signaling pathways (https://rgd.mcw.edu/rgdweb/report/gene/main.html?id=1564076) | -1.7 | NR vs ND+Baf | Down |

We next annotated which apoptosis related genes were up regulated in which condition (Figure S10). Gene names preceded by (-) are apoptosis suppressors and (+) are apoptosis inducer genes. Based on the distribution of the apoptosis score, the most apoptotic experimental group was found to be the ND+Baf cells.

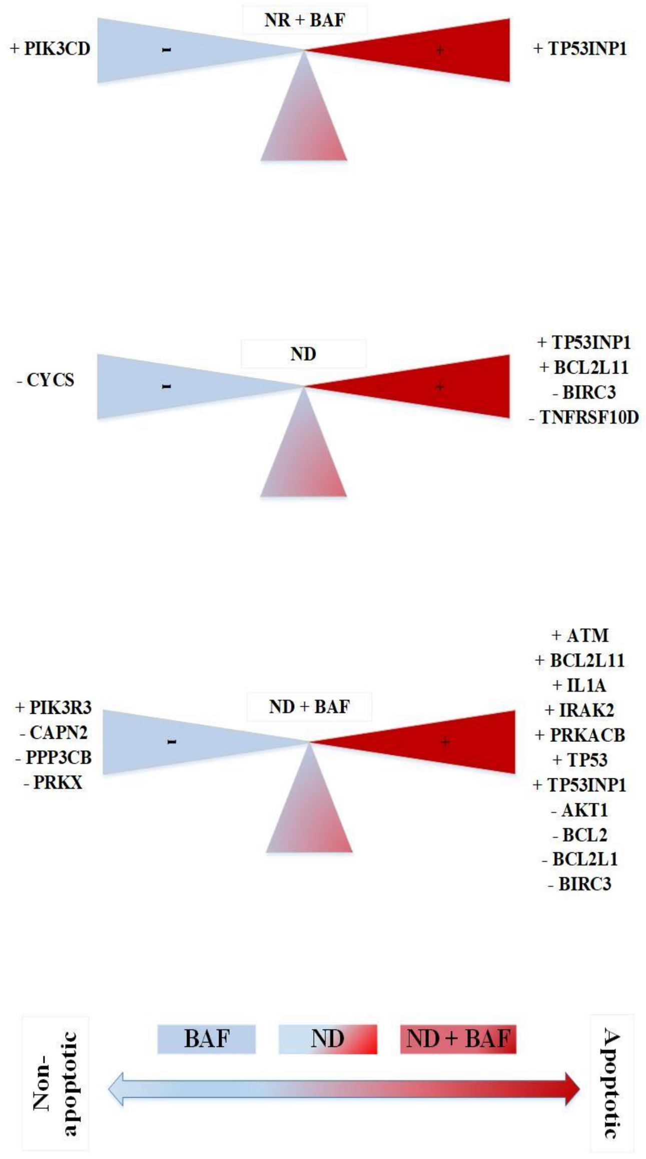


**Figure S10. The logic of apoptosis scoring.** Distribution of significantly altered apoptosis genes in experimental groups (Baf, ND and ND+Baf; NR as control) shows variations.

We next evaluated whether the combination of nutrient depletion and lysosomal alkalinization acted in an additive or synergistic manner to regulate the expression of apoptosis-related genes (Table S6). Genes showing significant differential expression (adj. $p < 0.01$) were categorized based on interaction type (synergy or additive), trend for regulation (increase or decrease), and direction of overall expression (up or down). Missing values indicate non-significant changes under the corresponding condition. Full dataset and detailed scoring logic are available in the GitHub repository (https://github.com/lnnehri/ND-Baf/).

**Table S6. Synergistic and additive response profiles of apoptosis-related genes under ND, Baf, and ND+Baf conditions.**

| ID | Gene Symbol | ND ($log_2FC$) | Baf ($log_2FC$) | ND+Baf ($log_2FC$) | Interaction Type | Regulation | Expression Trend |
|---|---|---|---|---|---|---|---|
| 10018 | **BCL2L11** | 1.64 | 0.945 | 1.71 | Additive | Increase | Up |
| 94241 | **TP53INP1** | 1.61 | 1.42 | 1.75 | Synergy | Decrease | Up |
| 5293 | **PIK3CD** | — | 1.41 | — | Synergy | Decrease | Up |
| 8503 | **PIK3R3** | 1.95 | — | 1.91 | Additive | Decrease | Up |
| 472 | **ATM** | — | — | 1.02 | Synergy | Increase | Up |
| 3552 | **IL1A** | — | — | 1.84 | Synergy | Increase | Up |
| 3656 | **IRAK2** | — | — | 1.11 | Synergy | Increase | Up |
| 5567 | **PRKACB** | — | — | 1.28 | Synergy | Increase | Up |
| 7157 | **TP53** | — | — | 1.18 | Synergy | Increase | Up |
| 596 | **BCL2** | — | — | -1.50 | Synergy | Decrease | Down |
| 598 | **BCL2L1** | — | — | -1.62 | Synergy | Decrease | Down |
| 824 | **CAPN2** | — | — | -1.19 | Synergy | Decrease | Down |
| 5532 | **PPP3CB** | — | — | -1.60 | Synergy | Decrease | Down |
| 5613 | **PRKX** | — | — | -1.74 | Synergy | Decrease | Down |
| 207 | **AKT1** | — | — | -1.13 | Synergy | Decrease | Down |
| 330 | **BIRC3** | -1.29 | — | -1.08 | Synergy | Increase | Down |
| 54205 | **CYCS** | -1.54 | — | — | Synergy | Increase | Down |
| 8793 | **TNFRSF10D** | -1.05 | — | — | Synergy | Increase | Down |
| 23052 | **ENDOD1** | -1.36 | -0.852 | -1.89 | Synergy | Decrease | Down |

Except PIK3R3 and BCL2L11, all detected apoptosis genes exhibited a synergistic effect in the ND+Baf condition, contrasting with their individual behavior in the ND and NR+Baf conditions (https://github.com/lnnehri/ND-Baf). Among the upregulated apoptosis genes demonstrating a synergistic effect, ATM, IL1A, IRAK2, PRKACB, and TP53 showed a synergistic effect resulting in an increase in RNA-seq expression values in the ND+Baf condition, whereas TP53INP1 and PIK3CD exhibited a synergistic effect leading to a decrease. Among the downregulated apoptosis genes, BCL2, BCL2L1, CAPN2, PPP3CB, PRKX, and AKT1 showed a synergistic effect resulting in a further decrease in RNA-seq expression values in the ND+Baf condition compared to ND and NR+Baf alone. Conversely, BIRC3, CYCS, and TNFRSF10D, which were also downregulated, exhibited a synergistic effect leading to an increase in expression in the ND+Baf condition.

Among the genes identified in the ND+Baf cells, ATM, PIK3R3, TP53, TP53INP1, AKT1, BCL2, and BCL2L1 are associated with both EMT and apoptosis. Among these, AKT1, BCL2, BCL2L1 were downregulated (logFC2<-1) and ATM, PIK3R3, TP53, TP53INP1 were upregulated (logFC2>1) (p_adj<0.01). The combination of both up and down regulated genes in this set shows a highly connected and dense PPI network (Figure S11). Moreover, only upregulated proteins and only down-regulated proteins also showed a connected PPI network.

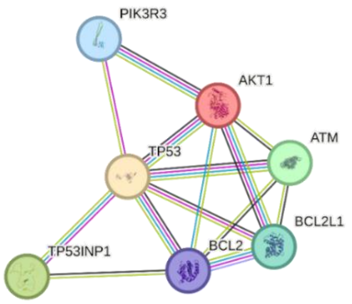


**Figure S11. Protein-protein interaction of the significantly altered genes that have function in both EMT and apoptosis in ND+Baf.**

The GO terms for Figure S11 network were as follows: negative regulation of long-chain fatty acid import across plasma membrane (raw P value=1.36E-03 and FDR value=5.03E-02), negative regulation of reactive oxygen species metabolic process (raw P value=9.31E-05 and FDR value=7.51E-03) and negative regulation of cellular pH reduction (raw P value=3.40E-04 and FDR value=1.93E-02) and oxidative stress-induced premature senescence (raw P value=1.36E-03 and FDR value=5.10E-02). In the ND group, only TP53INP1 was detected to share both apoptosis and EMT characteristics and was upregulated in ND (p_adj<0.01). This suggests that apoptosis genes that are also functional in the EMT process were differentially regulated in ND+Baf, but not in ND alone.

#### 7. DepMap analyses of apoptosis genes

The STRING platform does not incorporate disease-specific conditions by default. Rather, it provides a generalized, context-independent perspective on PPI networks. To understand how these proteins function in cancer-specific, phenotype-specific, or cell-line-specific context regarding the impact of the apoptosis-related proteins we identified in the EMT spectrum we utilized context-specific dependency data available on the DepMap portal (https://depmap.org/portal/data_explorer_2/). We used the "Compare EMT-high to EMT-low adherent models" feature to identify mesenchymal-specific dependencies. CRISPR dependency scores were compared between conditions (EMT-high vs. EMT-low), (Figure S12), while all other parameters were left at their default settings.

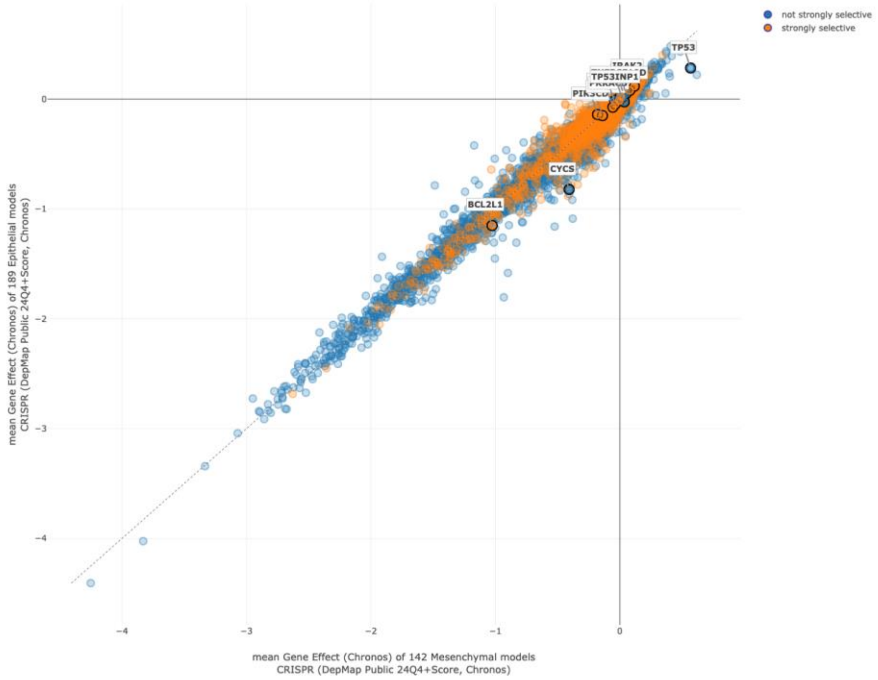


**Figure S12. CRISPR dependency scores of apoptosis genes of ND+Baf in comparison between conditions (EMT-high vs. EMT-low).** CRISPR dependency scores represent how essential a gene is for the survival or proliferation of a cell line, based on CRISPR-Cas9 knockout screens. A lower (more negative) score indicates that the gene is critical for cell viability, its loss significantly impairs growth, whereas scores closer to zero suggest that the gene is non-essential. In the context of EMT-high vs. EMT-low conditions, comparing these scores revealed that apoptosis genes were more critical in the mesenchymal state (DepMap, 2024).

Next, expression levels were plotted on the x- and y-axes, and a linear regression line was added to highlight the correlation (Figure S13); again, all other analysis parameters were maintained as default. The apoptosis-related genes were found to be strongly selective in distinguishing epithelial from mesenchymal phenotypes, further supporting their relevance in EMT-related cellular transitions.

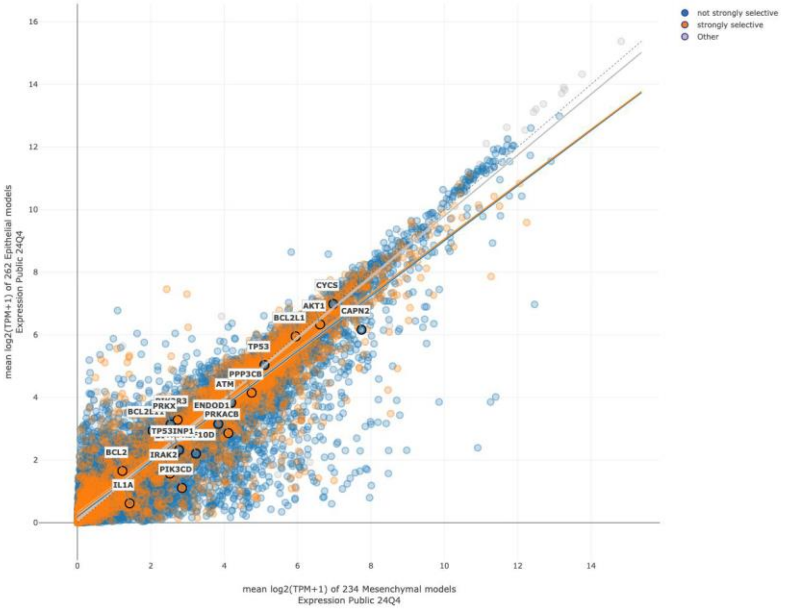


**Figure S13. Gene expressions of apoptosis genes of ND+Baf in comparison between conditions (EMT-high vs. EMT-low).**

#### 8. Validation of findings from independent data sets

***Validation using scRNA-seq data:***

Gene ranking based on principal components (PCs) derived from single-cell data is presented in Figure S14A. This analysis identified the most significant genes contributing to the observed variance in the dataset, highlighting key players in cellular processes relevant to the single cell study (Figure S14B). By considering these hot spot genes, we further analyzed the results with the apoptosis related genes.

**A**

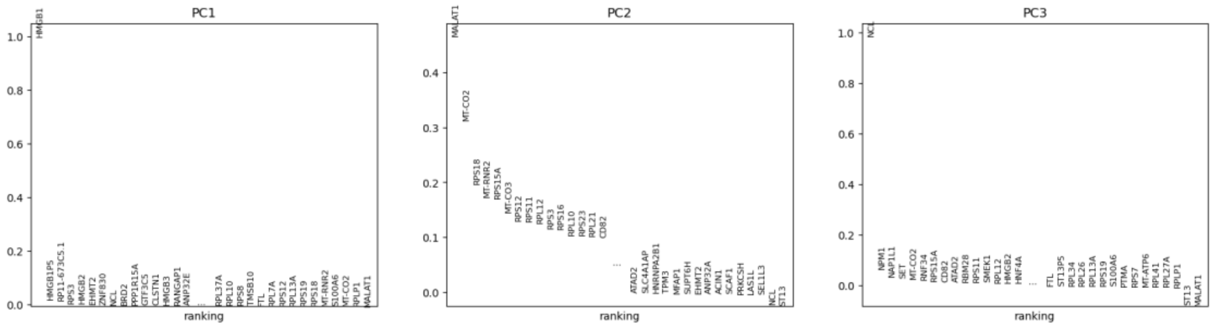

B

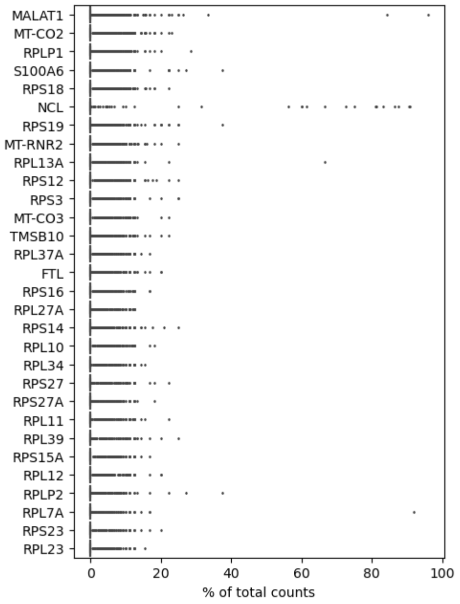


**Figure S14. Analysis of single cell RNA seq data (PRJNA611719) between metastatic and non-metastatic cells.** A. Gene ranking based on principal components. B. List of most highly expressed genes in all analyzed cells.

To evaluate the relationship between the differentially expressed genes between metastatic and non-metastatic cells [2] and the apoptosis related genes identified from GSE245402 [1], the genes were subjected together to a STRING analysis and a highly enriched PPI connection was obtained. One of the most well-connected clusters we observed was composed primarily of ribosomal proteins. Since overly-abundant rRNA transcripts may obscure other low abundance transcripts of interest, we removed all ribosomal proteins from the network. A new PPI network was constructed using the following proteins: BCL2L11, TP53INP1, BIRC3, CYCS, TNFRSF10D, PIK3CD, PIK3R3, BCL2, BCL2L1, CAPN2, PPP3CB, PRKX, ATM, IL1A, IRAK2, PRKACB, TP53, AKT1, NCL, MT-CO3, MT-CO2, MTRNR2L12, FTL, TMSB10, and S100A6. The resulting network was also highly connected (Figure S15).

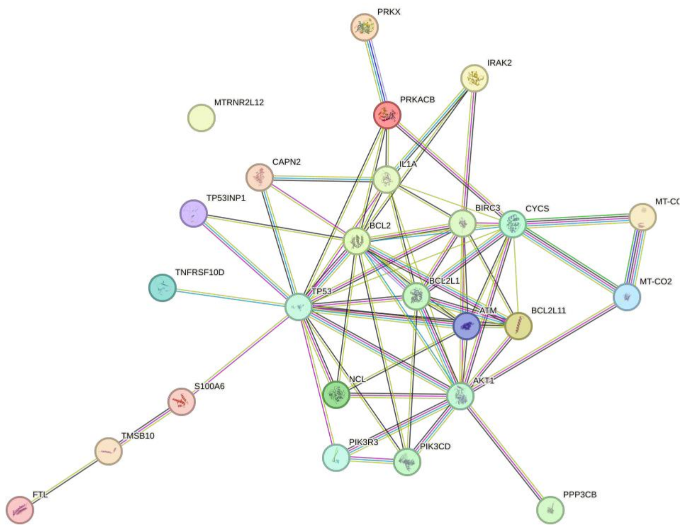


**Figure S15. The PPI network of the most highly expressed genes in all analyzed cells in the PRJNA611719 and apoptosis genes from NR vs ND+Baf GSE245402 after removal of ribosomal proteins.**

The GO analysis of these proteins (Figure S15) revealed significant biological enrichment in the following terms: mitochondrial electron transport, cytochrome c to oxygen (raw p value = 2.40E-06, FDR = 6.74E-04), negative regulation of extrinsic apoptotic signaling pathway in absence of ligand (raw p value = 8.64E-08, FDR = 7.14E-05), positive regulation of release of cytochrome c from mitochondria (raw p value = 2.93E-04, FDR = 2.06E-02), positive regulation of mitochondrial membrane permeability (raw p value = 4.52E-04, FDR = 2.86E-02), negative regulation of intrinsic apoptotic signaling pathway in response to DNA damage (raw p value = 6.04E-04, FDR = 3.55E-02), regulation of endoplasmic reticulum stress-induced intrinsic apoptotic signaling pathway (raw p value = 8.22E-04, FDR = 4.28E-02), and positive regulation of cell migration (raw p value = 4.37E-05, FDR = 5.42E-03). For cellular component enrichment, the enriched GO terms included mitochondrial membrane (raw p value = 2.87E-04, FDR = 4.78E-02), phosphatidylinositol 3-kinase complex, class IA (raw p value = 5.07E-05, FDR = 2.03E-02), and Bcl-2 family protein complex (raw p value = 1.89E-07, FDR = 3.77E-04). Among the enriched molecular function GO terms were BH3 domain binding (raw p value = 2.12E-05, FDR = 1.80E-02) and cytochrome-c oxidase activity (raw p value = 1.10E-04, FDR = 4.30E-02).

**9. Validation of data using an Independent Clinical Proteomic Cohort:**

To further validate our findings at the proteome level, we conducted a focused re-analysis of the proteogenomic data from Tanaka et al. [3], via PyCharm. We utilized the clinical and molecular dataset provided in Table S6 of the manuscript: Clinical attributes of cohort 1 and 2, related to Figures 1–7, encompassing 258 patient samples across primary and metastatic colorectal cancer cohorts [3].

We segregated the samples into two distinct groups based on their classification as primary or metastatic tumors and subsequently identified stably expressed genes within each group using a two-step filtering strategy. First, for each gene, we computed the standard deviation (std_dev) of expression values across all samples within the respective cohort and retained genes with a standard deviation less than 2.5, indicating low variability. Second, we removed genes exhibiting expression outliers based on the interquartile range (IQR) method—specifically excluding those with values below Q1 - 1.5×IQR or above Q3 + 1.5×IQR.

Following this filtration, the resulting datasets contained the following: 212 primary-specific stable genes, 66 metastatic-specific stable genes, and 71 stable genes shared between both cohorts. Next, we focused on secondary (metastatic)-specific stable genes, conducting functional analyses to uncover their biological relevance. We input this gene set into the STRING database for protein-protein interaction analysis, applying default k-means clustering. One prominent cluster (highlighted in red in the STRING output) included the following genes: GADD45GIP1, HSD3B7, LARP7, MRPL22, MRPL58, POLDIP3, PRCC, TM7SF2, XAB2. GO enrichment analysis of this cluster revealed significant associations with: Organelle inner membrane (Cellular Component; raw p = 6.13E-05, FDR = 1.36E-02) and Mitochondrial large ribosomal subunit (Cellular Component; raw p = 1.50E-06, FDR = 1.50E-03). These genes were primarily linked to biological functions such as RNA splicing, cholesterol biosynthesis, and mitochondrial enrichment. Further enrichment analysis using the Enrichr platform (https://maayanlab.cloud/Enrichr/) confirmed pathway-level associations. Key enriched pathways included: Mitochondrial translation and gene expression, Lipid metabolism-related processes, including cholesterol and steroid biosynthesis. Together, these results highlight a mitochondrial and lipid metabolic signature specific to the metastatic CRC cohort, pointing to potential functional mechanisms underpinning metastatic progression.

**10. Coordinated activation of metabolic and stress response pathways.**

Mitochondrial proteins were also incorporated because fatty acid oxidation (FAO), ROS regulation, and apoptosis converge at this organelle. As the central hub for oxidative metabolism and redox balance, mitochondria integrate lipid metabolism and stress signaling, making them essential for interpreting coordinated pathway activation. Therefore, we evaluated the mitochondrial proteins and their relations with apoptotic proteins in ND+Baf condition (Figure S16).

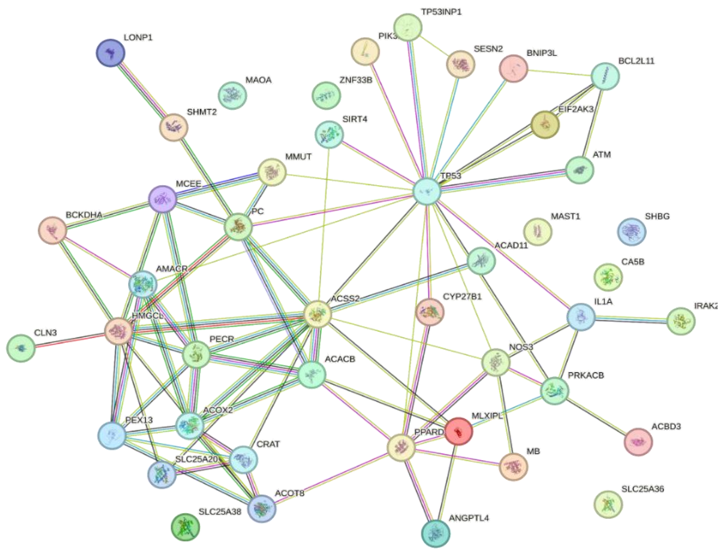


**Figure S16. PPI network of significantly altered apoptosis-, fatty acid synthesis-, fatty acid oxidation-, and mitochondria-related genes in the ND+Baf condition.** The connected network highlights the functional relationships among these pathways, illustrating their coordinated regulation under lysosomal alkalinization and nutrient-deprived conditions.

To assess whether the metabolic and stress-response pathway was activated in a coordinated manner under both nutrient and lysosomal acidity perturbations, pathway scores were calculated and compared across the different experimental groups (Figure S17). These scores were calculated by aggregating normalized inducer and suppressor gene expression values for fatty acid oxidation (FAO, orange) and reactive oxygen species regulation (ROS, blue). Each bar represents the overall activity score per condition (Groups 2: ND, 3: NR+Baf, and 4: ND+Baf), normalized to the dynamic range of the dataset. The results illustrate coordinated regulation between lipid metabolism and oxidative stress response under nutrient depletion and lysosomal alkalinization.

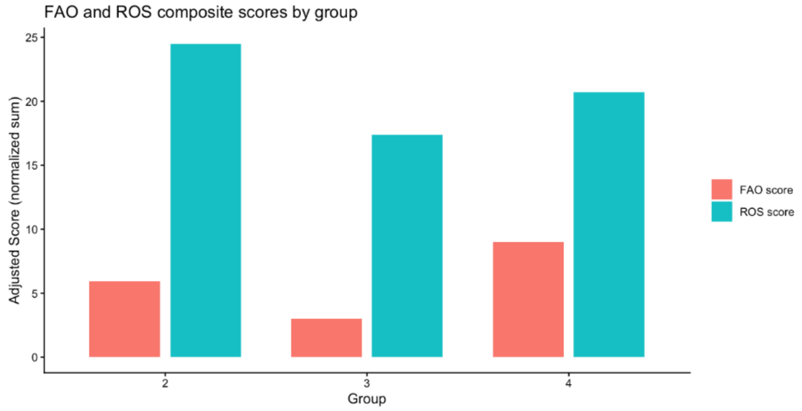


**Figure S17. Integrated comparison of FAO and ROS scores across the different experimental groups (ND, NR+Baf and ND+Baf for Groups 1, 2 and 3, respectively).**

#### 11. Analysis of Calcium mediated signaling

A synergy analysis was carried out with the DEGs in the 4 experimental groups (NR, ND, NR+Baf, ND+Baf) to evaluate whether there is a synergistic effect in the genes that were most highly upregulated (logFC2>4 and p_adj<0.01) and their relation to calcium metabolism. The genes were as follows: ANK1, ARFGEF3, ASPA, BIRC7, C19orf38, DNAJC12, FGF19, LAMP3, LINC03099, MB, MKX, NPHS1, PERCC1, PLIN5, SLC8A1, SSTR3, TCP11L2, VAV3, YPEL2, ARHGAP9, ATCAY, ATF6-DT, ATP13A4, C11orf86, C11orf96, C2CD4B, CCDC149, CHRM1, COLQ, CTNND2, DACT1, DSCAML1, FOLR2, GFY, HIPK4, HOXD1, HRK, IGHG4, INHBE, KLHDC7B, LINC00955, LINC01847, LRRC2, MATN1, MRPL23-AS1, MYOCD, MYRIP, NIBAN1, NKX6-1, NWD2, PIERCE2, PIEZO2, RAB39B, RAB3C, SDK2, SNPH, TCAF2, TRIM50, TRPM8.

The synergistic proteins were subjected to STRING analysis, and subjected to MCL clustering (Inflation parameter=3), following which the main nodes of connected proteins were identified (Figure S18). Five proteins were identified. TRPM8 is a non-selective transient receptor potential (TRP) channel that is $Ca^{2+}$-permeable; PIEZO2 (Piezo Type Mechanosensitive Ion Channel Component 2) is a mechanosensitive monatomic ion channel; TCAF2 (TRPM8 Channel Associated Factor 2), participates in the regulation of anion channel activity and positive regulation of cell migration and is located in the cell junction and plasma membrane. ANK1 (Ankyrin 1) connects integral membrane proteins to the underlying spectrin-actin cytoskeleton and MB (Myoglobin) facilitates the storage and transfer of oxygen from the cell membrane to the mitochondria. These genes highlight a specific role of Ca in cellular motility in the ND+Baf cells.

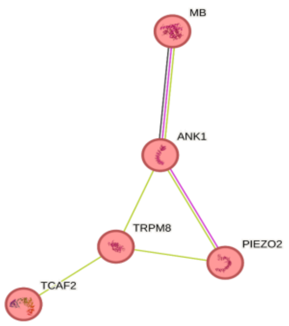


**Figure S18. Synergy among the most highly upregulated genes in ND+Baf highlights calcium signaling as a central convergent pathway.**

To examine how stress-responsive pathways converge at the protein level in ND+Baf cells, we carried out PPI clustering of the significantly upregulated apoptosis-, calcium-, fatty acid–, and mitochondria-associated proteins in the ND+Baf cells (Figure S19).

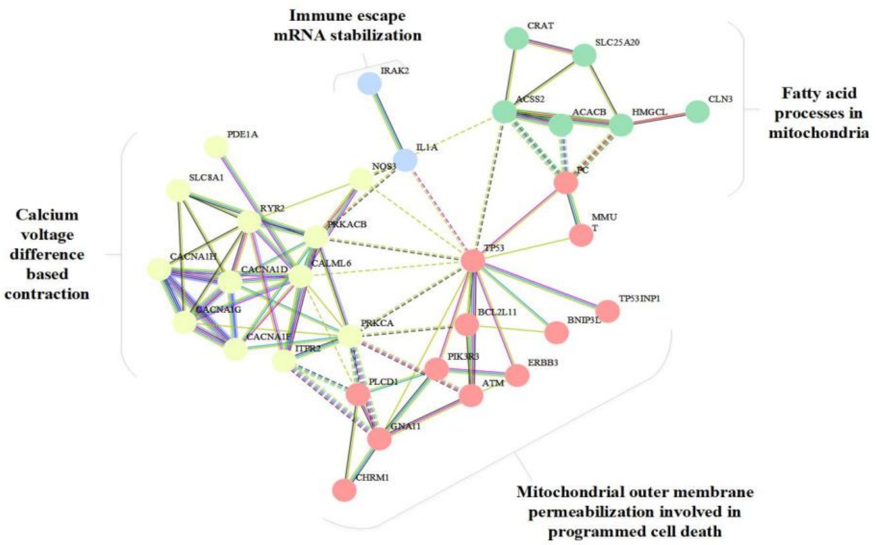


**Figure S19. Protein-protein interaction clustering reveals coordinated coupling of apoptosis, mitochondrial metabolism, fatty acid metabolism and $Ca^{2+}$ signaling in ND+Baf cells.**

We observed that the core proteins associated with apoptosis, $Ca^{2+}$ response, fatty acid metabolism and mitochondrial genes formed a highly connected network in the ND+Baf cells (compared to NR). The clustering of proteins shows that the apoptosis response and fatty acid metabolism was linked via mitochondria, and the mitochondria - apoptosis (red cluster) and mitochondrial - fatty acid (green cluster) clusters were all connected to each other. The yellow cluster is related to calcium and muscle contraction that is caused by voltage difference, and this cluster was related to the mitochondria-apoptosis cluster (red cluster). In addition, the blue cluster (4) formed by IL1A and IRAK2, which contains only 2 proteins, has functions such as inducing escape against the immune response (IL1A) and stabilizing the mRNAs generated for it (IRAK2), and this cluster is involved in apoptosis response in association with TP53.

We next evaluated whether exocytosis was activated in the ND+Baf cells. We identified the following significant GO terms: positive regulation of calcium ion-dependent exocytosis (raw P value=1.89E-07 and FDR value=6.14E-05), SNARE complex assembly (raw P value=5.29E-05 and FDR value=1.02E-02), cellular response to calcium ion (raw P value=9.22E-06 and FDR value=1.29E-02) and protein secretion (raw P value=1.97E-04 and FDR value=2.89E-02).

We observed a clustering of core connected networks of significantly upregulated exocytosis proteins in NR vs ND+BAF (logFC2>1 and p_adj<0.01) (Figure S20). The same proteins were identified to be downregulated in ND+Baf vs NR cells (logFC2<-1 and p_adj<0.01). In the green cluster, TRPV6 (Transient receptor potential cation channel subfamily V member 6), which is a Calcium selective cation channel that mediates $Ca^{2+}$ uptake, was connected to Ankyrin-1 (ANK1), which attaches integral membrane proteins to cytoskeletal elements, and ANK1 was connected to Neurogenic locus notch homolog protein 1 (NOTCH1), which plays a significant role in apoptosis. The NOTCH1 protein was connected to serine/threonine-protein kinase B-raf (BRAF), which activates the MAPK pathway.  In the blue cluster, several well-known exocytosis related Rab proteins were identified. In the red cluster, we identified SNARE complex related proteins, which are critical for exocytosis. At the core of the network were Syntaxin-1A (STX1A), which has an essential role in calcium-dependent exocytosis of hormones and neurotransmitters, and Synaptotagmin-2 (SYT2), which has calcium-dependent phospholipid and inositol polyphosphate binding characteristic. These data suggest that calcium plays a prominent role in exocytosis.

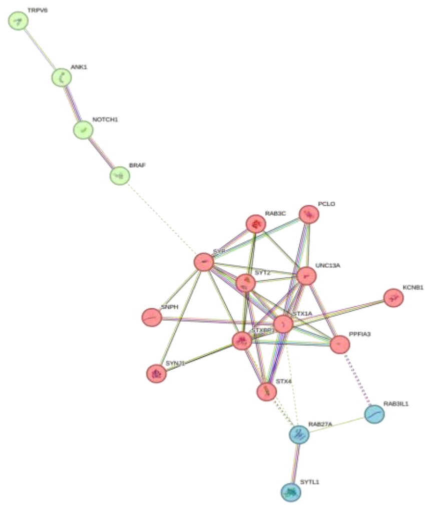


**Figure S20. PPI network showing a critical role of calcium metabolism for cellular exocytosis in the ND+Baf cells.**

To assess how nutrient and lysosomal stress can alter intercellular communication, we examined the clustering of cell-cell adhesion proteins that were significantly downregulated in NR (as control) relative to ND+Baf (as treatment) (Figure S21). The same proteins were upregulated in ND+Baf (as control) and NR (as treatment), (logFC2>1 and p_adj<0.01). The yellow cluster includes Cadherins which are calcium-dependent cell adhesion proteins and green cluster includes Claudins which may be regulated with calcium for cell adhesion. These clusters are connected with the red cluster which includes actin related proteins and regulation of heterotypic cell-cell adhesion.

Overall, these data suggest that rather than cell-cell communication, $Ca^{2+}$ signaling was more relevant to the process of exocytosis in the ND+Baf cells.

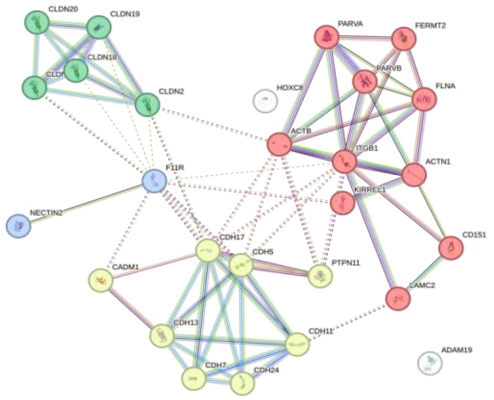


**Figure S21. Clustering of significantly downregulated (logFC2<-1 and p_adj<0.01) cell to cell communication proteins of NR vs ND+BAF.**

To better understand the biological relevance of genes associated with the ND+Baf, a risk-based prioritization approach was employed. Genes with an adjusted p-value (p_adj < 0.01) were first identified using CTpathway analysis (http://www.jianglab.cn/CTpathway). Among these, genes with a risk score greater than 0.8 were selected for further investigation. The risk score reflects the strength of association between each gene and the condition of interest, incorporating information such as pathway involvement and gene expression patterns. Selecting high-risk genes allows researchers to focus on those most likely to contribute to the observed phenotype. To explore the functional context of these high-risk genes, a Gene Ontology (GO) enrichment analysis was conducted. Enrichment analysis was performed for genes with p_adj<0.01 based on the output number 20240611094125368 from CTpathway for the ND+Baf group. The significant GO terms included regulation of response to calcium ion (raw P value=4.86E, FDR value=043.88E-02) and smooth endoplasmic reticulum calcium ion homeostasis (raw P value=4.86E-04, FDR value=3.86E-02). The enrichment analysis revealed that the genes with the highest risk scores in ND+Baf cells were predominantly associated with calcium regulation, in other words, genes involved in calcium regulation were found to have high risk scores, implying a strong association between calcium-related pathways and the cellular effects of ND+Baf treatment.

##### 12. Validation of the role of Calcium Signaling in Cellular Motility using scRNA-seq Data:

To verify whether alterations calcium mediated signaling could contribute to an elongation in shape and EMT characteristics, the DEGs obtained for ND+Baf vs NR were compared with the data reported by Bernal et al. (2020). The results showed that the GO terms related to the common genes that were both downregulated and upregulated indeed included calcium-mediated signaling. Additionally, the main node of these common genes from the STRING network (with MCL clustering = 1.1), included the GO terms regulation of sequestering of calcium ion (raw P value = 2.14E-04 and FDR = 4.52E-02), calcium-mediated signaling (raw P value = 4.21E-05 and FDR = 1.53E-02), and calcium ion homeostasis (raw P value = 1.64E-04, FDR = 3.90E-02).

##### 13. Mathematical modeling to predict phenotypic alterations in Caco-2 cells

**13.1 Phenotype probability:**

We modeled the probability of the acquisition of different phenotypes of Caco-2 cells as Living (L), Elongated (E) and Dead (D). Considering E(t) to denote the number of elongated cells at time t, L(t) the number of dormant (rounded) cells, and D(t) the number of dead cells, the total cell population (N) can be denoted as N=E(t)+L(t)+D(t) (Figure S22). Cells can transition between these states in response to changing conditions, with the rate of transition between states described by the function k. Living cells are capable of lipid metabolism, which dead cells are not; consequently, dead cells were excluded from the heterotypic signaling network of living cells. Transitions between the live and elongated states among the living cells are regulated by intracellular calcium accumulation. In addition, the living cell population is capable of sensing death-associated signals emitted by neighboring dead cells.

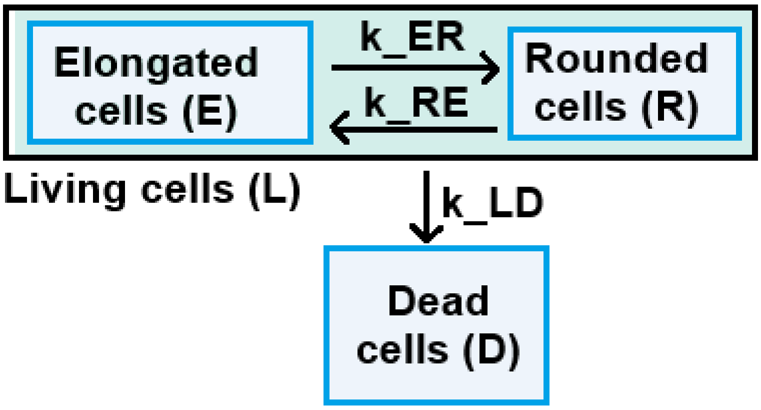


**Figure S22. Schematic diagram to model the different states of Caco-2 cells upon nutrient depletion and lysosomal alkalinization (ND+Baf).** E(t) is the number of elongated cells at time t, L(t) is the number of dormant cells (rounded cells), D(t) is the number of dead cells and N(t)=E(t)+L(t)+D(t) is the total number of cells. The cells may change from one state to another when the conditions change. The function of k shows the rate of the changes between cell types from one state to another.

We next modeled early phenotypic composition of the Caco-2 cells (rounded/living, elongated, dead cells) and time-to-decision dynamics using Poisson- and Gamma-based probability distributions (Figure S23). First, phenotype proportions up to 48 h were estimated with a Poisson model: the death probability was computed as P(death at k=1 | λ_death) with dpois(1, λ_death) where $\lambda$_death was derived from the experimentally observed death counts (e.g., 12/100), and the probability of being alive was 1 − P(death). Among living cells, cell elongation used dpois(1, λ_meta) with $\lambda$_meta = 26 / 126 (elongated cell subpopulation among living cells), and the live fraction was 1 − P(rounded). Joint probabilities (elongated, live, and dead) were verified to sum to 1 and visualized with ggplot2. Next, Gamma distributions were used to link lipid metabolism and calcium mediated signaling to phenotypes (live, elongated, dead). Probability density curves were generated with dgamma (x, shape = a, rate = λ), assigning dead, live, and elongated groups different (shape, rate) pairs derived from the Poisson-based probabilities (e.g., shape = prob_*, rate = prob_*).

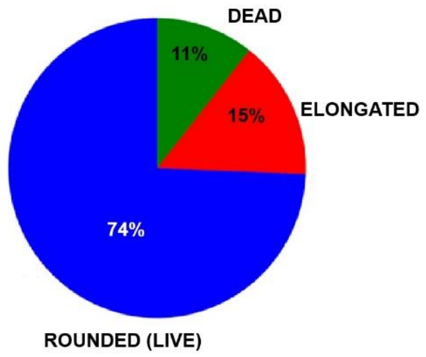


**Figure S23. Caco-2 sub-phenotypes via Poisson distribution**. After 48 hours of nutrient depletion and lysosomal alkalinization of Caco-2 cells, the λ_elongation was 0.21 and λ_dead was 0.12. Since each well of the cell culture plate is a discrete unit and the defined events (such as death or elongation) may happen at any time up to the 48th hour, it was used a single plate cell population to model Poisson distribution of being dead until the 48th hour and being elongated among the living cells until the 48th hour.

**13.2 Categorical comparison of gamma-modeled phenotypes:**

To compare the distributions associated with metastatic, rounded, and death phenotypes, we generated synthetic samples from Gamma distributions parameterized by probabilities estimated in the Poisson stage (Figure S24).

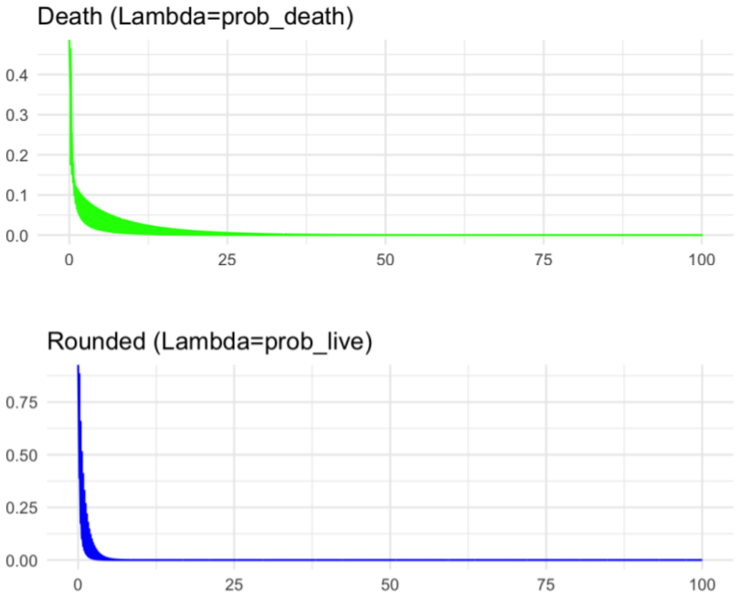

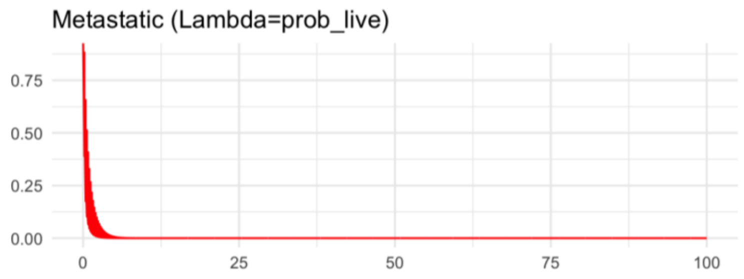


**Figure S24.** Gamma distribution of cell phenotypes derived from assumed parameters inferred through bioinformatic analyses. The parameters, (fatty acid oxidation and calcium release), were assumed to be the key determinants shaping the Gamma distributions. Levene's test results indicated significant differences in variance across phenotypic groups, consistent with the Gamma distribution profiles. Together, these analyses suggest that elongated cells exhibit greater molecular heterogeneity compared to rounded and dead cell populations. In the figure, red = elongated (metastatic), blue = live, and green = dead cells.

Using a fixed seed (set.seed(123)), we drew n = 500 values per group with rgamma: metastatic (shape = prob_live, rate = prob_metastasis), rounded (shape = prob_live, rate = prob_rounded), and death (shape = prob_death, rate = prob_death). Continuous draws were discretized into five bins with equal width (0–10, 10–20, 20–30, 30–40, 40–50) using cut, and observed bin frequencies were computed for each group. For each phenotype, a Chi-square goodness-of-fit test (chisq.test) was applied to assess deviation of the observed bin counts from an equal-probability categorical reference, expected frequencies taken as the column means across groups, normalized to sum to 1 (Figure S25). To evaluate the homogeneity of variances among phenotypic groups (Elongated, Rounded, and Death), a Levene's test was performed. Variance equality was assessed by comparing the absolute deviations of each observation from its group mean. The test statistic and p-value were used to determine whether the null hypothesis of equal variances could be rejected.

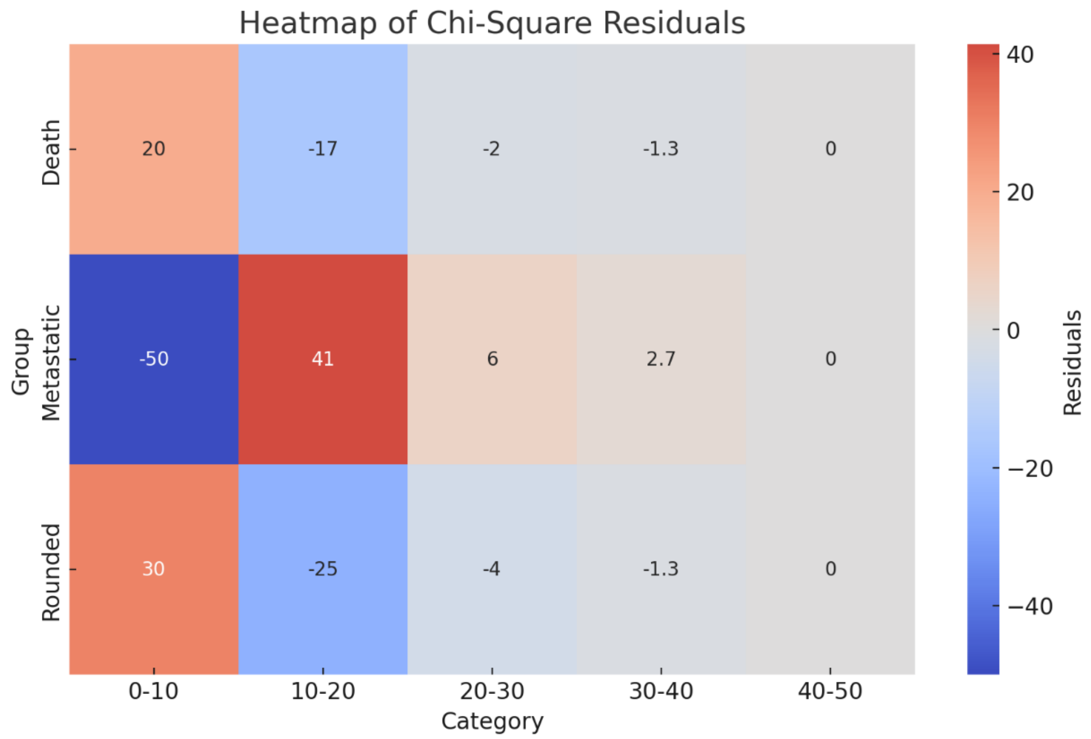


**Figure S25.** Heatmap of Chi-square residuals of the gamma distribution, which indicates that the differences among groups.

##### 13.3 Cumulative modeling:

Modeling the time-to-elongation based on calcium signaling and FAO dynamics predicted that the transition toward an elongated morphology began during the early lifetime of the population, highlighting the temporal coupling between metabolic and signaling processes (Figure S26, Supplementary Methods).

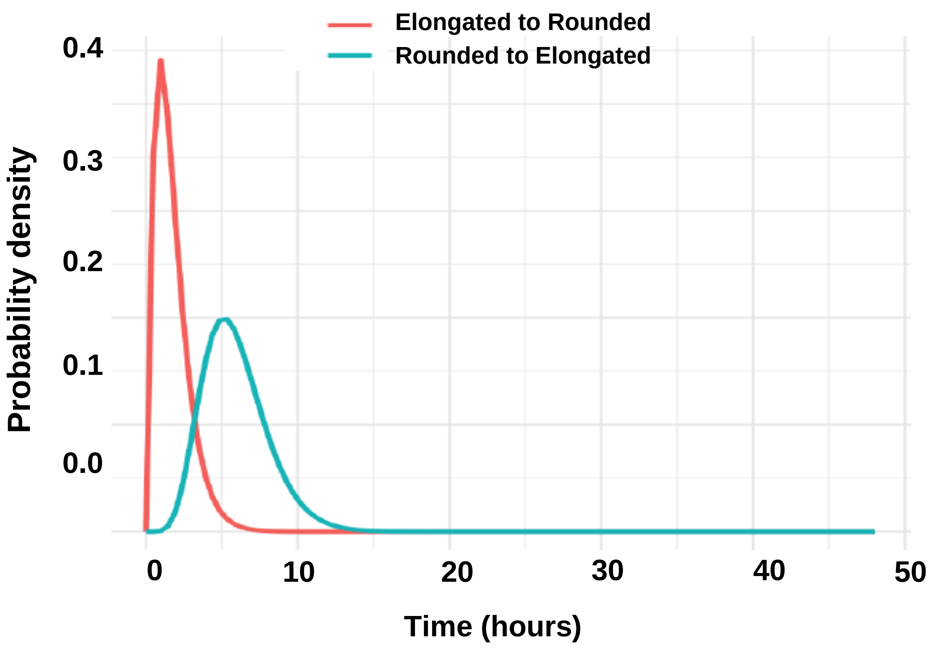

**Figure S26:** Model of the decision point for shape change from rounded to elongated. The transition from a rounded to an elongated shape is driven by a critical decision point just prior to one-fourth of the cell's half-life during nutrient depletion. At this stage, calcium mediated signaling may trigger lysosomal exocytosis while fatty acid oxidation may provide the energy required for survival and morphological transformation.

In the cumulative model, T represents the average decay rate of the cell population, and 1/T corresponds to the mean population lifespan (~48 hours under nutrient depletion and lysosomal alkalinization-ND+Baf). The cumulative probability framework therefore constrains the elongation decision to occur within a fraction of this lifespan. To adopt an elongated phenotype before population decline, cells must commit to shape transition within approximately the first quarter of 1/T. The gamma distribution used to model transition timing was parameterized as follows: the shape parameter (k) was determined by the probability of intracellular calcium accumulation, such that higher calcium probability increased k and thus accelerated the likelihood of transition. The scale parameter (θ) was defined as inversely proportional to the probability of lipid accumulation; lower lipid accumulation increased θ, extending the distribution tail. Consequently, cells exhibiting sustained calcium signaling with relatively limited lipid accumulation were predicted to commit to elongation earlier, consistent with increased outcome probability among viable rounded and elongated subpopulations (Figure S27).

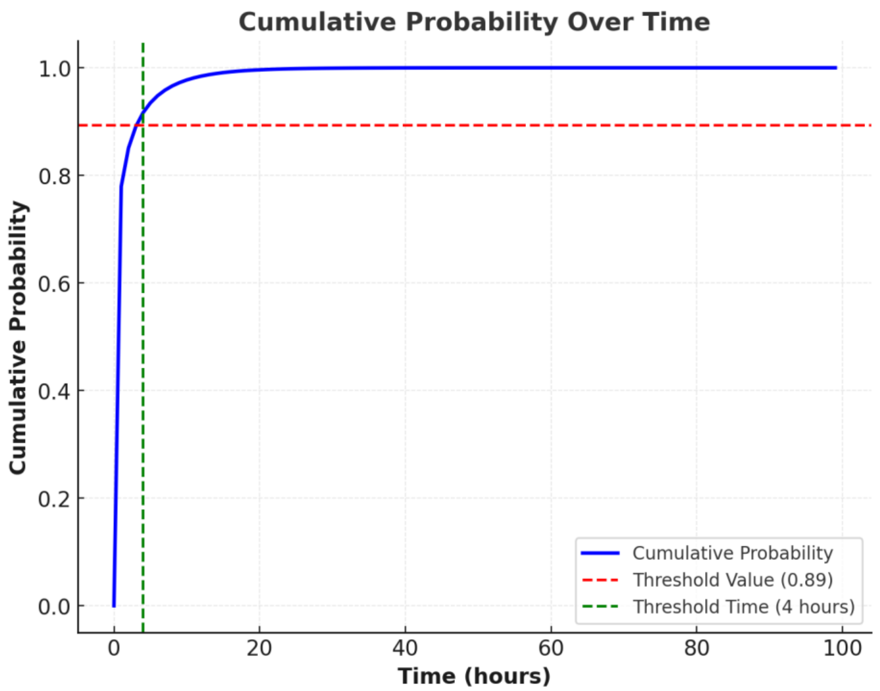


**Figure S27. Decision point to change shape from rounded to elongated shape based on the accumulation of Calcium.** The transition from a rounded to an elongated shape is driven by a critical decision point at before one-fourth of the cell's half-life during starvation. At this stage, calcium accumulation triggers cytoskeletal remodeling and adhesion reorganization, while sufficient FAO provides the energy required for morphological transformation. Together, these factors push cells toward a metastatic phenotype by enabling shape change and increased motility. The figure was created by using R.

##### 13.4 Bayesian modeling:

The Bayesian network summarizes the inferred relationships between nutrient depletion, lysosomal alkalinization, ROS, fatty acid oxidation, calcium signaling, and cell phenotype. In the model (Figure S28), nutrient depletion and lysosomal alkalinization were considered to act as upstream stress inputs that modulate ROS production and fatty acid oxidation, which subsequently influence calcium signaling. These signaling states collectively determine cell phenotype, classified as rounded (live), elongated, or dead.

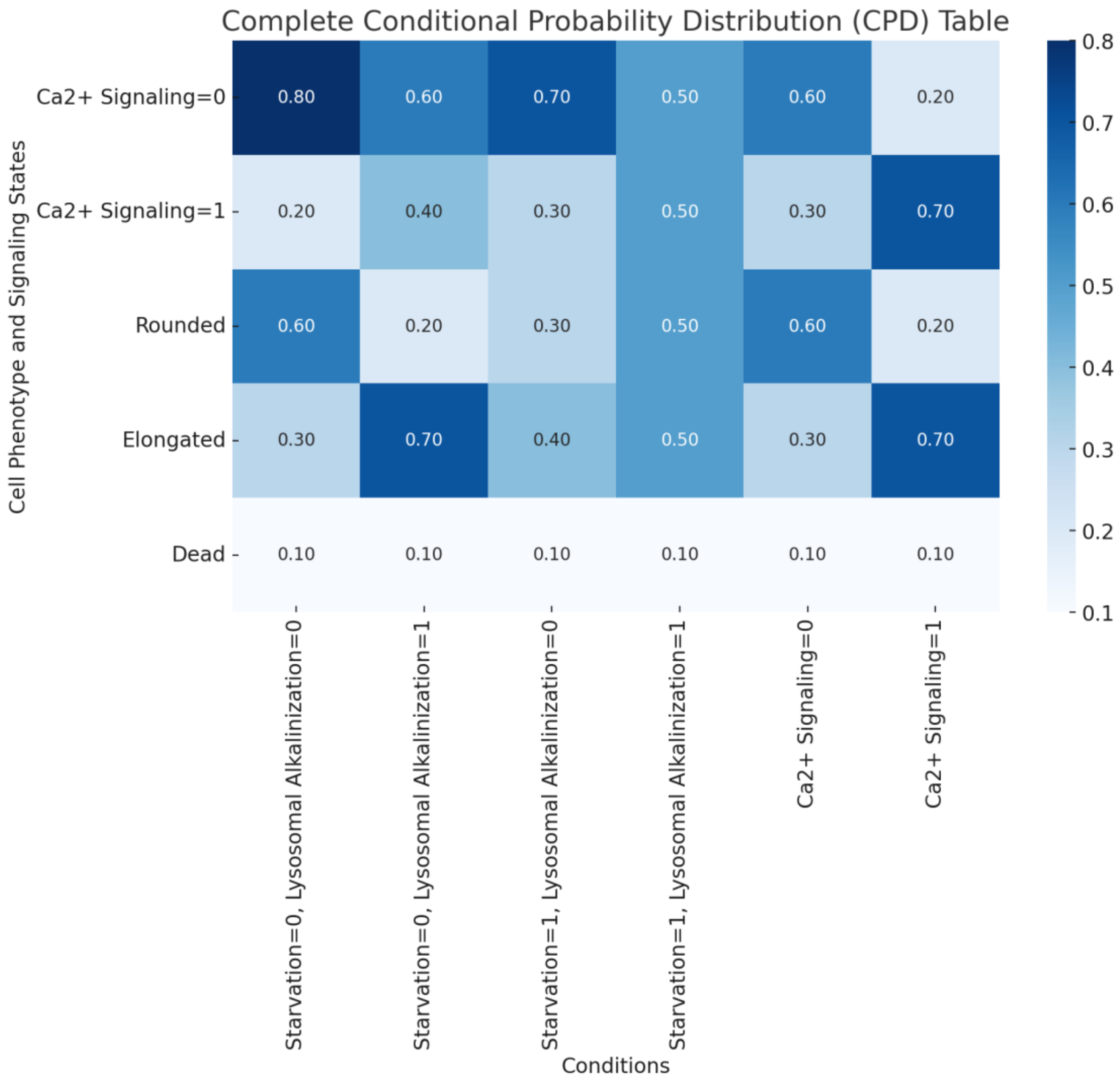


**Figure S28. Conditional probability table of the Bayesian network.**
Heatmap representation of the conditional probability distributions used in the Bayesian network. Columns correspond to combinations of starvation and lysosomal alkalinization states, while rows indicate calcium signaling states and cell phenotypes (rounded, elongated, dead). Values denote the probability assigned to each state under the specified conditions. The CPDs were defined based on

experimentally validated observations and prior bioinformatic analyses, and implemented in Python. Darker color intensities indicate higher probabilities.

The eight evidence combinations used as model inputs were defined as follows:
Combo_0: Starvation = 0, Lysosomal alkalinization = 0, ROS = 0, Lipid oxidation = 0
Combo_1: Starvation = 0, Lysosomal alkalinization = 0, ROS = 0, Lipid oxidation = 1
Combo_2: Starvation = 0, Lysosomal alkalinization = 0, ROS = 1, Lipid oxidation = 0
Combo_3: Starvation = 0, Lysosomal alkalinization = 0, ROS = 1, Lipid oxidation = 1
Combo_4: Starvation = 0, Lysosomal alkalinization = 1, ROS = 0, Lipid oxidation = 0
Combo_5: Starvation = 0, Lysosomal alkalinization = 1, ROS = 0, Lipid oxidation = 1
Combo_6: Starvation = 1, Lysosomal alkalinization = 0, ROS = 0, Lipid oxidation = 0
Combo_7: Starvation = 1, Lysosomal alkalinization = 1, ROS = 1, Lipid oxidation = 1

These combinations were used as evidence nodes to infer phenotype probabilities within the Bayesian network framework (Figure 4B). The conditions and results shown in Figure 4B and 4C represent a simulation based on the prior probabilities assigned in the probability tables presented above (Figure S28), with the outcomes illustrated in Figure 4B.
The conditional probabilities for the Bayes network were as follows:

1. Conditional probability for ROS:

Let $S \in \{0,1\}$denote starvation status (0 = absent, 1 = present). Then:

$$P(ROS = High \mid S) = \begin{cases} \alpha_1 & \text{if } S = 1 \\ \alpha_0 & \text{if } S = 0 \end{cases}$$
$$P(ROS = Low \mid S) = 1 - P(ROS = High \mid S)$$

2. Conditional probability for fatty acid oxidation:

$$P(LipidOx = High \mid S) = \begin{cases} \beta_1 & \text{if } S = 1 \\ \beta_0 & \text{if } S = 0 \end{cases}$$
$$P(LipidOx = Low \mid S) = 1 - P(LipidOx = High \mid S)$$

3. $Ca^{2+}$ signaling conditional probability:

Let $A \in \{0,1\}$denote lysosomal alkalinization (0 = absent, 1 = present):

$$P(Ca^{2+} = Activated \mid A) = \begin{cases} \gamma_1 & \text{if } A = 1 \\ \gamma_0 & \text{if } A = 0 \end{cases}$$
$$P(Ca^{2+} = Not\,Activated \mid A) = 1 - P(Ca^{2+} = Activated \mid A)$$

4. Cell phenotype conditional probability:

Let

- $R \in \{High, Low\}$(ROS)

- $L \in \{High, Low\}$(Lipid oxidation)
- $C \in \{Activated, NotActivated\}$($Ca^{2+}$ signaling)
- $CP \in \{Rounded, Elongated, Dead\}$

Then:

$$P(CP = c \mid R, L, C) = \theta_{c,R,L,C}$$

with normalization constraint:

$$\sum_{c \in \{Rounded, Elongated, Dead\}} \mathrm{P(CP = c \mid R, L, C) = 1}$$

**14. Source files for western blots**

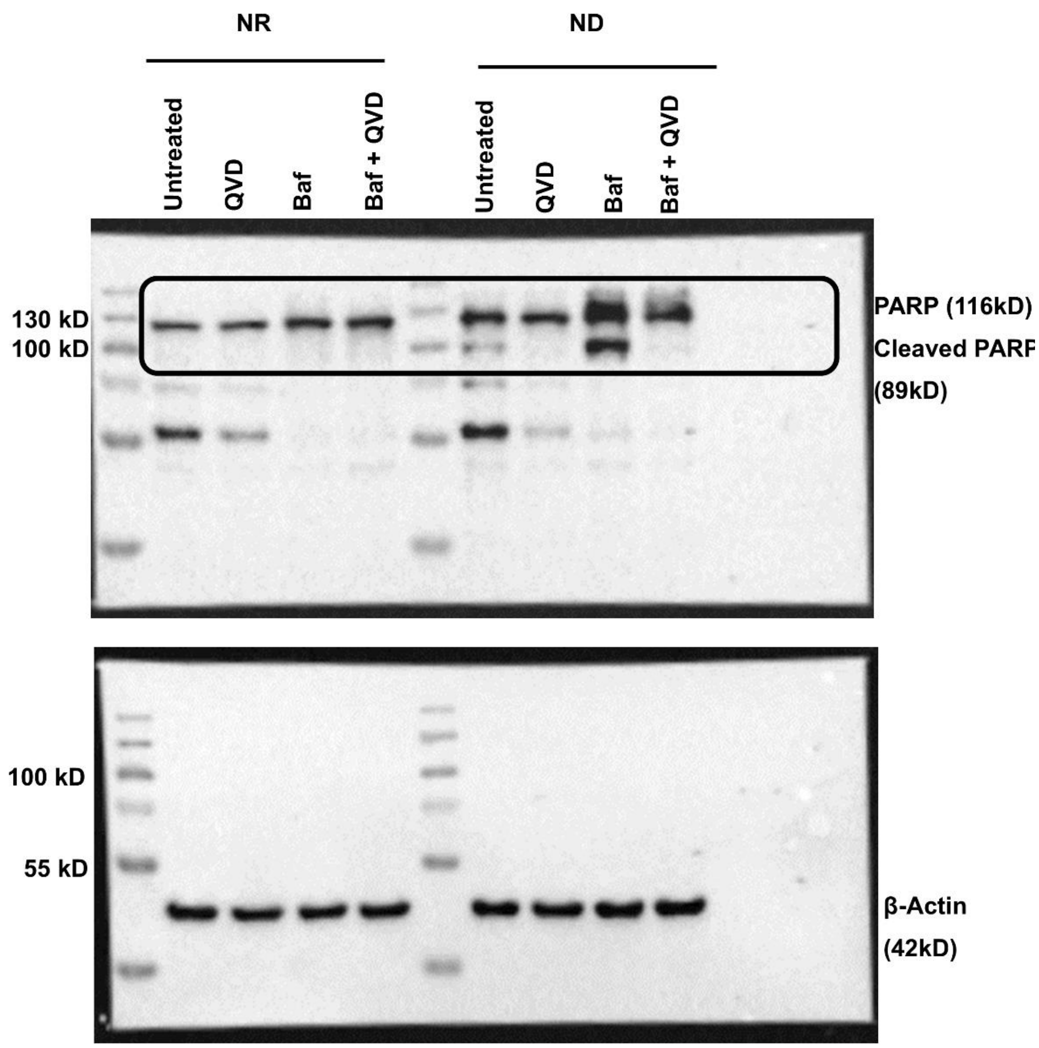
NR
ND
Untreated
QVD
Baf
Baf + QVD
Untreated
QVD
Baf
Baf + QVD
130 kD
100 kD
PARP (116kD)
Cleaved PARP
(89kD)
100 kD
55 kD
β-Actin
(42kD)

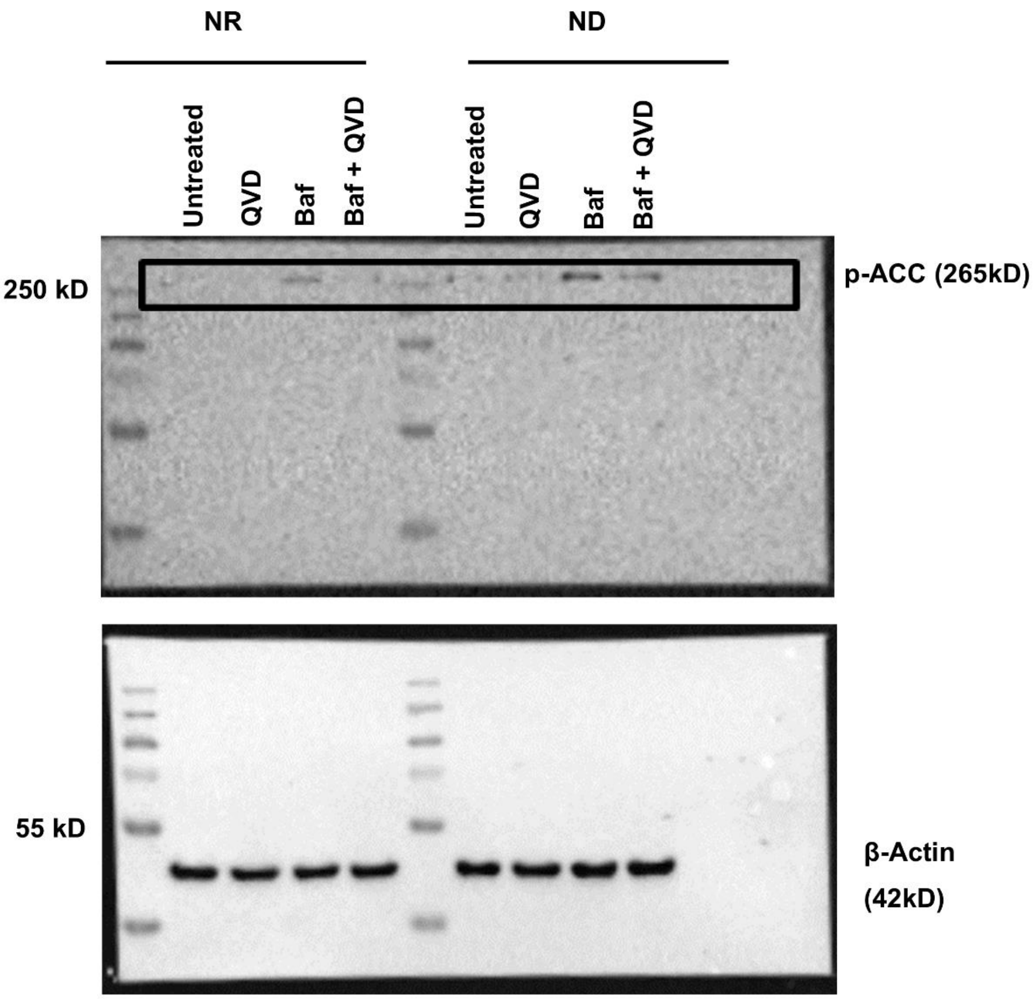
NR
ND
Untreated
QVD
Baf
Baf + QVD
Untreated
QVD
Baf
Baf + QVD
250 kD
p-ACC (265kD)
55 kD
β-Actin
(42kD)

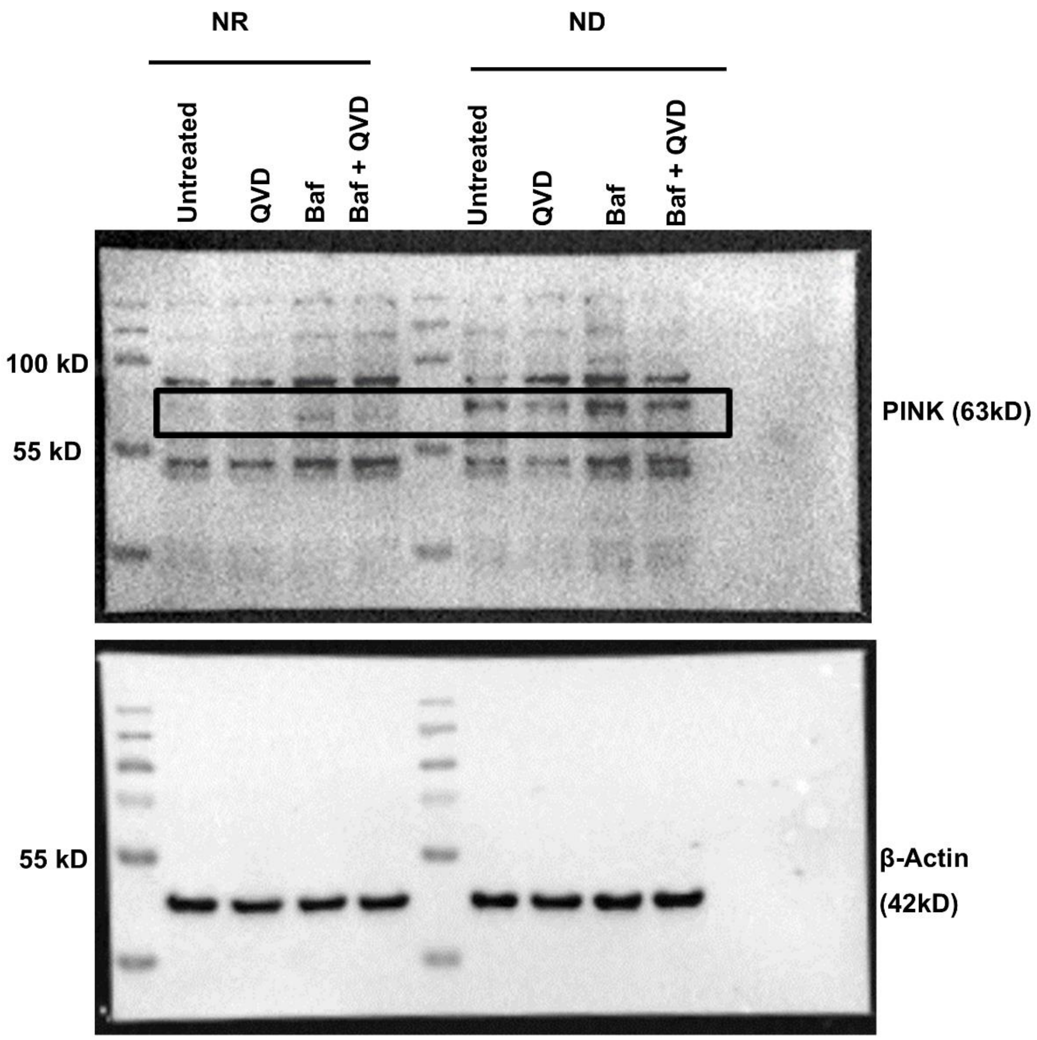


**Figure S29: Uncropped western blot images for Figures 2D, 3C and 3G.**

The same membrane was probed with the three antibodies after stripping. This is the reason why the same beta actin band is shown for all three images.
